# Identification of 370 genetic loci for age at first sex and birth linked to externalising behaviour

**DOI:** 10.1101/2020.05.06.081273

**Authors:** Melinda C. Mills, Felix C. Tropf, David M. Brazel, Natalie van Zuydam, Ahmad Vaez, eQTLGen Consortium, BIOS Consortium, Tune H. Pers, Harold Snieder, John R.B. Perry, Ken K. Ong, Marcel den Hoed, Nicola Barban, Felix R. Day, on behalf of the *Human Reproductive Behaviour Consortium*

## Abstract

Age at first sexual intercourse (AFS) and age at first birth (AFB) have implications for health and evolutionary fitness. In the largest genome-wide association study to date (AFS, N=387,338; AFB, N=542,901), we identify 370 independent signals, 11 sex-specific, with a 5-6% polygenic score (PGS) prediction. Heritability of AFB shifted from 9% [CI=4-14] for women born in 1940 to 22% [CI=19-25] in 1965. Signals are driven by the genetics of reproductive biology and externalising behaviour, with key genes related to follicle stimulating hormone (FSHB), implantation (ESR1), infertility, and spermatid differentiation. Polycystic Ovarian Syndrome leads to later AFB, linking with infertility. Late AFB is protective against later-life disease and associated with parental longevity. Higher childhood socioeconomic circumstances and those in the highest PGS decile (90%+) experience markedly later reproductive onset. Results are relevant for improving teenage and late-life health, for understanding longevity, and guiding experimentation into mechanisms of infertility.

---

The timing of onset of human reproductive behaviour – age at first sexual intercourse (AFS) and age at first birth (AFB) – has implications for reproductive health, adolescent development and evolutionary fitness. First sexual intercourse has occurred increasingly earlier, by the age of 16 for one-third of contemporary UK teenagers.^1^ Early reproductive onset is associated with teenage pregnancy^2^ but also adverse health outcomes such as cervical cancer, depression, sexually transmitted diseases^2^ and substance use disorders.^3,4^ In contrast to earlier sexual debut, we have witnessed progressively later ages at first birth for women, now reaching an average of 30 years in many modern societies, even later for men (Supp Info Fig S3).^5^ Late reproductive behaviour is associated with lower fecundity and subfertility^6^ and infertility traits such as endometriosis and early menopause,^7,8^ with over 20% of women born after 1970 in many modern countries now remaining childless.^9^ Earlier ages of sexual debut and later ages at first birth has marked the decoupling of reproduction from sexual behaviour in many contemporary societies, with implications for sexual, reproductive and later-life health (Supp Note Fig S2).

Since reproductive behaviour is shaped by biology and environment, a multidisciplinary approach is required to understand the common genetic aetiology and how it relates to health, reproductive biology, environment and externalising behaviour.^10^ Since the onset of reproductive behaviour generally occurs in adolescence to early adulthood, it is often linked to externalising behaviour such as self-control and psychiatric (e.g., ADHD) and substance use disorders (e.g., smoking, alcohol use), often moderated by the environment (e.g., childhood socioeconomic conditions).^10^ Furthermore, it may be that individuals inherit a common genetic liability for a spectrum of interlinked complex traits related to reproduction, health and longevity. There has been limited attention to understanding how these genetic effects are stratified by sex or across different socioeconomic and historical contexts.

In a previous GWAS of AFS (*n*=125,667)^11^ and AFB (*n*=343,072),^8^ we identified 38 and 10 novel independently-associated single-nucleotide polymorphisms (SNPs), respectively. The current study comprises a markedly expanded sample size for AFS (*N=387,338*) and AFB (*N*=542,901), uncovering 370 independent autosomal or X chromosomal loci, some of which are sex-specific, with 99 candidate genes expressed at the protein level in the brain, glands and reproductive organs. The multiple methods applied in this study (Fig S1) reveal underlying genetic drivers, common genetic liabilities, heterogeneity by childhood socioeconomic status and historical period and further evidence of the relationship of later reproductive onset with fewer later-life metabolic life diseases and increased longevity.

## Results

### Changes in reproductive behaviour and heritability over time

We first examine phenotypic changes in human reproductive behaviour and heritability over time. Descriptive analyses using the UK Biobank illustrate shifts in the mean AFS and AFB, changes in the shape of the distribution by birth cohort, and a bi-modal distribution of AFS in earlier cohorts (Fig 1A, Supp Note Fig S3). Whereas AFB was often in the early 20s for older birth cohorts, this distribution has spread and shifted to older ages over time, with a marked drop in Pearson’s correlation between AFS and AFB from those born <1941 (0.60) to those born >1960 (0.31) (Supp Note Fig S2). Using GREML,^12,13^ we found a steady increase in SNP-heritability by birth cohort for AFB for women from 9% [CI = 4-14] for those born in 1940, climbing to around 22% [19–25] for the latest cohorts born in 1965. For AFS, heritability ranges between 12 [7–18] for women born around 1940 and 23 [17–28] % for men born around 1940 with a trend for women similar to AFB and a suggestive U-shaped trend for men (Fig 1B for women, Supp Note Fig 4A-B, also for men).

**Figure 1.**
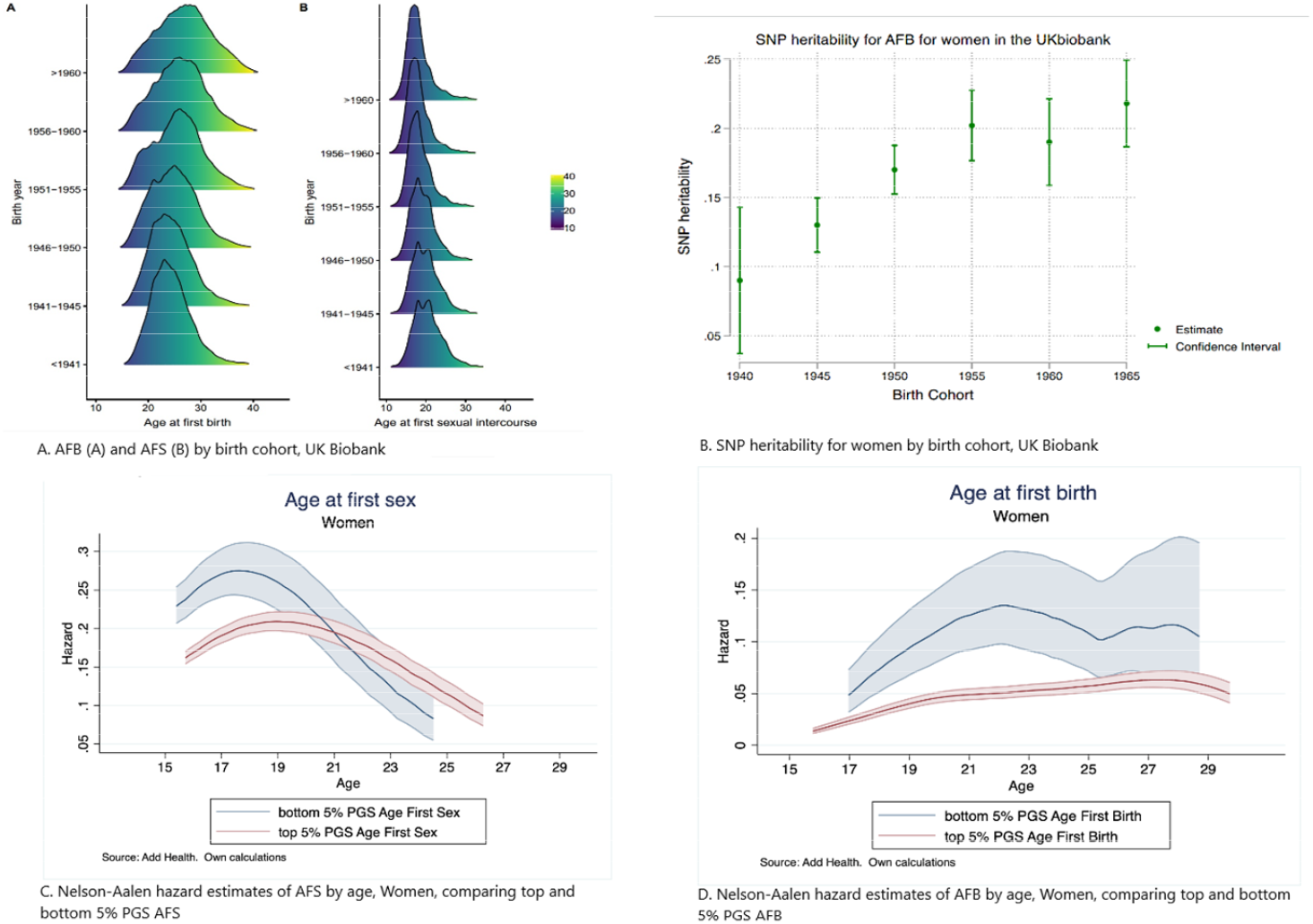
Age at first sex (AFS) and age at first birth (AFB) changes over time, heritability by birth cohort and polygenic score (PGS) prediction. (A) AFB (A) and AFS (B) by birth cohort, UK biobank shows a shift in the ages over time. (B) Increased SNP heritability for women over time by birth cohort, UK Biobank. (C) Nelson-Aalen hazard estimates of AFS by age for women comparing the upper and lower 5% of the PGS for AFS. (D) Nelson-Aalen hazard estimates of AFB by age for women comparing the upper and lower 5% of the PGS for AFB.

### Meta-analysis GWAS Human Reproductive Behaviour

We conducted a meta-analysis of GWAS results from 36 cohorts for AFS and AFB in individuals of European ancestry (defined by genetic principal components). We imputed to the 1000 Genomes Project reference panel in a pooled sample and then stratified the analysis by sex (Supp Note Tables S1-8). In total, we discovered 370 associated loci. The GWAS of AFS identified 281 (271 pooled of which 4 on the X chromosome; 2 women; 8 men) independent SNPs at genome-wide significance (*p*<5 × 10^-8^, Fig S5A-C; Table S10). The GWAS of AFB identified 89 (84 pooled of which 4 on the X chromosome; 1 women) independent SNPs at genome-wide significance (*p*<5 × 10^-8^, Fig S6A-C; Table S9). The distribution of genome-wide test statistics for AFS and AFB showed significant inflation (λ_GC_ = 1.84 and 1.47, respectively), however LD score regression showed that this could be attributed almost entirely to polygenicity rather than to population substructure (LD intercept AFS 1.07 (SE = 0.01); AFB 1.03 (SE = 0.01, Supp Note). The LD Score intercept test confirmed that only a very small percentage (5.5%) of the observed inflation in the mean *χ*^2^ statistic was due to population stratification or other confounders, rather than to a polygenic signal.

### Polygenic score prediction

We then calculated polygenic scores (PGSs) using three different specifications (Fig 3A, Supp Note, Sect 5). To validate the performance of the PGSs, we performed out-of-sample prediction in Add Health (a survey of adolescence to young adulthood in the US) and UKHLS (a representative survey of adults in the UK) cohorts using ordinary least-squares (OLS) regression models and report the R^2^ as a measure of goodness-of-fit of the model (Fig 3A, Supp Note 5). PGSs including all SNPs explain up to 5.8% of the variance for AFS and 4.8% for AFB. The difference between the out of sample prediction in the UKHLS and Add Health is related to heterogeneity between the initial GWAS sample, including a large UK older population from the UK Biobank, which is more comparable with the UKHLS population. Add Health participants also have higher levels of right-censoring (i.e., not yet experienced a birth). Previous research has demonstrated that meta-analyses of complex behavioural traits using populations from diverse national contexts and birth cohorts can be influenced by this hidden heritability.^13^ A 1 standard deviation (SD) change in the AFS/AFB PGS is associated with a 7.3 and 6.3 month delay in AFS and AFB, respectively. We then ran survival models to account for right-censoring, which occurs when an individual does not experience the event of first sex or birth by the time of the interview.^14^ Using Add Health data, we estimated nonparametric hazard functions and then compared individuals at the top and bottom 5% of the PGS (see Fig 1C AFS, 1D AFB for women, Supp Note Fig S8-9 men). Those in the top 5% PGS for AFS (i.e., genetic predisposition for later AFS) are less likely to have their sexual debut before age 19. AFS PGSs appear more relevant in explaining women’s AFS in comparison to men. Those in the top 5% PGS for AFB (i.e., genetic predisposition for later AFB) postpone AFB across all ages until approximately age 27, with similar curves for both sexes.

**Figure 2.**
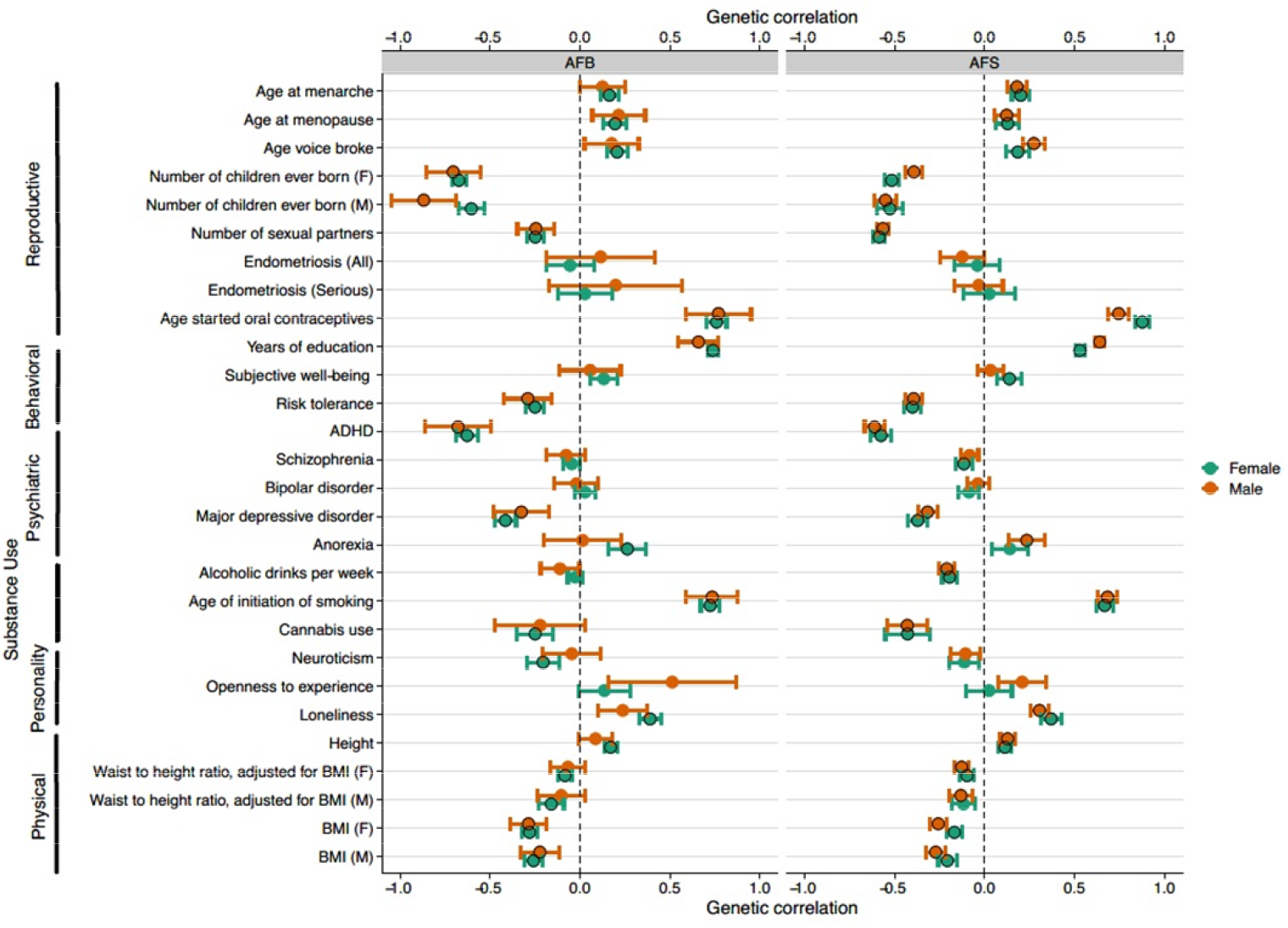
Genetic correlations of age at first birth (A) and age at first sex (B) with a selection of related traits. The horizontal bars represent 95% confidence intervals. If the trait was initially assessed separately for males and females, this is indicated on the left in brackets with F referring to females and M to males. The black circles represent the Bonferonni significant correlations but the magnitude of the effect (rg, the genetic correlation) is the most informative. Definitions and sources of all traits can be found in the Supplementary Note and full results are in Supplementary Table S13.

**Figure 3.**
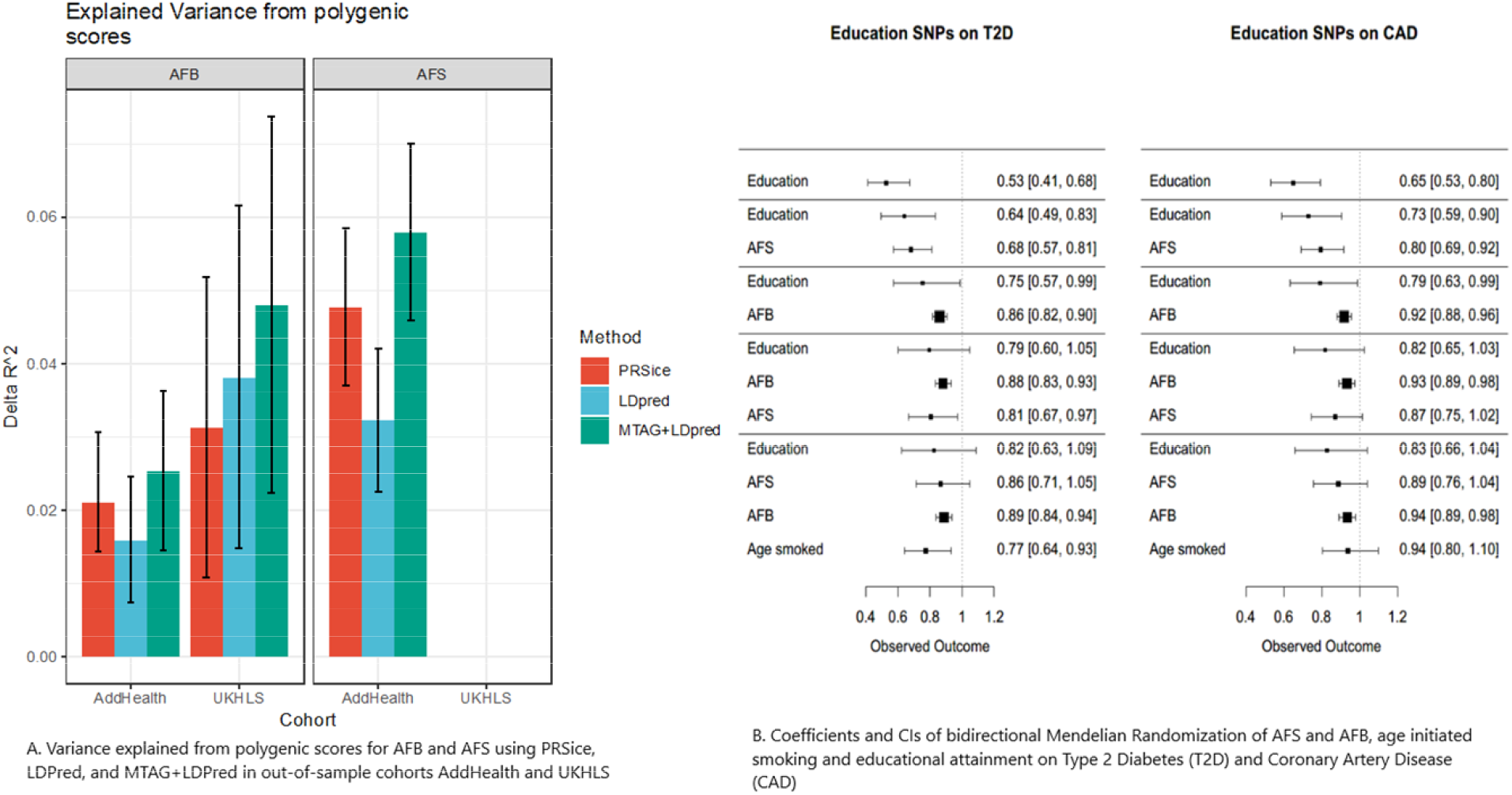
Variance explained from polygenic scores for age at first birth (AFB) and age at first sex (AFS) using various methods (A) and coefficients (CIs) of Bidirectional Mendelian Randomization of AFB and AFS, age initiated smoking and educational attainment on Type of Diabetes (T2D) and Coronary Artery Disease (CAD).

We next examined whether these genetic effects were environmentally moderated by childhood socioeconomic status. Disadvantaged socioeconomic status is highly related to early sexual behaviour and teenage pregnancy.^15^ To explore the impact of environmentally moderated parental genetic effects on our PGSs, we examined PGS prediction across low (0-10%), medium (50-60%) and high (90-100%) PGS percentiles by parents’ education (college versus no college) as a proxy for childhood socioeconomic status (Supp Note Fig 10A-B). Indeed, those in the highest decile of the PGS (90-100%) for later AFB have a higher AFB, particularly past age 27, which is accentuated for those with highly educated parents (Fig S10A). Being in the highest PGS decile for AFS is associated with later sexual intercourse (difference between highest and lowest decile is 2.08 years), especially for those from highest socioeconomic childhood households (2.39 years difference among high SES, compared to 1.62 years for low SES families) (Supp Note Fig 10B).

### Genetic correlations

To test the relationships of AFS and AFB with related phenotypes, we calculated genetic correlations using LD score regression^16^ (Fig 1, Supp Note Fig S11, S13, Tab S11). Given previous evidence,^8^ we examined 25 traits by sex from six relevant categories including: reproductive (e.g., age at menarche, number of sexual partners), infertility (e.g., endometriosis and severe endometriosis), behavioural (e.g., years of education, risk tolerance), psychiatric disorders (e.g., ADHD, schizophrenia), substance use (e.g., age of initiation of smoking, cannabis use), personality (e.g., openness to experience) and anthropometric (e.g., BMI, height). Logically, the strongest genetic correlations were observed for reproductive traits, particularly for number of children ever born. We remain cautious regarding the estimates of endometriosis considering the smaller sample size and larger confidence intervals (CIs) of our estimates. In our Mendelian Randomization analysis, discussed shortly, we explore the relationship with infertility in more detail. Behavioural traits also show strong genetic correlations, particularly with AFB and educational attainment in women (0.74, ±0.01), compared to AFS (0.53, ±0.01). There was also a negative genetic correlation between adult risk tolerance and AFS/AFB (AFS ~–0.40; AFB ~–0.25); i.e., those less genetically prone to risk are more genetically predisposed to postpone reproductive behaviour. Amongst psychiatric traits, often related to externalising behaviour, the strongest correlation was with ADHD (AFS females –0.58, ±0.03, males –0.61, ±0.03; AFB females –0.63, ±0.03; males – 0.68, ±0.09) and Major Depressive Disorder (MDD) (AFS females – 0.37, ±0.03, males –0.32, ±0.03; AFB females, –0.42, ±0.03; AFB males, –0.33, ±0.08). We also observed strong genetic correlations with age at onset of smoking (AFS ~ 0.68, ±0.03; AFB ~0.74, ±0.03), a trait that provides a unique window into adolescent substance use and externalising behaviour around the same time of early reproductive behaviour. Genetic factors influencing early smoking, early sexual debut and teenage pregnancy are thus – to some extent – shared. As shown in Fig 2, there are few sex differences in these correlations, with the exception of small variations in number of children, anorexia and openness to experience.

### Aetiology and causality

To explore aetiology and causality we employed GenomicSEM, Exploratory Factor Analysis and Bi-Directional Mendelian Randomization. To understand the relationships underlying these genetic correlations, we first used GenomicSEM.^17^ GenomicSEM uses structural equation modelling to decompose the genetic covariance matrix, calculated using multivariate LD score regression, of a set of traits. Parameters are estimated by minimising the difference between the observed genetic covariance matrix and the covariance matrix derived from the model (Supp Note, Sect 8). We fit a series of genetic regression models in which AFB (or AFS) was regressed on both years of education and one other possible mediating trait, such as openness, cognitive performance, ADHD and age of initiation of smoking (Supp Note Tab S12A-L, Fig S12A-B). In other words, we wanted to test whether the strong genetic correlation of AFS/AFB with education was the result of another mediating trait such as personality, ADHD or substance use. We found that the genetic correlation of years of education with AFB and AFS was independent of factors like risk tolerance, substance use, and psychiatric disorders. This suggests that the genetic correlation between years of education and AFB is largely a product of a strong bidirectional relationship between these traits, rather than being both downstream of a common identified cause. The exception was age at initiation of smoking – as noted previously, a window into risky adolescent behaviour – which partially mediated the relationship of AFB and AFS with years of education.

Exploratory factor analysis (EFA) was then used to examine whether the genetic signal of the onset of reproductive behaviour originated from two genetically distinguishable subclusters of reproductive biology versus externalising behaviour. Using a two-factor EFA model to fit the genetic covariance matrix AFS and AFB with these two additional traits, we found that the entire model accounted for 47% of the overall variance, with 22% attributed to risk tolerance and 4% to age at menarche. In a more robust analysis we fit a Genomic SEM for AFB in women and regressed several genetic measures of reproductive biology (age at menarche, age at menopause) and a latent factor representing a common genetic tendency for externalising behaviour (age at initiation of smoking, age first used oral contraception, ADHD) (Fig S14). These genetic factors predicted 88% of the variance, with the majority of variance significantly predicted by externalising factors (0.90,±0.02), followed by age at menopause (0.20, ±0.04) and age at menarche (0.16,±0.03). We note that selection bias, induced by the fact that AFB can only be measured among individuals with at least one live birth, may have potentially inflated this estimate.

Given the strong genetic correlations between the phenotypes discussed above, we used Mendelian Randomization (MR)-based analyses^18^ to explore causality and assess the direction of effect between AFB, AFS and years of education^19^ as well as risk taking (measured in adulthood)^4^ and age at smoking initiation^20^ (Supp Note Sect 9, Tab S13A). For the majority of pairs of phenotypes we found strong evidence of bi-directionality, which was also seen after applying Steiger fitting. The relationship between AFB and years in education appeared to be the explanatory factor that linked AFB to the two risk taking phenotypes. This was not the case, however, for AFS where the analysis suggests that age at initiation of smoking (and the environment and processes that lead to this) are upstream of the start of AFS. In that case the relationship was significant when assessed as age at smoking to AFS but not the other way round. Of note, associations were much stronger for age at smoking initiation than for risk-taking behaviour assessed in adulthood, suggesting that the timing of this behaviour is key.

A second set of MR analyses examined whether AFS and AFB PGSs have effects on type 2 diabetes (T2D)^21^ and coronary artery disease (CAD).^22^ Acknowledging the substantial overlap between the most significant signals for AFB, AFS and education related phenotypes, we used adjusted models to estimate effects independent of years of education and BMI (Fig 3B, Tab S13B, Fig S16). Given the extent of the overlap, we were also interested to investigate if AFB or AFS might attenuate any association particularly with education. T2D and CAD were chosen since they are two common major diseases, with broadly defined behavioural risk factors. Findings show that the association with years of education and later life diseases are substantially attenuated by the effects of AFB. We also found a strong association of the BMI weights with the AFS SNPs but note that even when BMI is included the model, the level of attenuation of the educational attainment results by AFB remains striking. This concurs with a large body of research that has established a biological association with the timing of AFB and metabolic diseases including early AFB linked to high blood pressure,^23^ obesity^24^ and diabetes.^25^ Reproductive timing thus appears to capture a latent variable that detects these metabolic effects and is a marker of a broader social trajectory that serves as a more powerful predictor of later life disease than years of education alone. This also suggests that many of the associations with diseases that have been previously ascribed to years of education, may result from this more broadly defined socio-behavioural trajectory captured by AFB.

Finally, since we were also interested in infertility-related phenotypes, bidirectional MR was performed with AFS and AFB with PCOS (polycystic ovarian syndrome) and given the nature of the disease, on women only.^26^ Our findings (Tab S13C) suggest that PCOS leads to later AFB. We find no effect of PCOS on AFS or of either AFS or AFB to PCOS, suggesting that the causal link is infertility-related with PCOS contributing to later AFB.

### Cox proportional hazard models of a polygenic score for AFB on longevity

The disposable soma theory of evolution hypothesises that longevity demands investments in somatic maintenance – such as remaining in education – that in turn reduces resources available for reproduction. To test trade-offs between reproductive behaviour and senescence as argued in the ageing and longevity literature,^27^ we conducted Cox proportional hazard models analyses to test whether our AFB PGS was associated with (parental) longevity (Supp Note, Tab S14). We first estimated a baseline Cox proportional hazard model of our AFB PGS on parental longevity and then included the PGS for educational attainment and risk covariates followed by a final model including number of siblings as a proxy for parental fertility. We found that a genetically predicted 1 SD increase in the PGS for AFB is associated with a 2-4% reduction in parental mortality at any age, suggesting that there is likely a trade-off between the timing of reproduction and longevity.

### Gene prioritization

To understand the biology represented by the variants associated with AFS and/or AFB, we performed a gene prioritization analysis that connected variants to genes and prioritized candidate genes based on likely involvement in reproductive biology or psychiatric traits. To this end, we used predicted gene function,^28^ single-cell RNA sequencing data in mice,^29,30^ literature mining,^31^ *in silico* sequencing,^32^ and Summary-data based Mendelian Randomization(SMR)^33^ using eQTL data from brain and blood.^34^ Integrating results across all approaches resulted in the prioritization of 386 unique genes; 314 genes in 159 loci for AFS and 106 genes in 42 loci for AFB (Supp Tab 15A-19C). Of these, 99 were expressed at the protein level in cell types of brain, glands, and/or (fe)male reproductive organs^35^ (Fig 4). Gene prioritization in sex-specific loci resulted in the prioritization of 11 genes for AFB in women, one gene for AFS in women and 23 genes for AFS in men. Of these, 12 genes at three loci were expressed at the protein level in relevant tissues (Supp Note, Fig S17).

**Figure 4.**
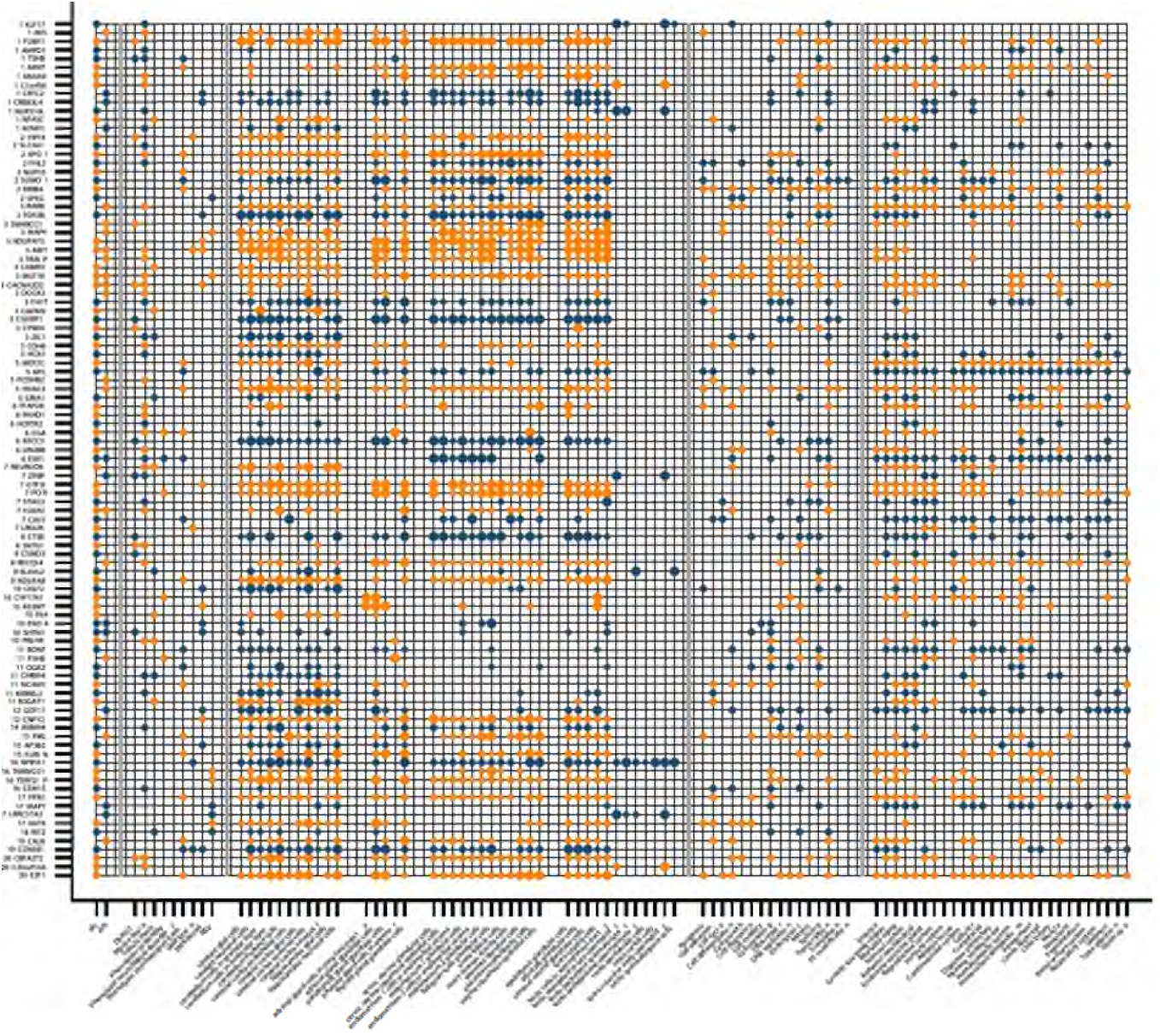
Gene prioritization of age at first sex (AFS) and age at first birth (AFB) Information for 99 genes prioritized in loci identified by GWAS for age at first sex and/or age at first birth that are located within 1 million bp of lead SNPs and are expressed at the protein level in brain, glands and/or reproductive organs. Transitions in colour from blue to orange highlight whether the gene in the next row is still within the same locus or not. Numbers before the genes show the chr. Panels are separated by vertical grey lines. The first panel (left) indicates if the locus was identified as being associated at genome-wide significance with age at first sex (AFS) and/or age at first birth (AFB). The second panel shows which bioinformatic approaches highlighted the gene as a candidate. The third panel shows – from left to right-the cell types in brain, glands, female reproductive organs, and male reproductive organs in which the genes are expressed at a low, moderate or high level (small, medium and large circles), based on data from the Human Protein Atlas. The fourth panel shows gene functions as extracted from Entrez, Uniprot and GeneCards. The fifth panel indicates which phenotypes were observed in mutant mice, as reported by the Mouse Genome Informatics (MGI) database.

Genes that play a role in follicle stimulating hormone (*CGA* ^36^), oocyte development (*KLF17*^37^), and implantation and placental growth (*ESR1*, *SUMO1*^38^, *ARNT*,^39^ *CAV1*,^40^ *E2F1*^41^) were prioritized for AFS in data from men and women combined, while *FSHB*^42^ and *ESR1* were (also) prioritized for AFB. Other genes prioritized in loci identified in the pooled meta-analyses were expressed at the protein level in (developing) sperm – highlighting a role for spermatid differentiation (*KLF17*^43^) for AFS – as well as for sperm morphogenesis and binding between acrosome-reacted sperm and the zona pellucida (*ZPBP*^44^) for AFB. The meta-analysis in data from only women yielded genes related to endometriosis (*CCR1*)^45^ and spontaneous abortion (*CXCR6*) for AFB (Supp Note Fig S19).^46^ Taken together, these results suggest that intrinsic biological processes that influence fertility also influence the onset of sexual behaviour in men and women. Interestingly, *NUP210L* – prioritized for AFS and highly expressed in developing and mature sperm^35^ – is normally testis-specific, but was recently shown to be expressed in prefrontal cortex neurons of G allele carriers in rs114697636 (MAF 3%, D’ 0.90 with AFS lead SNP rs113142203), attributed to allele-specific activation through improved binding affinity for testis receptor 2.^47^ Methylation of, and variants near *NUP210L* have been associated with psychologic development disorders, intelligence, and mathematical ability,^48^ illustrating how a testis-specific gene could be linked to brain function in some individuals. We note, however, that the fact that some prioritized proteins are expressed in some relevant tissues does not provide clear evidence supporting a causal role for the prioritized genes.

Several genes prioritized in AFS-associated loci in data from men and women combined have previously been implicated in risk seeking behaviour, sociability and anxiety (*GTF2I*,^49^ *TOP2B*,^50^ *E2F1*^51^, *NCAM1*,^52^ *NFASC*,^53^ *MEF2C*^54^). In the sex-specific meta-analysis for AFS, a role for externalising behaviour was supported through *ERBB4* in women; and through *SLC44A1* and *NR1H3* in men. *ERBB4* has previously been linked to fear, anxiety,^55^ schizophrenia,^56^ and polycystic ovary syndrome (PCOS);^57^ *SLC44AI* encodes a choline transporter that plays a key role in cerebral inhibition related to substance use and depressive disorders^58^; and *NR1H3* has been implicated in major depressive disorder (MDD).^59^ These genes provide concrete examples of how an innate predisposition for externalising behaviour may influence initiation of reproductive behaviour.

The gene prioritization results partly mirror and compliment the rigorous post-GWAS *in silico* association analyses we performed for loci identified for AFS and AFB. However, experimental validation is required before firm conclusions can be drawn about the involvement of, and mechanisms through which prioritized candidate genes influence AFS and AFB. More information on protein-protein interaction hubs, as well as on genes highlighted by literature mining^31^ are provided in the Supplementary Note.

## Discussion

In this study, we presented the results of the largest GWAS to date of the onset of human reproductive behaviour of age at first sex (AFS) (N=387,338) and age at first birth (AFB) (N=542,901). We identified 370 independent signals harbouring at least 386 prioritized candidate genes, using 1000G imputed genotype data and an X-Chromosome analysis, which allowed us to detect considerably more signals than ever before (Fig 5). In comparison, a recent GWAS for type 2 diabetes,^60^ for instance, detected 243 loci. Similar to previous work, we showed that the total SNP heritability accounted for 10-22% of phenotypic variance and varied by birth cohort.^13,61^ The incremental R^2^ of our PGSs based on significantly associated loci is around 5-6%, similar to direct effects observed for commonly used demographic and social variables (e.g., years of education, age at marriage), classically used to explain the timing of human reproductive behaviour by social scientists. Comparatively, a PGS of 5-6% is in the range observed for other complex traits, like BMI (5.8%)^62^ and schizophrenia (8.4%).^63^ The number of signals also opened up opportunities for functional follow-up analyses which suggested a role for spermatid differentiation and oocyte development. The analyses of the correlation and underlying aetiology of these traits revealed a common genetic basis of both AFS and AFB with externalising behaviour and substance use and links to infertility.

**Figure 5.**
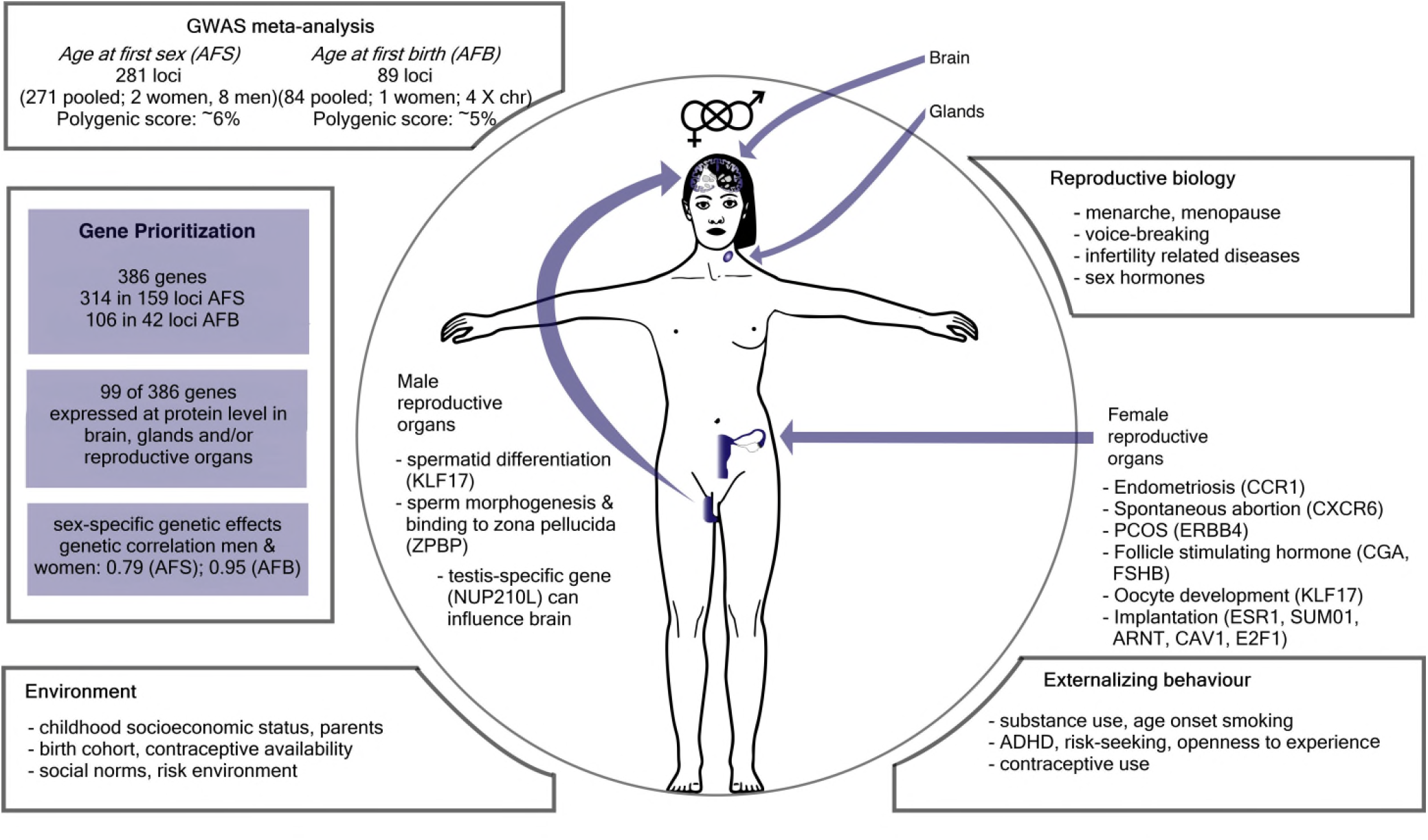
Summary genome-wide association study of timing of onset of reproductive behaviour: age at first sex (AFS) and age at first birth (AFB).

Finally, we showed that AFB is an important predictor for late age at onset of disease and longevity, and that it substantially attenuates the effect of years in education. We note that there are some situations where we have a significant Eggers intercept in the MR analysis, including some for the bidirectional data. Here there is likely to some heterogeneity in the data (AFB to education, and AFS to risk), where there are likely important pleotropic effects. However, this does not impinge on our central finding that there is widespread bi-directionality. Since we also find a significant intercept for AFB to CAD, and since in the adjusted model there are not significant effects, we are confident that we are not at risk of a false positive.

Although we opened many new avenues for research, the present GWAS still faces certain limitations. First, the sample sizes for men were appreciably smaller than for women since reproductive and fertility data is routinely collected less often from men. Yet to understand the causes of infertility in men this information needs to be taken into consideration in future data collection. The paucity of sex differences in the genetic correlations we observe between AFB, AFS, and a variety of related traits, including sex-specific traits such as age at menarche, suggests that the relevant processes overlap between the sexes. Initial within-family analyses showed that our discovery GWAS may actually overestimate causal effects (Supp Note); genotypes associated with later onset of reproductive behaviour genotypes are also associated with parental reproductive genotypes, likely leading to a social environment that affects reproductive and other behaviours. Collection and analysis of family data is clearly a future area of research for reproductive and related complex behaviour. The lack of accessibility of publically available summary statistics from some published research, meant that we were unable to examine the relationship with other traits, particularly with infertility related traits (e.g., larger studies of endometriosis). Future data collection could benefit from focussing on externalising and behavioural disinhibition markers that appear to be highly related to self-control, which has implications for disease prevention and behavioural interventions into lifestyle factors related to obesity, Type 2 diabetes or substance use disorders. A final glaring limitation is our focus on European-ancestry individuals in Western countries. Whilst common in this area of research,^64^ extension to other ancestries and geographical contexts is required in the future. This is particularly relevant in the context of parent gene-environment interactions, which may be specific to the social and environmental makeup of the sample.

Our detailed correlation, GenomicSEM and MR analyses also provided a deeper understanding of the underlying aetiology of related traits and pleiotropy and the associations between human reproductive behaviour and disease risk. We anticipate that our results will provide leads to address important interventions in infertility, teenage sexual and mental health, as well as for functional follow-up experiments that will likely yield targets that can be translated in efficient medication to improve fertility (e.g., in IVF) but also for interventions on reproductive health related to earlier sexual debut and teenage pregnancy.

## Methods

This article has a Supplementary Note with more details.

### Samples

For Age at First Sexual intercourse (AFS), we included 397,338 pooled individuals (n=182,791 males; n=214,547 females) from the UK Biobank. For Age at First Birth (AFB), we included 542,901 individuals (n=124,088 males; n=418,758 females) from 36 studies. We performed a GWAS separately restricted to European ancestry individuals that passed quality control. European ancestry was chosen in this discovery study due to the availability of samples^64^ and for no biological or substantive reason. We acknowledge that social science research has found large differences in the earlier initiation of AFS and AFB by lower socioeconomic status, which often coincides with societal inequality^65,66^ and the socially (not biologically) constructed categories of race and ethnicity. Socioeconomic differences are examined in this article, but results are only applicable to European Ancestry groups, with a need for further cross-ancestry discovery research.

The *Human Reproductive Behaviour Consortium*. This consortium is a collaboration studying the GWAS of human reproductive behaviour including age at first sex and birth, number of children ever born, childlessness and related traits. In some cases we used summary statistics from our first GWAS of AFB and NEB^8^ on discovery cohorts (see Supp Note Tables S1-S3b).

### Phenotype measurements, genotyping, imputation and meta-analysis

AFS is treated as a continuous measure with individuals considered as eligible if they had given a valid answer and ages lower than 12 excluded (see Supp Note 1.2). Since AFS has a non-normal distribution, a within-sex inverse rank normal transformation is required. AFB is also treated as a continuous measure, assessed for those who have ever given birth to a child. Details about participating cohorts, sample inclusion criteria, genotyping and imputation, models used to test for association, X chromosome analysis, quality control filters and diagnostics, and meta-analysis are in the Supp Note. A sample-size weighted meta-analysis of quality-controlled cohort-level results was performed using the METAL software.^67^ We performed conditional and joint multiple SNP analyses (COJO) to identify further independent SNPs and sex-specific analyses.

### Sex-specific genetic effects

We used LD score bivariate regression^68^ to estimate the genetic correlation between men and women based on the sex-specific summary statistics from the meta-analysis results. There was a large genetic overlap among the sexes for AFB (0.95) and a somewhat lower overlap for AFS (0.79), suggesting sex-specific effects would be important to examine. In order to determine if there was evidence for sex-specific effects, we compared the allelic effects for these SNPs between men and women and derived a *p*-value for heterogeneity.^69^ A multiple testing correction was applied (0.05/242=2 × 10^-4^) to identify sex-specific associations. We then selected a region of ±1Mb around these lead SNPs to identify the genes that may be represented by these lead SNPs, followed by gene prioritization as we did for the main AFB and AFS analyses.

### X chromosome analysis

For AFS, the UK Biobank provided results for between 977,536 and 990,735 variants on the X chromosome after QC (see Table S8). For AFB, 13 cohorts provided information on the X chromosome. Overall, we received 23 files, 13 for women, 8 for men and 2 for the pooled analysis in case there were individuals who were relatives in the data. On average, 275,023 variants survived QC with a minimum of 99,794 in women from WLS to 998,304 for the women in the UK Biobank sample (see TableS7 for full descriptives). Association analyses on the X chromosome were performed using software suggested in the analysis plan (XWAS, SNPtest or BOLT-LMM) using BOLT-LMM for AFS as this was only assessed in the UK Biobank data, for AFB, METAL was used as described above (see sup. note 3.5)

### Phenotypic and genotypic historical changes

Descriptive analyses and correlations were undertaken using the UK Biobank data to illustrate phenotypic shifts in the age of AFS and AFB by birth cohort, in addition to changes in the spread of the distribution. Pearson’s correlation coefficients were calculated and correlation graphs illustrate the changing relationship between the two phenotypes over time. Genotypic changes and SNP-heritability by birth cohort were quantified in UK Biobank data using GREML^12^ as described earlier.^13^

### MTAG: Multi-trait analysis of GWAS

MTAG results^70^ were calculated using GWA meta-analysis results of the following related phenotypes: AFS, AFB, number of children ever born, childlessness. Using summary statistics from the pooled GWAS of each of the traits, MTAG uses bivariate LD score regression to account for unobserved sample overlap.

### Polygenic score prediction

We performed out-of-sample prediction in two cohorts, the National Longitudinal Study of Adolescence to Adult Health (Add Health),^71^ based in the US and the UK Household Longitudinal Study-Understanding Society (UKHLS).^72^ We calculated three sets of polygenic risk scores (PGS) with weights based on meta-analysis results excluding the specific cohort from the calculation. First, pruning and thresholding of all SNPs was performed (250kb window; r^2^=.1) using PRSice^73^. Second, LDpred PGSs^74^ with the LD reference were calculated from the same genotyped files, using prior distributions for the causal fraction of SNPs equal to 1 and LDpred weights calculated under the infinitesimal model. Third, MTAG + LDpred PGSs were calculated using the same methodology as in the second PGSs, but this time based on MTAG results^70^. For both traits, we ran ordinary least-squares (OLS) regression models and report the incremental R^2^ as a measure of goodness-of-fit of the model. Confidence intervals are based on 1,000 bootstrapped samples.

### Testing population stratification, survival models and environmentally moderated parental genetic effects of childhood socioeconomic status

To test whether population stratification biased our results or lead to false positives, we used the LD Score intercept method.^75^ For each phenotype, we used the “eur_w_ld_chr” files of LD Scores.^16^ These LD Scores were computed with genotypes from the European-ancestry samples in the 1000 Genomes Project using only HapMap3 SNPs with MAF > 0.01. We then ran survival models to account for right-censoring, which occurs when an individual does not experience the event of first sex or birth by the time of the interview.^14^ Using Add Health data, we estimated nonparametric hazard functions based on Nelson-Aalen estimates and then compared individuals at the top and bottom 5% of the PGS and plotted the estimated hazards. To further explore the impact of environmentally moderated parental genetic effects on our PGSs, we examined PGS prediction across low (0-10%), medium (50-60%) and high (90-100%) PGS percentiles by parent’s education status (college versus no college), which serves as a proxy for childhood socioeconomic status.

### Genetic correlations

Genetic correlation (r_g_) values were computed to estimate the genetic correlation between the two traits using all polygenic effects captured by the SNPs and LD-score regression.^76^ We used summary statistics and the 1000 Genomes reference set, and restricted the analysis to European populations. We also followed the common convention of restricting our analyses to SNPs with MAF >0.01, thus ensuring that all analyses were performed using a set of SNPs that were imputed with reasonable accuracy across all cohorts. The standard errors (SEs) were produced by the LDSC python software package that uses a block jackknife over the SNPs. We estimated the genetic correlation between 28 different traits, pooled by both sexes and then divided by sex. Traits were divided into the six categories of: reproductive, behavioural, psychiatric disorders, substance use disorders, personality and anthropometric.

### Genomic SEM (structural equation modelling) and Exploratory Factor Analysis (EFA)

In an attempt to understand the aetiology of the correlations, we used the R package GenomicSEM to fit genetic multivariable regression models. GenomicSEM^17^ uses structural equation modelling to decompose the genetic covariance matrix, calculated using multivariate LD score regression, of a set of traits. Formally, structural equation models subsume many statistical methods and are quite flexible. We fit a series of genetic multivariable regression models, in which AFB was regressed on EA (educational attainment) and a trait X, in which we modelled various relevant traits such as openness, cognitive performance and AI (age initiation smoking). We also fit an analogous series of models in which AFS was regressed on EA.

Exploratory factor analysis (EFA) and Genomic SEM by reproductive biology and externalising behaviour. EFA was used to examine whether the genetic signal of the onset of reproductive behaviour originated from two genetically distinguishable sub-clusters of a biological component and an externalising behaviour component. This would suggest distinct causal mechanisms and subtypes of individuals. We tested this by fitting a two factor EFA model to the genetic covariance matrix of AFB, AFS, NEB, and the proxies age at menarche (biological component) and risk tolerance (externalising behaviour). To test this further, we estimated a more robust and additional measures of reproductive biology and externalising behaviour and a sex-specific analysis of AFB for women. We fit a genomic structural equation model (Genomic SEM) where AFB in women is regressed on age at menopause, age at menarche, and a latent factor representing the common genetic tendency to externalising behaviour. The factor is measured by AFS in women, age at initiation of smoking, age first used oral contraception, and ADHD, with the model scaled to unit variance for the latent factor.

### Bi-directional Mendelian Randomization

We then tested whether causal pathways linking these phenotypes are potentially bidirectional and whether our phenotypes might offer distinct contributions. We identified 1000 Genomes proxies for our SNPs and used these in multivariate Mendelian Randomisation (MR) models. All data was assumed to be on the forward strand, and as many of these data sets included UK biobank, allele identifier were matched to this study as a reference. First, we modelled the interplay between AFB, AFS and EA (educational attainment)^77^ as well as risk taking (measured in adulthood)^4^ and age at smoking initiation (AI).^20^ In each case IVW^78^ and MR-EGGER^79^ methods were performed, with an additional round of IVW performed once a Steiger filter^80^ had been applied to remove SNPs that appears to show a primary association with the outcome rather than the exposure. Multivariate MR was use to try to dissect causal pathways.^81^ A second set of MR analyses focused on links to late life diseases, namely type 2 diabetes (T2D)^82^ and coronary artery disease (CAD)^22^, using the same methods. Acknowledging the substantial overlap between the most significant signals for AFB, AFS and education related phenotypes, we use multivariate methods to test whether AFS or AFB had independent effects once the well-established links to length of educational attainment, and BMI were controlled for. Finally, as there was a particular interest in infertility related phenotypes, bidirectional MR was performed with AFS and AFB and PCOS (polycystic ovarian syndrome). In this case, given the sex-specific nature of the disease, a specific analysis was also performed on women only.

### Cox proportional hazard models of AFB polygenic score on longevity

To test trade-offs between reproductive behaviour and senescence, we conducted additional analyses to test whether our PGS for AFB was predictive of (parental) longevity. We restricted our models to mortality after age 60 to limit the possibility that early mortality affects parental fertility (i.e., collider bias).^83^ We calculated PGSs for AFB, Educational attainment (EA)^84^ and risky behaviour^4^ from the UK Biobank adopting the following procedure. We first split the sample in 10 random groups. We then iteratively estimated genome-wide association results for 9/10^th^ of the sample and used these association results as weights for the calculation of polygenic scores in the remaining 1/10^th^ of the sample. Polygenic scores were calculated using PRSice on a set of independent genotyped SNPs. We then estimated three sets of Cox Proportional hazard models to estimate the effect of the PGS of AFB on maternal and paternal age at death. All models control for the first 10 Genetic Principal Components, sex and year of birth, and are stratified by Local Authority District at birth calculated using the geo-coordinates provided in the UK Biobank due to differences in mortality related to material deprivation.^85^ We first estimated a baseline model and then included PGSs for EA and risk as covariates, followed by a final model including number of sibling (proxy for parental fertility).

### Gene prioritization

We prioritized candidate genes in pooled and sex-specific GWAS-identified loci using predicted gene functions,^86^ single-cell RNA sequencing data in mice,^29,87,88^ literature mining,^31^ *in silico* sequencing,^32^ and synthetic Mendelian Randomization^89^ using eQTL data from brain and blood.^90,91^

DEPICT and CELLECT for tissue, cell type and gene prioritization were used. First, DEPICT was used to perform pathway analyses, identify enrichment for cell types and tissues, and prioritize candidate genes.^86^ DEPICT is agnostic to the outcomes analyzed in the GWAS and employs predicted gene functions. For both AFS and AFB, all SNPs with *p*<1×10^-5^ in the pooled GWAS meta-analysis were used as input. Based on the results of the tissue enrichment analysis, we used CELLECT^88^ to identify nervous system cell types that are enriched for expression of genes in loci reaching *p*<1×10^-5^ in the GWAS, using RNAseq data from mouse brain.^29^ A similar approach using tabula muris RNAseq data^87^ helped prioritize additional central nervous system and pancreatic cell types for AFS. For enriched cell types from mouse brain and tabula muris, the top-10 contributing genes were selected as candidate genes resulting in the prioritization of 296 genes for AFS and 95 for AFB based on mouse brain; and 97 genes for AFS based on tabula muris data.

Phenolyzer to integrate prior knowledge and phenotype information. We used Phenolyzer (v1.1) to prioritize candidate genes by integrating prior knowledge and phenotype information.^92^ Here we used the regions defined by DEPICT v1.1, reflecting loci reaching *P*<1×10^-5^ in first instance. Phenolyzer takes free text input and interprets these as disease names by using a word cloud to identify synonyms. It then queries precompiled databases for the disease names to find and score relevant seed genes. The seed genes are subsequently expanded to include related (predicted) genes based on several types of relationships, e.g., protein-protein interactions, transcriptional regulation and biological pathways. Phenolyzer uses machine-learning techniques on seed genes and predicted gene rankings to produce an integrated score for each gene. We used search terms capturing three broad areas, i.e., (in)fertility, congenital neurological disorders and psychological traits, based on results from pathway, tissue and cell type enrichment analyses.

*In silico* sequencing. We used *in silico* sequencing to identify non-synonymous variants with an R^2^ for LD>0.7 with the lead SNPs in AFS and AFB-associated loci,^32^ which yielded genes that may drive the GWAS associations through direct effects on protein function.

Summary data-based Mendelian Randomization (SMR) and Heterogeneity in Dependent Instruments (HEIDI).^89^ We conducted this using eQTL data from brain^93^ and whole blood.^91^ This approach provided a list of genes that showed Bonferroni corrected significant evidence (thresholds for blood <3.2×10^-6^ brain <6.7×10^-6^) of mediating the association between our phenotypes and GWAS-identified loci based on results from brain and blood.

Integration of findings across all functional approaches. We integrated findings across all approaches and retained genes in loci that reached genome-wide significance, and that were located within 1M bp of a GWAS lead SNP. We next used data from the Human Protein Atlas^35^ to identify genes amongst 387 genes that are expressed at a low, medium or high protein level in brain, glands, and/or reproductive organs at a ‘supported’ or ‘enhanced’ degree of reliability. For the 97 genes that fulfilled these criteria, we mapped the brain, glandular and reproductive cell types in which they are highly expressed at the protein level;^94^ used a text-mining approach to extract functions from entries in Entrez, GeneCards and Uniprot; and identified phenotypes in mutant mice from the Mouse Genome Informatics (MGI) database^95^.

## Data availability

Our policy is to make genome-wide summary statistics widely and publically available. Upon publication, summary statistics will be available on the GWAS Catalog website: https://www.ebi.ac.uk/gwas/downloads/summary-statistics The phenotype and genotype data for separate studies used in this GWAS are available upon application to each of the participating cohorts who can be contacted directly to follow their different data access policies. Access to the UK Biobank is available through application with information available at: http://www.ukbiobank.ac.uk).

## Code availability

No custom code was used with all analyses and modelling using standard software as described in the Methods section and in detail in the Supplementary Information.

## Acknowledgements

A detailed list of acknowledgments for each cohort can be found in Section 12.2 of the Supplementary Note. The research leading to these results has received funding from PI M.C.Mills from the European Research Council (ERC) Consolidator Grant SOCIOGENOME (615603, www.sociogenome.org), ERC Advanced Grant CHRONO (835079), Economic & Social Research Council (ESRC) UK, National Centre for Research Methods (NCRM) grant SOCGEN (ES/N011856/1), Wellcome Trust ISSF and a large Centre grant from the Leverhulme Trust, Leverhulme Centre for Demographic Science. The funders had no role in study design, data collection and analysis, decision to publish or preparation of the manuscript. We thank Evelina T. Akimova and Stine Møllegaard for administrative work in the organization of the cohort information and author list.

## Author contributions

MCM and FRD designed and led the study. MCM wrote the paper and supplementary note with contributions by authors for respective analyses and comments by all authors. DMB conducted phenotypic changes, phenotype preparation, LD Score and genetic correlations, Genomic SEM and exploratory factor analysis and sex-specific effects. NB conducted GWAS meta-analysis, MTAG, PGS prediction, survival models, and Cox models of longevity. FCT and FRD conducted the cohort QC. FCT conducted GREML cohort heritability analysis and phenotype preparation in UKBB. FRD ran Mendelian Randomization, conducted GWAS analyses and JRBP conducted COJO and X-Chromosome analysis. NvZ conducted DEPICT and Phenolyzer analyses. AV and HS conducted in silico sequencing and SMR analyses. TP conducted cell type enrichment analyses. MdH integrated gene prioritization results and performed downstream analyses, e.g. Human Protein Atlas; Entrez, GeneCards and Uniprot mining; and STRING Protein-Protein interaction analyses. Authors in the Human Reproductive Behaviour Consortium conducted cohort specific GWAS and other analyses, and contributed through the administration, management, and data collection for the participating cohorts. The eQTLGen and BIOS Consortiums provided data for additional analyses. All authors reviewed and approved the final version of the paper.

## Competing interests

The main authors declare no competing interests. The views expressed in this article are those of the author(s) and not necessarily those of the NHS, the NIHR, or the Department of Health. MMcC (Mark McCarthy) has served on advisory panels for Pfizer, NovoNordisk and Zoe Global, has received honoraria from Merck, Pfizer, Novo Nordisk and Eli Lilly, and research funding from Abbvie, Astra Zeneca, Boehringer Ingelheim, Eli Lilly, Janssen, Merck, NovoNordisk, Pfizer, Roche, Sanofi Aventis, Servier, and Takeda. As of June 2019, MMcC is an employee of Genentech, and a holder of Roche stock.

## Corresponding authors

Melinda C. Mills (https://orcid.org/0000-0003-1704-0001), and Felix R. Day (https://orcid.org/0000-0003-3789-7651)

## Author information

These authors contributed equally: Melinda C. Mills, Felix C. Tropf, David M. Brazel, Felix R. Day, Nicola Barban, Marcel den Hoed

These authors designed and led the study: Melinda C. Mills, Felix R. Day

### Affiliations

**Leverhulme Centre for Demographic Science, University of Oxford & Nuffield College, UK**

Melinda C. Mills, Felix C. Tropf, David M. Brazel

**MRC Epidemiology Unit, Institute of Metabolic Science, University of Cambridge, Cambridge, United Kingdom**

Felix R. Day, John R.B. Perry, Ken K. Ong

**Department of Statistics, University of Bologna, Bologna, Italy**

Nicola Barban

**École Nationale de la Statistique et de L’administration Économique (ENSAE) & Center for Research in Economics and Statistics (CREST), Paris, France**

Felix C. Tropf

**Department of Epidemiology, University of Groningen, University Medical Center Groningen, Groningen, The Netherlands**

Harold Snieder, Ahmad Vaez

**The Beijer Laboratory and Department of Immunology, Genetics and Pathology, Uppsala University and SciLifeLab, Uppsala, Sweden**

Marcel den Hoed, Natalie van Zuydam

**The Novo Nordisk Foundation Center for Basic Metabolic Research, Faculty of Health and Medical Sciences, University of Copenhagen, Copenhagen, Denmark**

Tũne H. Pers

**Department of Bioinformatics, Isfahan University of Medical Sciences, Isfahan, Iran**

Ahmad Vaez

## Consortia

### Human Reproductive Behaviour Consortium

Author list ordered alphabetically

Evelina T. Akimova, Sven Bergmann, Jason D. Boardman, Dorret I. Boomsma, Marco Brumat, Julie E. Buring, David Cesarini, Daniel I. Chasman, Jorge E. Chavarro, Massimiliano Cocca, Maria Pina Concas, George Davey-Smith, Gail Davies, Ian J. Deary, Tõnu Esko, Oscar Franco, Audrey J. Gaskins, Eco J.C. de Geus, Christian Gieger, Giorgia Girotto, Hans Jörgen Grabe, Erica P. Gunderson, Kathleen Mullan Harris, Fernando P. Hartwig, Chunyan He, Diana van Heemst, W. David Hill, Georg Homuth, Bernando Lessa Horta, Jouke Jan Hottenga, Hongyang Huang, Elina Hyppönen, M. Arfan Ikram, Rick Jansen, Magnus Johannesson, Zoha Kamali, Maryam Kavousi, Peter Kraft, Brigitte Kühnel, Claudia Langenberg, Lifelines Cohort Study, Penelope A. Lind, Jian’an Luan, Reedik Mägi, Patrik K.E. Magnusson, Anubha Mahajan, Nicholas G. Martin, Hamdi Mbarek, Mark I. McCarthy, George McMahon, Matthew B. McQueen, Sarah E. Medland, Thomas Meitinger, Andres Metspalu, Evelin Mihailov, Lili Milani, Stacey A. Missmer, Stine Møllegaard, Dennis O. Mook-Kanamori, Anna Morgan, Peter J. van der Most, Renée de Mutsert, Matthias Nauck, Ilja M. Nolte, Raymond Noordam, Brenda W.J.H. Penninx, Annette Peters, Chris Power, Paul Redmond, Janet W. Rich-Edwards, Paul M. Ridker, Cornelius A. Rietveld, Susan M. Ring, Lynda M. Rose, Rico Rueedi, Kári Stefánsson, Doris Stöckl, Konstantin Strauch, Morris A. Swertz, Alexander Teumer, Gudmar Thorleifsson, Unnur Thorsteinsdottir, A. Roy Thurik, Nicholas J. Timpson, Constance Turman, André G. Uitterlinden, Melanie Waldenberger, Nicholas J. Wareham, Gonneke Willemsen, and Jing Hau Zhao

### Affiliations

Author list ordered alphabetically

**Leverhulme Centre for Demographic Science, University of Oxford, Oxford, United Kingdom**

Evelina T. Akimova

**Department of Computational Biology, University of Lausanne, Lausanne, Switzerland**

Sven Bergmann, Rico Rueedi

**Swiss Institute of Bioinformatics, Lausanne, Switzerland**

Sven Bergmann, Rico Rueedi

**Department of Integrative Biomedical Sciences, University of Cape Town, Cape Town, South Africa**

Sven Bergmann

**Department of Sociology and Institute of Behavioral Science, University of Colorado at Boulder, Boulder, CO, United States of America**

Jason D. Boardman

**Department of Biological Psychology, Amsterdam Public Health Research Institute, Vrije Universiteit Amsterdam, Amsterdam, The Netherlands**

Dorret I. Boomsma, Eco J.C. de Geus, Jouke Jan Hottenga, Hamdi Mbarek, Gonneke Willemsen

**Department of Medical, Surgical and Health Sciences, University of Trieste, Trieste, Italy**

Marco Brumat, Giorgia Girotto

**Brigham and Women’s Hospital, Boston, MA, United States of America**

Julie E. Buring, Daniel I. Chasman, Lynda M. Rose, Paul M. Ridker

**Harvard Medical School, Boston, MA, United States of America**

Julie E. Buring, Daniel I. Chasman, Paul M. Ridker

**Department of Economics, New York University, New York, NY, United States of America**

David Cesarini

**Research Institute for Industrial Economics, Stockholm, Sweden**

David Cesarini

**National Bureau of Economic Research, Cambridge, MA, United States of America**

David Cesarini

**Department of Epidemiology, Harvard T.H. Chan School of Public Health, Boston, MA, United States of America**

Jorge E. Chavarro, Hongyang Huang, Peter Kraft, Stacey A. Missmer, Janet W. Rich-Edwards, Constance Turman

**Department of Nutrition, Harvard T.H. Chan School of Public Health, Boston, MA, United States of America**

Jorge E. Chavarro, Audrey J. Gaskins

**Channing Division of Network Medicine, Brigham and Women’s Hospital and Harvard Medical School, Boston, MA, United States of America**

Jorge E. Chavarro, Audrey J. Gaskins, Janet W. Rich-Edwards

**Institute for Maternal and Child Health IRCCS “Burlo Garofolo”, Trieste, Italy**

Massimiliano Cocca, Maria Pina Concas, Giorgia Girotto, Anna Morgan

**MRC Integrative Epidemiology Unit, University of Bristol, Bristol, United Kingdom**

George Davey-Smith, Fernando P. Hartwig, Susan M. Ring, Nicholas J. Timpson

**Lothian Birth Cohorts, Department of Psychology, University of Edinburgh, Edinburgh, United Kingdom**

Gail Davies, Ian J. Deary, W. David Hill, Paul Redmond

**Estonian Genome Center, University of Tartu, Tartu, Estonia**

Tõnu Esko, Reedik Mägi, Andres Metspalu, Evelin Mihailov, Lili Milani

**Broad Institute of the Massachusetts Institute of Technology and Harvard University, Cambridge, MA, United States of America**

Tõnu Esko

**Department of Epidemiology, Erasmus Medical Center, Rotterdam, The Netherlands**

Oscar Franco, M. Arfan Ikram, Maryam Kavousi

**Department of Epidemiology, Rollins School of Public Health, Emory University, Atlanta, GA, United States of America**

Audrey J. Gaskins

**Research Unit of Molecular Epidemiology, Helmholtz Zentrum München, German Research Center for Environmental Health, Neuherberg, Germany**

Christian Gieger, Brigitte Kühnel, Melanie Waldenberger

**Department of Psychiatry and Psychotherapy, University Medicine Greifswald, Greifswald, Germany**

Hans Jörgen Grabe

**Division of Research, Kaiser Permanente Northern California, Oakland, CA, United States of America**

Erica P. Gunderson

**Department of Sociology, Carolina Population Center, University of North Carolina at Chapel Hill, Chapel Hill, NC, United States of America**

Kathleen Mullan Harris

**Postgraduate Program in Epidemiology, Federal University of Pelotas, Pelotas, Brazil**

Fernando P. Hartwig, Bernando Lessa Horta

**University of Kentucky Markey Cancer Center, Lexington, KY, United States of America**

Chunyan He

**Department of Internal Medicine, Division of Medical Oncology, University of Kentucky College of Medicine, Lexington, KY, United States of America**

Chunyan He

**Department of Internal Medicine, Section of Gerontology and Geriatrics, Leiden University Medical Center, Leiden, The Netherlands**

Diana van Heemst, Raymond Noordam

**Interfaculty Institute for Genetics and Functional Genomics, University of Greifswald, Greifswald, Germany**

Georg Homuth

**Australian Centre for Precision Health, University of South Australia Cancer Research Institute, Adelaide, Australia**

Elina Hyppönen

**South Australian Health and Medical Research Institute, Adelaide, Australia**

Elina Hyppönen

**Department of Psychiatry, Amsterdam Public Health and Amsterdam Neuroscience, Amsterdam UMC, Vrije Universiteit, Amsterdam, The Netherlands**

Rick Jansen

**Department of Economics, Stockholm School of Economics, Stockholm, Sweden**

Magnus Johannesson

**Department of Bioinformatics, Isfahan University of Medical Sciences, Isfahan, Iran**

Zoha Kamali

**Department of Biostatistics, Harvard T.H. Chan School of Public Health, Boston, MA, United States of America**

Peter Kraft

**MRC Epidemiology Unit, Institute of Metabolic Science, Cambridge Biomedical Campus, University of Cambridge School of Clinical Medicine, Cambridge, United Kingdom**

Claudia Langenberg, Jian’an Luan, Nicholas J. Wareham, Jing Hau Zhao

Zoha Kamali, Lifelines Cohort Study, Peter J. van der Most, Ilja M. Nolte

**Department of Genetics, University of Groningen, University Medical Center Groningen, Groningen, The Netherlands**

Lifelines Cohort Study, Morris A. Swertz

**Psychiatric Genetics, QIMR Berghofer Medical Research Institute, Herston Brisbane, Queensland, Australia**

Penelope A. Lind, Sarah E. Medland

**Department of Medical Epidemiology and Biostatistics, Karolinska Institutet, Stockholm, Sweden**

Patrik K.E. Magnusson

**Wellcome Centre for Human Genetics, University of Oxford, Oxford, United Kingdom**

Anubha Mahajan, Mark I. McCarthy

**Oxford Centre for Diabetes, Endocrinology and Metabolism, Radcliffe Department of Medicine, University of Oxford, Oxford, United Kingdom**

Anubha Mahajan, Mark I. McCarthy

**Genetic Epidemiology, QIMR Berghofer Medical Research Institute, Herston Brisbane, Queensland, Australia**

Nicholas G. Martin

**Qatar Genome Programme, Qatar Foundation, Doha, Qatar**

Hamdi Mbarek

**School of Social and Community Medicine University of Bristol, Bristol, United Kingdom**

George McMahon

**Department of Integrative Physiology, University of Colorado at Boulder, Boulder, CO, United States of America**

Matthew B. McQueen

**Institute of Human Genetics, Helmholtz Zentrum München, German Research Center for Environmental Health, Neuherberg, Germany**

Thomas Meitinger

**Institute of Molecular and Cell Biology, University of Tartu, Tartu, Estonia**

Andres Metspalu

**Division of Adolescent and Young Adult Medicine, Department of Medicine, Boston Children’s Hospital and Harvard Medical School, Boston, MA, United States of America**

Stacey A. Missmer

**Department of Obstetrics, Gynecology, and Reproductive Biology, College of Human Medicine, Michigan State University, Grand Rapids, MI, United States of America**

Stacey A. Missmer

**Department of Sociology, University of Copenhagen, Copenhagen, Denmark**

Stine Møllegaard

**Department of Clinical Epidemiology, Leiden University Medical Center, Leiden, The Netherlands**

Dennis O. Mook-Kanamori, Renée de Mutsert

**Department of Public Health and Primary Care, Leiden University Medical Center, Leiden, The Netherlands**

Dennis O. Mook-Kanamori

**Institute of Clinical Chemistry and Laboratory Medicine, University Medicine Greifswald, Greifswald, Germany**

Matthias Nauck

**Department of Psychiatry, EMGO Institute for Health and Care Research and Neuroscience**

**Campus Amsterdam, VU University Medical Center/GGZ inGeest, Amsterdam, The Netherlands**

Brenda W.J.H. Penninx

**Institute of Epidemiology, Helmholtz Zentrum München, German Research Center for**

**Environmental Health, Neuherberg, Germany**

Annette Peters, Doris Stöckl, Melanie Waldenberger

**Population, Policy and Practice Research and Teaching Department, UCL Great Ormond Street**

**Institute of Child Health, London, United Kingdom**

Chris Power

**Division of Women’s Health, Department of Medicine, Brigham and Women’s Hospital and**

**Harvard Medical School, Boston, MA, United States of America**

Janet W. Rich-Edwards

**Erasmus University Rotterdam Institute for Behavior and Biology, Rotterdam, The Netherlands**

Cornelius A. Rietveld, A. Roy Thurik, André G. Uitterlinden

**Department of Applied Economics, Erasmus School of Economics, Rotterdam, The Netherlands**

Cornelius A. Rietveld, A. Roy Thurik

**deCODE Genetics/Amgen Inc., Reykjavik, Iceland**

Kári Stefánsson, Gudmar Thorleifsson, Unnur Thorsteinsdottir

**Institute of Medical Biostatistics, Epidemiology and Informatics (IMBEI), University Medical**

**Center, Johannes Gutenberg University, Mainz, Germany**

Konstantin Strauch

**Institute of Genetic Epidemiology, Helmholtz Zentrum München, German Research Center for**

**Environmental Health, Neuherberg, Germany**

Konstantin Strauch

**Chair of Genetic Epidemiology, IBE, Faculty of Medicine, LMU Munich, Germany**

Konstantin Strauch

**Institute for Community Medicine, University Medicine Greifswald, Greifswald, Germany**

Alexander Teumer

**Montpellier Business School, Montpellier, France**

A. Roy Thurik

**Department of Internal Medicine, Erasmus University Medical Center, Rotterdam, The Netherlands**

André G. Uitterlinden

## eQTLGen Consortium

Author list ordered alphabetically

Mawussé Agbessi, Habibul Ahsan, Isabel Alves, Anand Kumar Andiappan, Wibowo Arindrarto, Philip Awadalla, Alexis Battle, Frank Beutner, Marc Jan Bonder, Dorret I. Boomsma, Mark W. Christiansen, Annique Claringbould, Patrick Deelen, Tõnu Esko, Marie-Julie Favé, Lude Franke, Timothy Frayling, Sina A. Gharib, Greg Gibson, Bastiaan T. Heijmans, Gibran Hemani, Rick Jansen, Mika Kähönen, Anette Kalnapenkis, Silva Kasela, Johannes Kettunen, Yungil Kim, Holger Kirsten, Peter Kovacs, Knut Krohn, Jaanika Kronberg, Viktorija Kukushkina, Zoltan Kutalik, Bernett Lee, Terho Lehtimäki, Markus Loeffler, Urko M. Marigorta, Hailang Mei, Lili Milani, Grant W. Montgomery, Martina Müller-Nurasyid, Matthias Nauck, Michel G. Nivard, Brenda Penninx, Markus Perola, Natalia Pervjakova, Brandon L. Pierce, Joseph Powell, Holger Prokisch, Bruce M. Psaty, Olli T. Raitakari, Samuli Ripatti, Olaf Rotzschke, Sina Rüeger, Ashis Saha, Markus Scholz, Katharina Schramm, Ilkka Seppälä, Eline P. Slagboom, Coen D.A. Stehouwer, Michael Stumvoll, Patrick Sullivan, Peter A.C. ‘t Hoen, Alexander Teumer, Joachim Thiery, Lin Tong, Anke Tönjes, Jenny van Dongen, Maarten van Iterson, Joyce van Meurs, Jan H. Veldink, Joost Verlouw, Peter M. Visscher, Uwe Völker, Urmo Võsa, Harm-Jan Westra, Cisca Wijmenga, Hanieh Yaghootkar, Jian Yang, Biao Zeng, Futao Zhang

## Affiliations

**Computational Biology, Ontario Institute for Cancer Research, Toronto, Canada**

Mawussé Agbessi, Isabel Alves, Philip Awadalla, Marie-Julie Favé

**Department of Public Health Sciences, University of Chicago, Chicago, United States of America**

Habibul Ahsan, Brandon L. Pierce, Lin Tong

**Singapore Immunology Network, Agency for Science, Technology and Research, Singapore, Singapore**

Anand Kumar Andiappan, Bernett Lee, Olaf Rotzschke

**Leiden University Medical Center, Leiden, The Netherlands**

Wibowo Arindrarto, Bastiaan T. Heijmans, Eline P. Slagboom, Maarten van Iterson

**Department of Computer Science, Johns Hopkins University, Baltimore, United States of America**

Alexis Battle, Yungil Kim, Ashis Saha

**Departments of Biomedical Engineering, Johns Hopkins University, Baltimore, United States of America**

Alexis Battle

**Heart Center Leipzig, Universität Leipzig, Leipzig, Germany**

Frank Beutner

**Department of Genetics, University Medical Centre Groningen, Groningen, The Netherlands**

Marc Jan Bonder, Annique Claringbould, Patrick Deelen, Lude Franke, Urmo Võsa, Harm-Jan Westra, Cisca Wijmenga

**European Molecular Biology Laboratory, Genome Biology Unit, 69117 Heidelberg, Germany**

Marc Jan Bonder

**Netherlands Twin Register, Department of Biological Psychology, Vrije Universiteit Amsterdam, Amsterdam Public Health research institute and Amsterdam Neuroscience, the Netherlands**

Dorret I. Boomsma, Jenny van Dongen

**Cardiovascular Health Research Unit, University of Washington, Seattle, United States of America**

Mark W. Christiansen, Sina A. Gharib, Bruce M. Psaty

**Oncode Institute, Utrecht, The Netherlands**

Annique Claringbould, Patrick Deelen, Lude Franke, Harm-Jan Westra

**Genomics Coordination Center, University Medical Centre Groningen, Groningen, The Netherlands**

Patrick Deelen

**Department of Genetics, University Medical Centre Utrecht, P.O. Box 85500, 3508 GA, Utrecht, The Netherlands**

Patrick Deelen

**Estonian Genome Center, Institute of Genomics, University of Tartu, Tartu 51010, Estonia**

Tõnu Esko, Anette Kalnapenkis, Silva Kasela, Jaanika Kronberg, Viktorija Kukushkina, Lili Milani, Natalia Pervjakova, Urmo Võsa

**Genetics of Complex Traits, University of Exeter Medical School, Royal Devon & Exeter Hospital, Exeter, United Kingdom**

Timothy Frayling, Hanieh Yaghootkar

**Department of Medicine, University of Washington, Seattle, United States of America**

Sina A. Gharib

**School of Biological Sciences, Georgia Tech, Atlanta, United States of America**

Greg Gibson, Urko M. Marigorta, Biao Zeng

**MRC Integrative Epidemiology Unit, University of Bristol, Bristol, United Kingdom**

Gibran Hemani

**Amsterdam UMC, Vrije Universiteit, Department of Psychiatry, Amsterdam Public Health research institute and Amsterdam Neuroscience, The Netherlands**

Rick Jansen, Brenda Penninx

**Department of Clinical Physiology, Tampere University Hospital and Faculty of Medicine and Health Technology, Tampere University, Tampere, Finland**

Mika Kähönen

**University of Helsinki, Helsinki, Finland**

Johannes Kettunen

**Genetics and Genomic Science Department, Icahn School of Medicine at Mount Sinai, New York, United States of America**

Yungil Kim

**Institut für Medizinische InformatiK, Statistik und Epidemiologie, LIFE – Leipzig Research Center for**

**Civilization Diseases, Universität Leipzig, Leipzig, Germany**

Holger Kirsten, Markus Loeffler, Markus Scholz

**IFB Adiposity Diseases, Universität Leipzig, Leipzig, Germany**

Peter Kovacs

**Interdisciplinary Center for Clinical Research, Faculty of Medicine, Universität Leipzig, Leipzig, Germany**

Knut Krohn

**Lausanne University Hospital, Lausanne, Switzerland**

Zoltan Kutalik, Sina Rüeger

**Department of Clinical Chemistry, Fimlab Laboratories and Finnish Cardiovascular Research Center-Tampere, Faculty of Medicine and Health Technology, Tampere University, Tampere, Finland**

Terho Lehtimäki, Ilkka Seppälä

**Integrative Genomics Lab, CIC bioGUNE, Bizkaia Science and Technology Park, Derio, Bizkaia, Basque Country, Spain**

Urko M. Marigorta

**IKERBASQUE, Basque Foundation for Science, Bilbao, Spain**

Urko M. Marigorta

**Department of Medical Statistics and Bioinformatics, Leiden University Medical Center, Leiden, The Netherlands**

Hailang Mei

**Institute for Molecular Bioscience, University of Queensland, Brisbane, Australia**

Grant W. Montgomery, Peter M. Visscher, Jian Yang, Futao Zhang

**Institute of Genetic Epidemiology, Helmholtz Zentrum München-German Research Center for**

**Environmental Health, Neuherberg, Germany**

Martina Müller-Nurasyid

**Department of Medicine I, University Hospital Munich, Ludwig Maximilian’s University, München, Germany**

Martina Müller-Nurasyid, Katharina Schramm

**DZHK (German Centre for Cardiovascular Research), partner site Munich Heart Alliance, Munich, Germany**

Martina Müller-Nurasyid

**Institute of Clinical Chemistry and Laboratory Medicine, Greifswald University Hospital, Greifswald, Germany**

Matthias Nauck

**German Center for Cardiovascular Research (partner site Greifswald), Greifswald, Germany**

Matthias Nauck

**Department of Biological Psychology, Faculty of Behaviour and Movement Sciences, VU, Amsterdam, The Netherlands**

Michel G. Nivard

**National Institute for Health and Welfare, University of Helsinki, Helsinki, Finland**

Markus Perola

**Garvan Institute of Medical Research, Garvan-Weizmann Centre for Cellular Genomics, Sydney, Australia**

Joseph Powell

**Institute of Human Genetics, Helmholtz Zentrum München, Neuherberg, Germany**

Holger Prokisch

**Institute of Human Genetics, Technical University Munich, Munich, Germany**

Holger Prokisch

**Kaiser Permanente Washington Health Research Institute, Seattle, WA, United States of America**

Bruce M. Psaty

**Centre for Population Health Research, Department of Clinical Physiology and Nuclear Medicine, Turku University Hospital and University of Turku, Turku, Finland**

Olli T. Raitakari

**Statistical and Translational Genetics, University of Helsinki, Helsinki, Finland**

Samuli Ripatti

**Institute of Genetic Epidemiology, Helmholtz Zentrum München-German Research Center for Environmental Health, Neuherberg, Germany**

Katharina Schramm

**Department of Internal Medicine and School for Cardiovascular Diseases (CARIM), Maastricht University Medical Center, Maastricht, The Netherlands**

Coen D.A. Stehouwer

**Department of Medicine, Universität Leipzig, Leipzig, Germany**

Michael Stumvoll, Anke Tönjes

**Department of Medical Epidemiology and Biostatistics, Karolinska Institutet, Stockholm, Sweden**

Patrick Sullivan

**Center for Molecular and Biomolecular Informatics, Radboud Institute for Molecular Life Sciences, Radboud University Medical Center Nijmegen, Nijmegen, The Netherlands**

Peter A.C. ‘t Hoen

Alexander Teumer

**Institute for Laboratory Medicine, LIFE – Leipzig Research Center for Civilization Diseases,**

**Universität Leipzig, Leipzig, Germany**

Joachim Thiery

**Department of Internal Medicine, Erasmus Medical Centre, Rotterdam, The Netherlands**

Joyce van Meurs, Joost Verlouw

**UMC Utrecht Brain Center, University Medical Center Utrecht, Department of Neurology, Utrecht University, Utrecht, The Netherlands**

Jan H. Veldink

**Interfaculty Institute for Genetics and Functional Genomics, University Medicine Greifswald, Greifswald, Germany**

Uwe Völker

**School of Life Sciences, College of Liberal Arts and Science, University of Westminster, 115 New**

**Cavendish Street, London, United Kingdom**

Hanieh Yaghootkar

**Division of Medical Sciences, Department of Health Sciences, Luleå University of Technology, Luleå, Sweden**

Hanieh Yaghootkar

**Institute for Advanced Research, Wenzhou Medical University, Wenzhou, Zhejiang 325027, China**

Jian Yang

## BIOS Consortium (Biobank-based Integrative Omics Study)

**Management Team** Bastiaan T. Heijmans (chair), Peter A.C. ‘t Hoen, Joyce van Meurs, Aaron Isaacs, Rick Jansen, Lude Franke.

**Cohort collection** Dorret I. Boomsma, René Pool, Jenny van Dongen, Jouke J. Hottenga (Netherlands Twin Register); Marleen MJ van Greevenbroek, Coen D.A. Stehouwer, Carla J.H. van der Kallen, Casper G. Schalkwijk (Cohort study on Diabetes and Atherosclerosis Maastricht); Cisca Wijmenga, Lude Franke, Sasha Zhernakova, Ettje F. Tigchelaar (LifeLines Deep); P. Eline Slagboom, Marian Beekman, Joris Deelen, Diana van Heemst (Leiden Longevity Study); Jan H. Veldink, Leonard H. van den Berg (Prospective ALS Study Netherlands); Cornelia M. van Duijn, Bert A. Hofman, Aaron Isaacs, André G. Uitterlinden (Rotterdam Study).

**Data Generation** Joyce van Meurs (Chair), P. Mila Jhamai, Michael Verbiest, H. Eka D. Suchiman, Marijn Verkerk, Ruud van der Breggen, Jeroen van Rooij, Nico Lakenberg.

**Data management and computational infrastructure** Hailiang Mei (Chair), Maarten van Iterson, Michiel van Galen, Jan Bot, Dasha V. Zhernakova, Rick Jansen, Peter van ‘t Hof, Patrick Deelen, Irene Nooren, Peter A.C. ‘t Hoen, Bastiaan T. Heijmans, Matthijs Moed.

**Data Analysis Group** Lude Franke (Co-Chair), Martijn Vermaat, Dasha V. Zhernakova, René Luijk, Marc Jan Bonder, Maarten van Iterson, Patrick Deelen, Freerk van Dijk, Michiel van Galen, Wibowo Arindrarto, Szymon M. Kielbasa, Morris A. Swertz, Erik. W van Zwet, Rick Jansen, Peter-Bram ‘t Hoen (Co-Chair), Bastiaan T. Heijmans (Co-Chair).

### Affiliations

**Molecular Epidemiology Section, Department of Medical Statistics and Bioinformatics, Leiden University Medical Center, Leiden, The Netherlands**

Bastiaan T. Heijmans, P. Eline Slagboom, Marian Beekman, Joris Deelen, H. Eka D.

Suchiman, Ruud van der Breggen, Nico Lakenberg, Maarten van Iterson, Matthijs Moed, René Luijk

**Department of Human Genetics, Leiden University Medical Center, Leiden, The Netherlands**

Peter A.C. ‘t Hoen, Michiel van Galen, Martijn Vermaat, Peter-Bram ‘t Hoen

**Department of Internal Medicine, ErasmusMC, Rotterdam, The Netherlands**

Joyce van Meurs, André G. Uitterlinden, P. Mila Jhamai, Michael Verbiest, Marijn Verkerk, Jeroen van Rooij

**Department of Genetic Epidemiology, ErasmusMC, Rotterdam, The Netherlands**

Aaron Isaacs, Cornelia M. van Duijn

**Department of Psychiatry, VU University Medical Center, Neuroscience Campus Amsterdam, Amsterdam, The Netherlands**

Rick Jansen

**Department of Genetics, University of Groningen, University Medical Centre Groningen, Groningen, The Netherlands**

Lude Franke, Cisca Wijmenga, Sasha Zhernakova, Ettje F. Tigchelaar, Dasha V. Zhernakova, Patrick Deelen, Marc Jan Bonder

**Department of Biological Psychology, VU University Amsterdam, Neuroscience Campus Amsterdam, Amsterdam, The Netherlands**

Dorret I. Boomsma, René Pool, Jenny van Dongen, Jouke J. Hottenga

Marleen MJ van Greevenbroek, Coen D.A. Stehouwer, Carla J.H. van der Kallen, Casper G. Schalkwijk

**Department of Gerontology and Geriatrics, Leiden University Medical Center, Leiden, The Netherlands**

Diana van Heemst

**Department of Neurology, Brain Center Rudolf Magnus, University Medical Center Utrecht, Utrecht, The Netherlands**

Jan H. Veldink, Leonard H. van den Berg

**Department of Epidemiology, ErasmusMC, Rotterdam, The Netherlands**

Bert A. Hofman

**Sequence Analysis Support Core, Leiden University Medical Center, Leiden, The Netherlands**

Hailiang Mei, Peter van ‘t Hof, Wibowo Arindrarto

**SURFsara, Amsterdam, The Netherlands**

Jan Bot, Irene Nooren

**Genomics Coordination Center, University Medical Center Groningen, University of Groningen, Groningen, The Netherlands**

Freerk van Dijk, Morris A. Swertz

**Medical Statistics Section, Department of Medical Statistics and Bioinformatics, Leiden University Medical Center, Leiden, The Netherlands**

Szymon M. Kielbasa, Erik. W van Zwet

## Supplementary Information

### Supplementary Tables

**Supplementary Tables S1-S19C.**

## Notes

### Summary of Updates

Additional bi-directional MR & correlation analyses related to infertility, Figure 2 & 3 revised, some methods description moved from SI to main text, clarification column names in Supplemental Table 15E

