## Supplementary Information for "Identification of 370 genetic loci for age at first sex and birth linked to externalising behaviour"

[1] Leverhulme Centre for Demographic Science, University of Oxford, Oxford, United Kingdom
[2] Nuffield College, University of Oxford, Oxford, United Kingdom
[3] École Nationale de la Statistique et de L'administration Économique (ENSAE), Paris, France
[4] Center for Research in Economics and Statistics (CREST), Paris, France
[5] The Beijer Laboratory and Department of Immunology, Genetics and Pathology, Uppsala University and SciLifeLab, Uppsala, Sweden
[6] Department of Epidemiology, University of Groningen, University Medical Center Groningen, Groningen, The Netherlands
[7] Department of Bioinformatics, Isfahan University of Medical Sciences, Isfahan, Iran
[8] The Novo Nordisk Foundation Center for Basic Metabolic Research, Faculty of Health and Medical Sciences, University of Copenhagen, Copenhagen, Denmark
[9] Department of Epidemiology Research, Statens Serum Institut, Copenhagen, Denmark
[10] MRC Epidemiology Unit, Institute of Metabolic Science, University of Cambridge, Cambridge, United Kingdom
[11] Department of Statistical Sciences, University of Bologna, Italy

† Denotes equal contribution

### List of Supplementary Figures

Figure S1. Summary and description of methods and main results, GWAS of age at first sex (AFS) and age at first birth (AFB)
Figure S2. Correlation plot between Age at first birth and Age at first sex by birth cohort, UK Biobank
Figure S3. Age at first birth (AFB) panel A and Age at first sex (AFS) panel B by birth cohort, UK Biobank
Figure S4A. SNP heritability for AFB, women in the UK Biobank
Figure S4B. SNP heritability for AFS, women and men in UK Biobank
Figure S5A-C. Manhattan plots, Age at first sex (AFS), Pooled (A), Women (B) and Men (C)
Figure S6A-C. Manhattan plots, Age at first birth (AFB), Pooled (A), Women (B) and Men (C)
Figure S7. Variance explained from Polygenic scores for Age at First Birth and Age at First Sex using PRSice, LDPred and MTAG+LDPred in out-of-sample cohorts
Figure S8. Nelson-Aalen hazard estimates of first sex by age. Comparison between top 5% and bottom 5% PGS of age at first sex
Figure S9. Nelson-Aalen hazard estimates of first birth by age. Comparison between top 5% and bottom 5% PGS of age at first birth
Figure S10A. AFB PGS score by percentile groups and parent's educational level
Figure S10B. AFS PGS score by percentile groups and parent's educational level
Figure S11. Genetic correlations and SNP heritabilities between and among reproductive, behavioural, psychiatric, substance use, personality and anthropometric traits
Figure S12A. A path diagram showing the structure of the genetic multiple regression model fit to EA and AFB
Figure S12B. A path diagram showing the structure of the genetic multiple regression model fit to AFS and EA
Figure S13. A heat map showing the genetic correlations between and among the fertility GWAS phenotypes, the sex hormone phenotypes, and other phenotypes related to reproductive biology, as calculated by LD score regression.
Figure S14. A path diagram for a GenomicSEM model of the relative associations of an externalizing latent factor, age at menopause, and age at menarche with age at first birth in women
Figure S15. A heat map showing the genetic correlations between and among the fertility GWAS phenotypes, the sex hormone phenotypes, and other phenotypes related to reproductive biology, as calculated by LD score regression.
Figure S16. Coefficients (and Cis) of bi-directional MR of human reproductive behaviour (AFB, AFS), age initiated smoking and educational attainment on Type 2 diabetes and Coronary Artery Disease later in life
Figure S17. Protein-protein interactions identified using STRING for genes that are highly expressed at the protein level in: A) brain and result in a nervous system or neurological behavior phenotype in mutant mice; B) glands and result in an endocrine/exocrine phenotype in mutant mice; C-D) female (C) or male (D) reproductive organs and result in a reproductive phenotype in mutant mice. Pink lines highlight experimentally determined interactions.
Figure S18. Genetic overlap amongst the sexes for AFS and AFB, LD score bivariate regression
Figure S19. Gene prioritization of AFS and AFB by sex

### List of Supplementary Tables

Table S1. Description of participating cohorts

Table S2 Cohort phenotype description

Table S3a. Sample size individuals with automsome chromosome information in participating cohorts

Table S3b. Sample size individuals with sex chromosome information in participating cohorts

Table S4. Genotyping and imputation

Table S5. Description of SNP filtering and cohort exclusion for age at first birth (AFB) analyses for women

Table S6. Description of SNP filtering and cohort exclusion for age at first sex (AFS) analyses for women

Table S7. Description of SNP filtering and cohort exclusion for age at first birth (AFB) analyses for women

Table S8. Description of SNP filtering and cohort exclusion for age at first sex (AFS) analyses for women

Table S9. Association Results for 88 independent SNPs that reached genome-wide significance (P < 5×10-8) in the pooled-sex GWAS of Age at First birth (AFB), AFB Males and Females

Table S10. Association Results for 261 independent SNPs that reached genome-wide significance (P < 5×10-8) in the pooled-sex GWAS of Age at First Sex (AFS), AFS Males and Females

Table S11. Genetic correlations (rg) AFB and AFS with selected phenotypes

Table S12A . Unstandardized results from genetic multivariate regression models examining the relationship between AFS in males and EA, accounting for the genetic correlation of AFS with a third phenotype

Table S12B. Unstandardized results from genetic multivariate regression models examining the relationship between EA and AFB, accounting for the genetic correlation of EA with a third phenotype

Table S12C. Standardized results from genetic multivariate regression models examining the relationship between AFS and EA, accounting for the genetic correlation of AFS with a third phenotype

Table 12D. Unstandardized results from genetic multivariate regression models examining the relationship between AFS and EA, accounting for the genetic correlation of AFS with a third phenotype

Table S12E. Standardized results from genetic multivariate regression models examining the relationship between EA and AFB in males, accounting for the genetic correlation of EA with a third phenotype

Table S12F. Unstandardized results from genetic multivariate regression models examining the relationship between EA and AFB in males, accounting for the genetic correlation of EA with a third phenotype

Table S12G. Standardized results from genetic multivariate regression models examining the relationship between EA and AFB in females, accounting for the genetic correlation of EA with a third phenotype

Table 12H. Unstandardized results from genetic multivariate regression models examining the relationship between EA and AFB in females, accounting for the genetic correlation of EA with a third phenotype

Table 12I. Standardized results from genetic multivariate regression models examining the relationship between AFS in males and EA, accounting for the genetic correlation of AFS with a third phenotype

Table 12J. Unstandardized results from genetic multivariate regression models examining the relationship between AFS in males and EA, accounting for the genetic correlation of AFS with a third phenotype

Table 12K. Standardized results from genetic multivariate regression models examining the relationship between AFS in females and EA, accounting for the genetic correlation of AFS with a third phenotype

Table 12L. Unstandardized results from genetic multivariate regression models examining the relationship between AFS in females and EA, accounting for the genetic correlation of AFS with a third phenotype

Table 13A. Bi-Directional MR, Years of education and AFB and AFB/AFS with risk taking and age at smoking initiation

Table S13B. Mendelian Randomization (MR) of age at first birth (AFB) to Coronary arterty disease (CAD) and Type 2 diabetes (T2D) and age at first sex (AFS) to CAD and T2D, and Educational Attainment to CAD and T2D

Table S14. Polygenic score (PGS) prediction of age at first birth (AFB), educational attainment (EA) and risk on parental longevity

Table S15A. Results from CELLECT tissue enrichment analysis for age at first sex (AFS)

Table S15B. Results from CELLECT tissue enrichment analysis for age at first birth (AFB)

Table 15C. Results from CELLECT gene prioritization for age at first sex (AFS)

Table S15D. Results from CELLECT gene prioritization for age at first birth (AFB)

Table S15E. Results from CELLECT cell type enrichment analysis using mouse brain RNA sequencing data for age at first sex (AFS)

Table S15F. Results from CELLECT cell type enrichment analysis using mouse brain RNA sequencing data for age at first birth (AFB)

Table S15G. Results from CELLECT cell type enrichment analysis using tabula muris RNA sequencing data for age at first sex (AFS)

Table S16A. Search terms used for the Phenolyzer analysis for the three areas of interest

Table S16B. Results of Phenolyzer analysis age at first birth (AFB) and age at first sex (AFS)

Table S17A. The results of in silico sequencing and in silico lookup of GWAS associations of AFB. AF_EUR indicates the allele frequency of the alternative allele (A2) in the European population.

Table S17B. The results of in silico sequencing and in silico lookup of GWAS associations of AFS. AF_EUR indicates the allele frequency of the alternative allele (A2) in the European population.

Table S18A. Summary data-based Mendelian Randomization (SMR) for age at first sex (AFS)

Table S18B. Summary data-based Mendelian Randomization (SMR) for age at first birth (AFB)

Table S19A. Summary of gene prioritization results across all approaches for age at first sex (AFS)

Table S19B. Summary of gene prioritization results across all approaches for age at first birth (AFB)

Table S19C. Summary of gene prioritization results across all approaches for age at first sex (AFS) and age at first birth (AFB)

### 1. Background and Phenotype Definitions

#### 1.1 Background

Previous studies have shown that the onset of human reproductive behaviour – age at first sexual intercourse (AFS) and age at first birth (AFB) – have a genetic basis. AFB has a SNP-heritability of 15%[1] and AFS 15-17% (see Section 3), with two genome-wide association studies (GWAS) in 2016 identifying 10 genetic loci linked to AFB[2] and 38 associated with AFS.[3] A detailed discussion of the motivation behind the study of these traits including the evolutionary causes of genetic variance in reproductive behaviour, additive and dominant genetic variation and environmental variation in fertility behaviour can be found in the Supplementary Note of Barban et al. (2016).[2] A description of the data and methods used in this study can be found in the online Methods section appended to the article.

The current analysis extends previous work in several appreciable ways. First, this study has a sizeable increase in sample size, making this the largest GWAS to date on these phenotypes. Previous work on AFS[3] examined a small sample of 125,667 individuals from the UK Biobank with a study on AFB[2] examining 251,151 individuals. The current study is considerably larger for both AFS (N=397,338 pooled; N=214,547 women; N=182,791) and AFB (N=542,901 pooled; N=418,758 women; N=124,008 men). A second extension is that we use 1000G imputed genotype data, which in addition to the larger sample, allows us to detect considerably more signals. Third, we include an X-Chromosome analysis, allowing us to uncover additional novel loci. A fourth advance is the ability to find markedly more biological signals. Fifth, our extensive analyses of the correlation and underlying etiology of these traits reveals an underlying genetic basis of AFS and AFB with other traits. This includes externalizing behaviour and substance use for early AFS and AFB and links to internalizing traits and infertility disease for later AFS and AFB. Sixth, we show that AFB is a stronger predictor for late age onset of disease and parental longevity, even beyond known standard predictors such as educational attainment. Finally, we demonstrate how that our polygenic scores are sensitive to gene-environment correlation (rGE) and childhood socioeconomic status.

#### 1.2 Phenotype Definitions

An overview of participating cohorts is found in Table S1, with a description provided shortly in Section 3. The detailed phenotype definitions and questions drawn from each of the cohorts are included in Supplementary Table S2.

**Age at first sexual intercourse (AFS)** is treated as a continuous measure and assessed using questions such as *What was your age when you first had sexual intercourse?* This is often defined by more detailed divisions such as (sexual intercourse includes vaginal, oral or anal intercourse). Ages less than 12 are normally excluded. The UKBiobank requires confirmation of ages in the range 4-12, and excludes all answers that were less than 4. For out of sample replication, other studies include 12 as the minimum age, if they do not have a study specific lower limit. Age at first sexual intercourse tends to have a markedly non-normal distribution, so a within-sex inverse rank normal transformation is required.

**Age at first birth (AFB)** is treated as a continuous measure either asked directly or created from several survey questions (e.g., birthdate of participant and date of birth of first child). The most common question was: *How old were you when you had your first child? Or What is the date of birth of your first child?* Individuals were eligible for inclusion if they were assessed for AFB and had given birth to a child.

#### 2. Summary of Methods, Purpose of Analysis and Main Results

The Methods section in the main article describes all methods used in this paper in detail, summarised in Figure S1. In this figure, dark grey boxes indicate the method of analysis, light grey the purpose of the analysis and white a summary of the main results.

**Historical changes.** We first examined historical and phenotypic changes in the age distributions for these phenotypes, followed by genotypic changes, estimating heritability by sex and across birth cohorts.

**Polygenic score (PGS) construction and prediction.** Following the GWAS, we then engaged in a variety of techniques to interrogate the PGS prediction. We constructed the PGS using multiple techniques and test out of sample prediction. We applied the LD Score intercept test to test for population stratification followed by survival models to examine the impact of right-censoring and sex differences in PGS prediction. We also studied the sensitivity of our PGSs by childhood socioeconomic status.

**Correlation, etiology, causality and prediction.** This was followed by five additional analyses to explore various substantive and methodological questions. We used LD score regression to examine the genetic correlations between traits. We engaged in Genomic SEM to understand the shared genetic etiology of related traits. Bi-directional Mendelian Randomization was used to measure causal pathways of our phenotypes with educational attainment, age at initiation of smoking and risk taking and whether our PGSs had independent effects on later life diseases (type 2 diabetes, coronary artery disease), once educational attainment and BMI were controlled for. Exploratory Factor Analysis allowed us to breakdown the underlying components of reproductive behaviour into those related to externalizing or disinhibition versus biological components. Finally, we estimated survival models to examine whether reproductive timing was linked to parental longevity.

**Biological annotation.** We also carried out a variety of biological analyses. This included DEPICT for candidate gene identification, CELLECT RNAseq mouse brain and Tabula muris RNAseq to identify enriched mouse nervous system cell types. Phenolyzer was used to prioritize candidates using prior knowledge of these phenotypes using machine learning on seed genes and predicted gene rankings. In silico sequencing allowed us to identify non-synonymous variants and summary-based Mendelian Randomization (SMR), HEIDI and eQTL to find evidence of gene expression. Sex-specific effects were identified using LD score regression. We then integrated of these biological results to prioritize the genes from all approaches and examine protein expression. This allowed us to identify the key genes related to reproduction and externalizing behaviour as well as gene prioritization of sex-specific loci.

### 3. Phenotypic and genotypic changes in the onset of human reproductive behaviour over time

#### 3.1 Phenotypic changes in the onset of human reproductive behaviour

Over the past forty years, there has been a rapid postponement of age at first birth (AFB) by 4-5 years to a mean AFB for women around 29 many advanced societies.[4] The biological ability to conceive a child already starts to decline for many women as early as 25, with around 50% of women sterile by the age of 40.[5] This postponement has been related to multiple social, economic, and cultural factors, which has been documented in several detailed reviews.[4,6] A central factor is the introduction of effective contraception and ability to control fertility and engage in individual choice since the late 1960s. Another key factor is the well-documented association between women's gains in educational attainment and that relationship with later fertility, particularly for more recent birth cohorts. This is related to women's stronger labour market attachment and their realization that fertility postponement avoids large motherhood wage penalties. In fact, by each a year a woman delays motherhood, she increases her career earnings by 9%.[6] Other factors are the strong cultural and ideational changes and norms surrounding sexual behaviour, entry into parenthood and the role of children who are often no longer strongly required for economic and labour support to parents. Finally, multiple structural factors such as the availability of childcare, gender equity, housing and resources all play a vital role.

Figure S2 documents how sexual debut was linked to first childbirth in earlier birth cohorts (<1941, r=0.60) to a relative uncoupling in more recent birth cohorts (>1960, r=0.31). Related to this is a large body of demographic work that has examined the decoupling of sex with marriage.[6]

Figure S2 illustrates the gradual decoupling of sexual initiation with reproduction. Here we see that the correlation or timing between AFS and AFB was concentrated and closer together in earlier birth cohorts whereas with more recent birth cohorts, it is increasingly more widely distributed over time. In other words, the classic association of sexual behaviour with marriage and childbearing held in earlier cohorts has waned over time, largely due to the introduction of effective contraception and changes in social norms about sexual behaviour outside of a marital union.[6] Note that the implausible 'immaculate conception' outliers (i.e., age at first birth before first sex) shown in this figure were removed prior to GWAS analysis.

Figure S3 (Panel A) examines phenotypic data from the UK Biobank and shows the shift in the distribution of AFB not only to later to ages, but also a wider spread in the distribution itself. Figure S3 (Panel B) of AFS shows that in earlier cohorts, there was a bi-modal distribution, one which had earlier sexual intercourse often tied to socio-economic circumstances, problem or risky behaviour.[7] The other group engaging in later sexual initiation, has been found to be tied to higher educational goals and achievement with early sexual intercourse tied to with higher calculated risk of pregnancy, which would disturb longer-term life planning and career goals.[8] The panel also shows a narrowing of the distribution over time to earlier ages.

Figure S1. Summary and description of methods and main results, GWAS of age at first sex (AFS) and age at first birth (AFB)

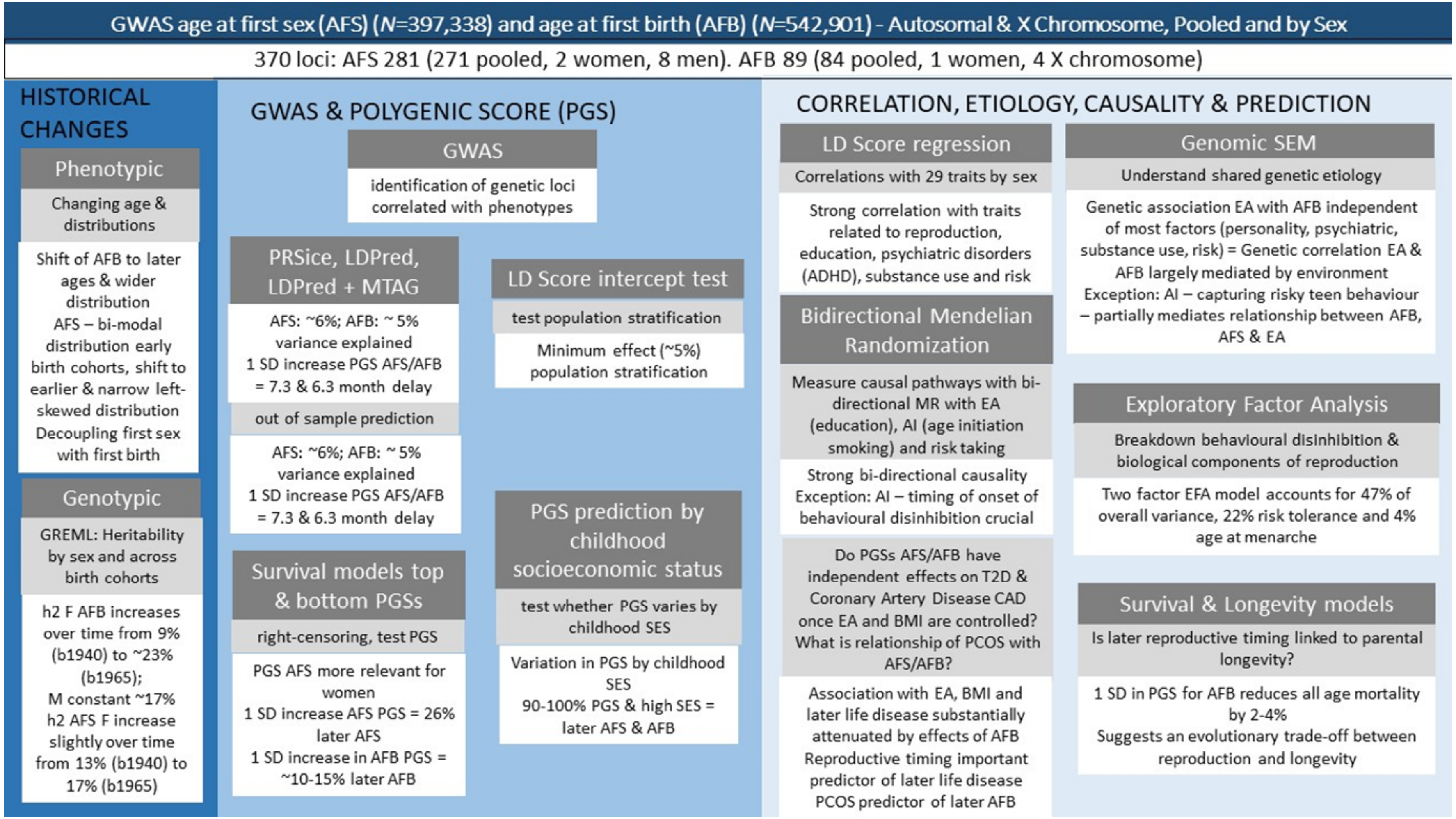

Dark grey indicates analysis method, light grey the purpose of the analysis and white the main results. AFB (age at first birth); AFS (age at first sex); PGS (polygenic score); MR (Mendelian Randomisation); EA (Educational Attainment); SES (socioeconomic status); EFA (Exploratory Factor Analysis); BMI (Body Mass Index); PCOS (polycystic ovarian syndrome); SD (Standard Deviation); F (females); M (males); SEM (structural equation model); LD (linkage disequilibrium)

Figure S1. Continued, Summary and description of methods and main results, GWAS of age at first sex (AFS) and age at first birth (AFB).

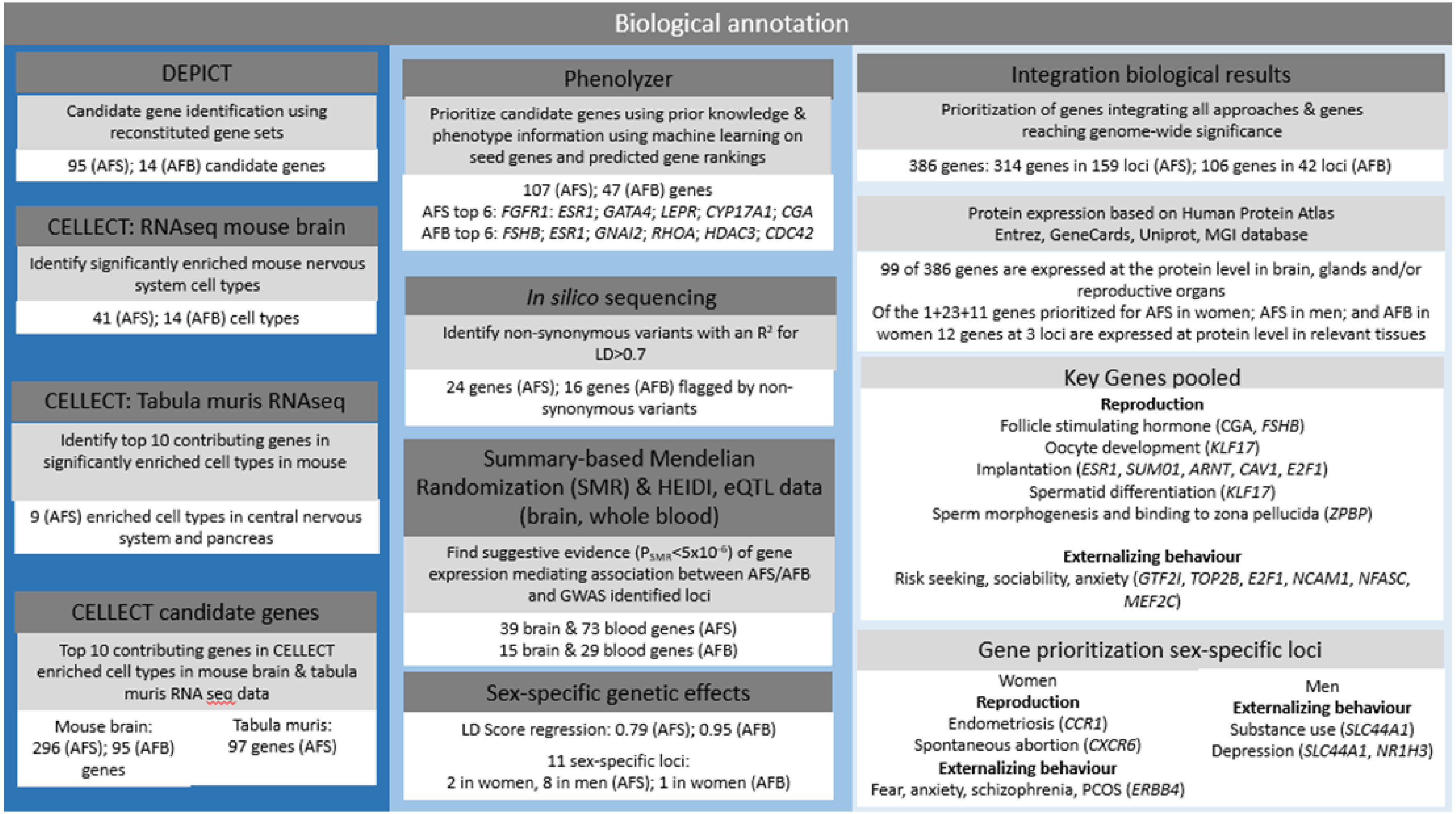

Dark grey indicates analysis method, light grey the purpose of the analysis and white the main results. AFB (age at first birth); AFS (age at first sex); PGS (polygenic score); MR (Mendelian Randomisation); EA (Educational Attainment); SES (socioeconomic status); EFA (Exploratory Factor Analysis); BMI (Body Mass Index); PCOS (polycystic ovarian syndrome); SD (Standard Deviation); F (females); M (males); SEM (structural equation model); LD (linkage disequilibrium)

Figure S2. Correlation plot between Age at first birth and Age at first sex by birth cohort, UK Biobank

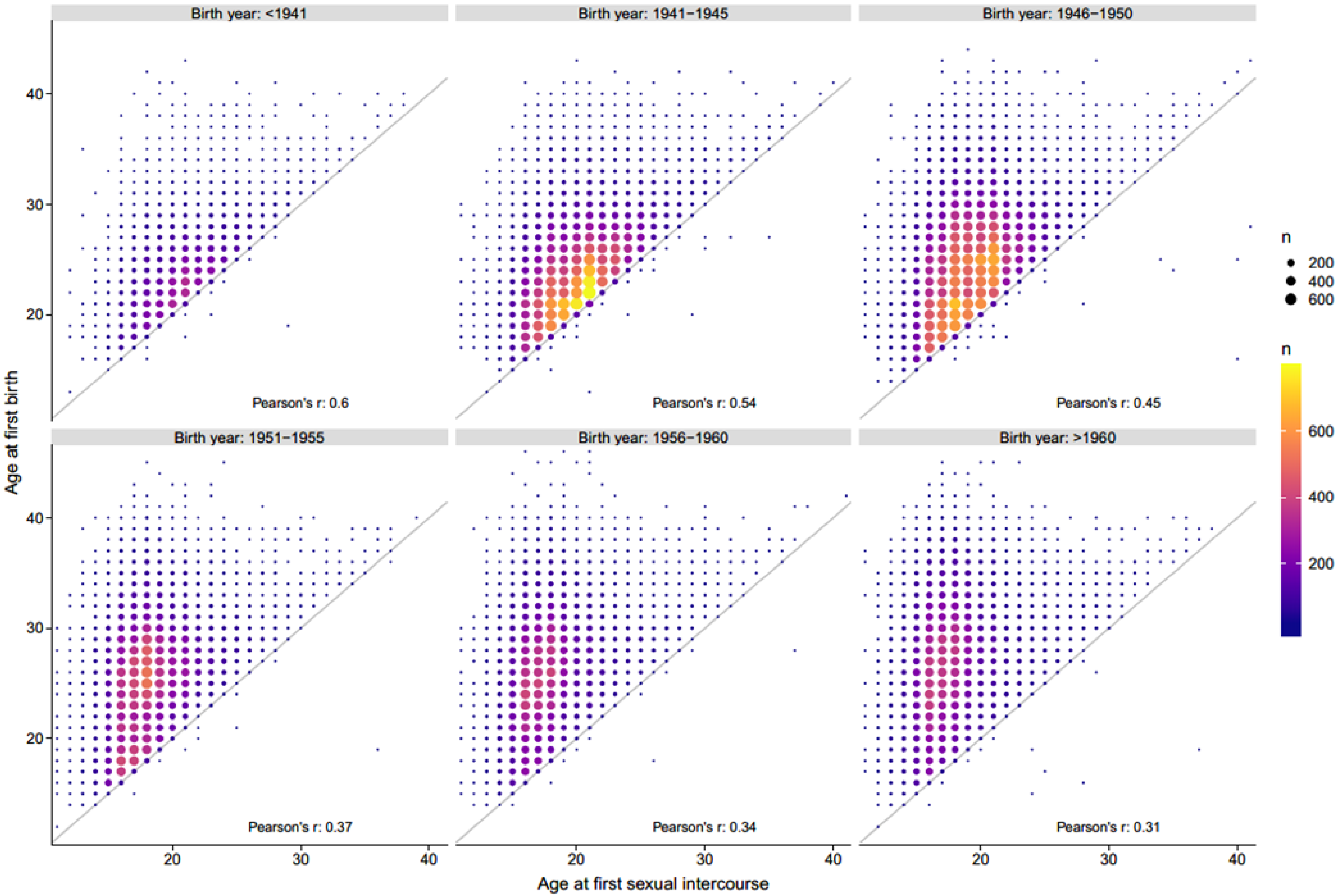

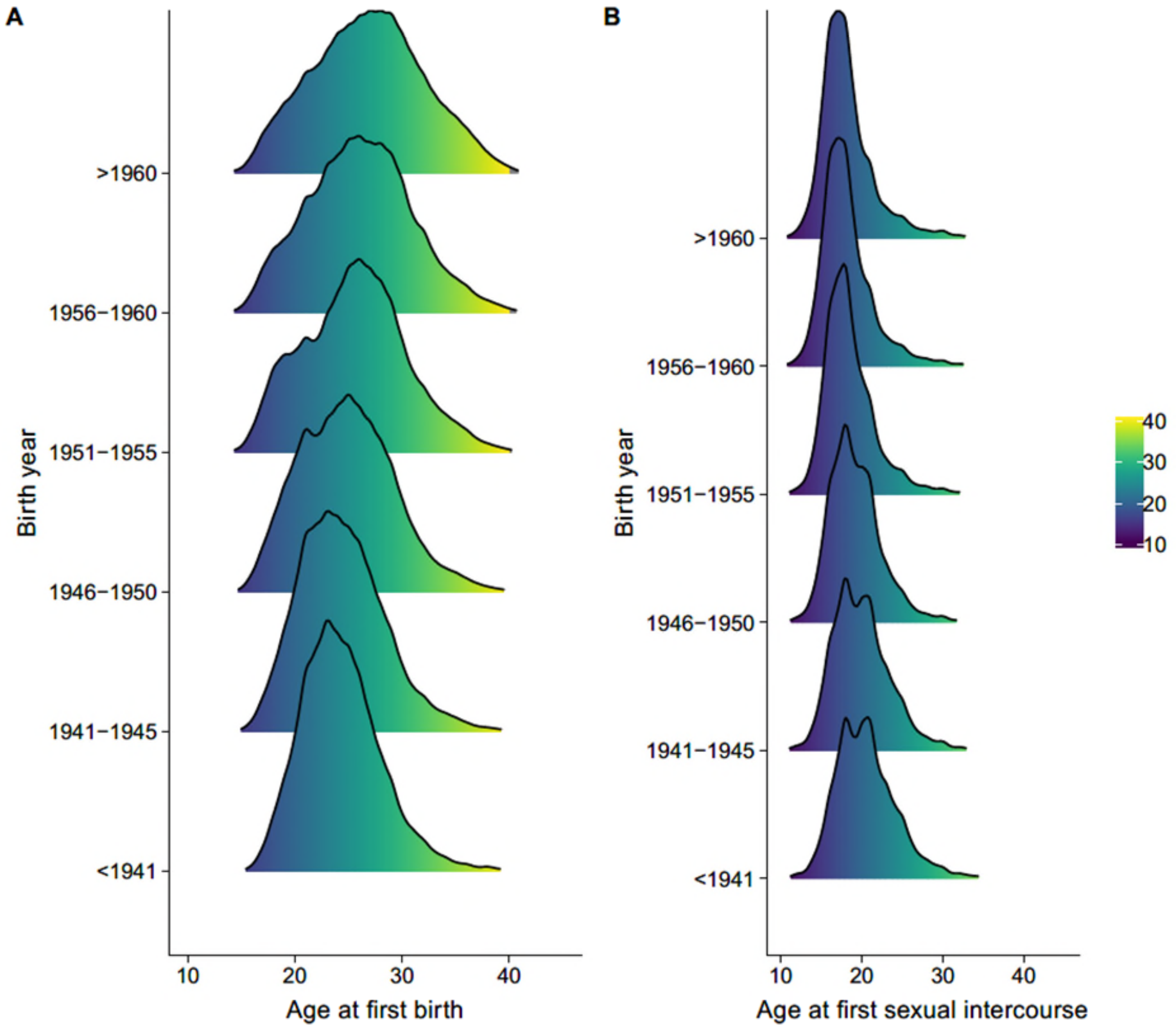

Figure S3. Age at first birth (AFB) panel A and Age at first sex (AFS) panel B by birth cohort, UK Biobank

#### 3.2 Heterogeneity in heritability across birth cohorts

A recent study demonstrated that estimates from GWAS discoveries are substantially smaller across populations compared to within populations.[9] Simulations showed that the results reflected heterogeneity in gene – environment interaction rather than genetic heterogeneity. In other words, particularly for complex traits and diseases such as reproductive behaviour or others such as educational attainment or BMI, it is more difficult to determine the influence of genetic versus socio-environmental factors. That study demonstrated that although GWA studies combine data from individuals across different time periods, it is implausible to assume that genetic effects are uniform across time.

To test whether this was a concern with our current analysis, Figures S4A and B show how the SNP heritability estimates change over time for AFB and AFS using the UK Biobank. Our SNP heritability estimate refers to the proportion of the additive genetic variance explained by common SNPs across the genome over the overall phenotypic variance ($\sigma_{\mathbf{Y}}^2$) of the trait: [10]

$$h_{SNP}^2 = \frac{\sigma_{\mathbf{G}}^2}{\sigma_{\mathbf{Y}}^2}$$

The phenotypic variance is the sum of additive genetic and environmental variance, i.e., $\sigma_{\mathbf{Y}}^2 = \sigma_{\mathbf{G}}^2 + \sigma_{\mathbf{E}}^2$ , where $\sigma_{\mathbf{G}}^2$ is the additive genetic variance explained by all common SNPs across the genome and $\sigma_{\mathbf{E}}^2$ is the residual variance. The methods we applied have been detailed elsewhere. [10–14] Briefly, we applied a linear mixed model

$$\mathbf{y} = \mathbf{X}\boldsymbol{\beta} + \mathbf{g} + \mathbf{e}$$

where **y** is an *N*×1 vector of dependent variables, *N* is the sample size, **β** is a vector for fixed effects of the *M* covariates in *N*×*M* matrix **X** (including the intercept and potential confounders such as birth year), **g** is the *N*×1 vector with each of its elements being the total genetic effect of all common SNPs for an individual, and **e** is an *N*×1 vector of residuals. We have **g**~*N*(0, $\mathbf{A}\sigma_{\mathbf{G}}^2$) and e~*N*(0, $\mathbf{I}\sigma_{\mathbf{E}}^2$). Hence, the variance matrix $\mathbf{V}$ of the observed phenotypes is:

$$\mathbf{V} = \mathbf{A}\sigma_{\mathbf{G}}^2 + \mathbf{I}\sigma_{\mathbf{E}}^2,$$

We used BOLT software[15] as an efficient solution for mixed-linear models and controlled for the first 20 principal components as well as assessment centre, chip and birth year of the participants. We only included individuals who self-reported as British individuals to reduce heterogeneity due to cultural background.

Figure S4A illustrates a steady increase in SNP-heritability by birth cohort for AFB of women (N = 164,486) from 9% [CI = 4-14] for those born in 1940, climbing to around 22% [19-25] for the latest cohorts born in 1965. We note, however, that these results should also be considered in relation to healthy volunteer sample selection and thus ascertainment bias in the UK Biobank, with replication across other large samples vital for future research. Importantly, the genetic correlation across birth cohorts in not significantly different from 1. For individuals born before and after 1950, for example, rG = 0.93 [0.84-1.01], the same genes appear to have gained in importance for AFB in women in the UK.

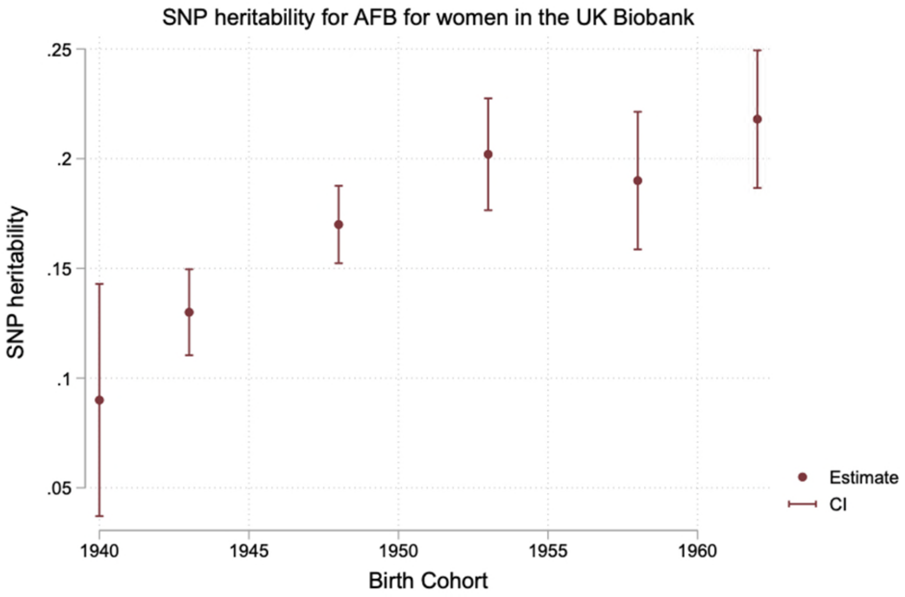

Figure S4A. SNP heritability for AFB, women in UK Biobank

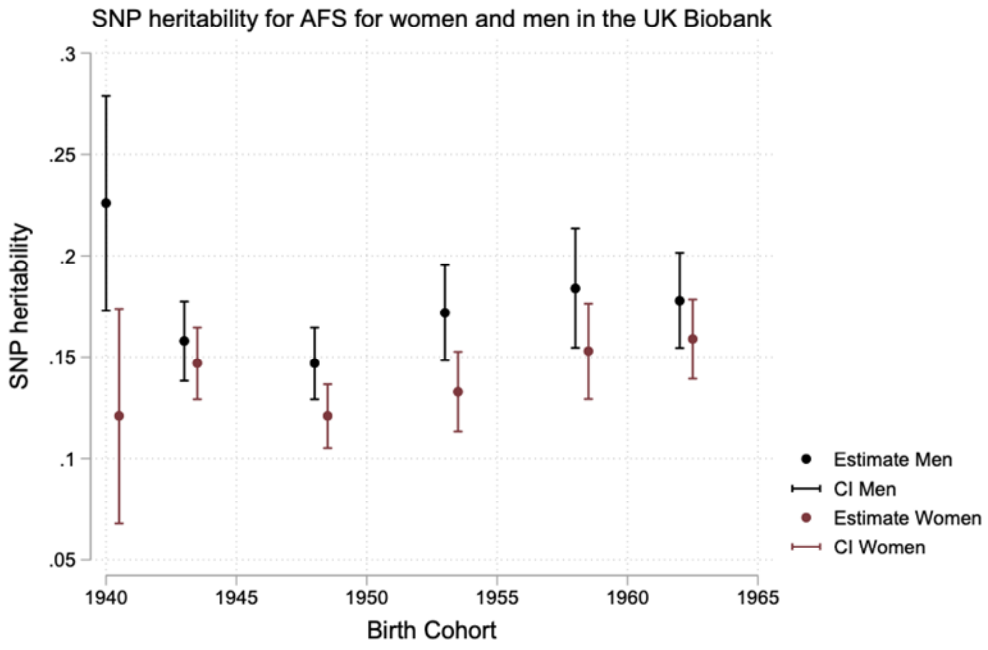

Figure S4B. SNP heritability for AFS, women and men in UK Biobank

Figure S4B shows the SNP heritability of AFS for men (178,988) and women (210,363), respectively, across birth cohorts. For men we observe a slight u-shaped pattern, with heritability ranging between 23 [17-28] % to just under 15% [13-17]. For women the trend is similar to AFB with a relatively steady increase in heritability by birth cohort over time from around 12% [6-18] in the oldest cohorts born in 1940 to around 16% [14-17] for those born in 1965 with a significant increase since the end of the 1940s (12 [11-14]). Again, genetic correlations from before 1950 and after could not be statistically distinguished from 1, both for women (rG = .96 [.91-1.05]) and men (0.98 [.90-1.05]. This is in line with work that has shown changes in genetic associations with smoking over time and across cohorts.[16,17]

### 4. Overview of GWAS meta-analysis

The discovery of genetic variants associated with AFB and AFS is based on cohort-level genome-wide association studies that were quality-controlled and meta-analysed by two separate independent centres at the University of Oxford and University of Cambridge. We followed standard QC protocol and employed the software packages QCGWAS[18] and EasyQC,[19] which allowed us to harmonize the files and identify possible sources of errors in association results. This procedure entailed that diagnostic graphs and statistics were generated for each set of GWAS results (i.e., for each file). In the case where apparent errors could not be amended by stringent QC, cohorts were excluded from the meta-analysis.

#### 4.1 Participating cohorts

A total of 36 cohorts participated in our study, with phenotype inclusion varying according to the availability of the phenotype. Table S1 provides a description of the cohorts, including the type of sampling, country, coverage of birth years of respondents, mean age and standard deviation and scientific reference for more information on each study. Table S2 provides details on the specific phenotype descriptions for each cohort. Table S3a-b provide the sample sizes of the adjusted pooled analysis and by women and men in the case of family data. Cohorts who agreed to participate followed an Analysis Plan posted on the Open Science Framework preregistration site https://osf.io/b4r4b/ on February 08, 2017. Although AFB and number of children ever born (NEB) were examined together in our previous research and considered in the additional analysis plan[2], due to the strong relationship between AFS and AFB, number of children ever born (NEB) and childlessness (CL) were examined separately in another paper.

As Table S3a shows, for autosomal chromosomes the total number of individuals in the pooled meta-analysis for AFB was 542,901, with a larger sample for women (418,758) than men (124,008). Table S3a provides the detailed information on the descriptive information of the cohorts with X Chromosome data, with aggregated numbers taken from Table S3a shown here as a summary. For AFS, only data from the UK Biobank was used in the initial GWAS (leaving additional cohorts for out of sample replication), with 397,338 individuals in total, 214,547 women and 182,791 men – both for autosomal chromosomes and X chromosome. Note that the samples for women and men do not directly equate to the pooled sample size since some cohorts are family-based and only participated in the pooled analysis

**Table S3a (excerpt). Summary Sample Sizes and Descriptives, age at first birth (AFB) and age at first sexual intercourse (AFS)**

| Sample | AFB | | AFS | |
|---|---|---|---|---|
| | Autosomal Chr | X Chr | Autosomal Chr | X Chr |
| **Women** | 418,758 | 320,987 | 214,547 | 214,547 |
| **Men** | 124,008 | 110,463 | 182,791 | 182,791 |
| **Pooled** | 542,901 | 431,450 | 397,338 | 397,338 |

#### 4.2 Sample inclusion criteria

Individuals were eligible for inclusion in analyses if they met the following conditions:

**Age at first birth (AFB)**

a. They were assessed for AFB and have given birth to a child (parous); both for females and for males.
c. All relevant covariates (year of birth) are available for the individual;
d. They were successfully genotyped genome-wide (recommended individual genotyping rate > 95%);
e. They passed the cohort-specific standard quality controls, e.g., excluding individuals who are genetic outliers in the cohort.
f. They were of European ancestry.

**Age at first sexual intercourse (AFS)**

a. They were assessed for AFS and have had sexual intercourse; both for females and for males.
c. All relevant covariates (year of birth, age) are available for the individual;
d. They were successfully genotyped genome-wide (recommended individual genotyping rate > 95%);
e. They passed the cohort-specific standard quality controls, e.g., excluding individuals who are genetic outliers in the cohort.
f. They were of European ancestry.

European ancestry samples were chosen in this discovery study due to the availability of large samples[20] and for no biological or substantive reason. We acknowledge that social science research has shown large differences in the initiation of AFS and AFB by socioeconomic differences and the socially constructed category of race and ethnicity.[21,22] Socioeconomic differences are examined in this article, but the results in the current GWAS are only applicable to European Ancestry groups and need further cross-ancestry discovery.

#### 4.3 Genotyping and imputation

Table S4 provides an overview of the cohort-specific details on the genotyping platform, pre-imputation quality control filters applied to the genotype data, imputation software used, the reference used for imputation and the presence of X chromosome data. We asked cohorts to include all autosomal SNPs imputed from the 1000G panel (at a minimum) to allow analyses across different genotyping platforms. Cohorts with denser reference panels were asked to communicate this to our team. Cohorts were asked to provide unfiltered results since filters on imputed markers and so forth would be applied at the meta-analysis stage.

#### 4.4 Models used to test for association

Analysts were asked to run linear regression on AFB and a transformed AFS variable. Since age at first sexual intercourse tends to have a markedly non-normal distribution (see Fig S1), we asked the analyst for a within-sex inverse rank normal transformation before running statistical models. Analysts were asked to include birth year of the respondent (represented by birth year – 1900/10), its square and cubic to control for non-linear birth cohort effects. For those with family-based data, we suggested controlling for non-independence of family members or only include one family member in the analyses. We furthermore asked studies with family data to run a pooled GWAS on both sexes. Combined analyses that included both men and women also needed to include interactions of birth year and its polynomials with sex. In general, we asked to include top principal components to control for population stratification and cohort specific covariates if appropriate. Some cohorts only used birth year and not its polynomials because of multi-collinearity issues/convergence of the GWA analysis. Omission of these nonlinear birth year effects is unlikely to lead to biased inferences, since genotypes are not usually considered to be truly associated with birth year. However, inferences might be less accurate (i.e., have larger standard errors), since omission of nonlinear birth year effects can lead to larger residual variation.

#### 4.5 Analysis of X chromosome

The analysis on the X chromosome was completed using one of three approaches. First, XWAS software (http://keinanlab.cb.bscb.cornell.edu/content/xwas) where we suggested using the --var-het-weightcommand to generate results that are were possible to meta-analyse. Second, SNPtest, using the -method newml while this assumes complete X-inactivation (i.e. that a male with an allele is the same as a homozygous female) the effect estimates and SE approximate ½ of the corresponding betas that are produced by the XWAS software. Third, the analysis could have been performed using BOLT-LMM, performed as a separate analysis from the autosomal variants, but which includes typed autosomal variants when fitting model parameters. This later method was used for the data in UK Biobank so the AFS data was wholly derived using this method.

#### 4.6 Quality Control (QC): filters & diagnostic checks

We followed the QC protocol of the GIANT consortium[23] and employed an adapted version software package QCGWAS[18] which allows the inclusion of structural variants, in order to standardize files across cohorts and EasyQC[19] to conduct the Quality Control (QC) filtering variants and producing diagnostic graphs and statistics as described below. Where errors could not be amended by combining stringent QC with file-inspections, queries to cohorts and corrections, cohorts were excluded from the meta-analysis. See also Tables S5 (AFB) and S6 (AFS) for QC results on autosomal and Tables S7 (AFB) and S8 (AFS) for X chromosomes for AFB and AFS.

##### 4.6.1 Filters

**a) Missing data:** We filtered variants where information on both reference and other allele were missing, where the estimated effect, *p*-value, standard error, expected allele frequency or number of observations were missing.

**b) Implausible values:** We filtered variants where p-values > 1 or < 0, standard errors = 0 or = infinite, expected allele frequency > 1 or < 0, N < 0, call rate > 1 or < 0, an SE of the effect estimate which is approximately 40% greater than the expected SE based on MAF and standard deviation and for those with an $R^2$ >10% (see Winkler et al [24] for an the approximation for quantitative and Rietveld et al.[25] for quantitative and binary traits).

**c) Quality thresholds:** We filtered variants where expected allele frequency = 1 or = 0 (monomorphic variants), N < 100 to guard against spurious associations due to overfitting of the model, minor allele count <6 to guard against spurious associations with low frequency-SNPs and genotyped SNPs which were not in Hardy-Weinberg Equilibrium (HWE), with significant thresholds of threshold of $10^{-3}$ in case N < 1,000, $10^{-4}$ in case 1,000 ≤ N < 2,000, $10^{-5}$ in case 2,000≤N<10,000 and no filter in case N>10,000, imputed markers with imputation quality < 40% and SNPs with a callrate < 95%, if discrepancies between reported and expected p-value based on effect estimates and standard errors are detected.

**d) Data harmonization**: We matched the cohort based summary statistics with a 1000 Genome reference panel phase 1 version 3 reference panel provided by Winkler et al.[19] EasyQC drops mismatched of variants which cannot be solved straight away such as duplicates, allele mismatches or missing or invalid alleles. Based on graphical inspections, we applied cohort specific filters to cut drop variant with obvious deviations between expected allele frequency based on the reference panel and observed allele frequency.

##### 4.6.2 Diagnostic graphs

We produced three key diagnostic graphs for visual inspection by the two independent QC centres in Oxford and Cambridge. If problems were detected which could not be resolved by more stringent QC we had to remove the cohort from the analysis. The key diagnostic graphs depicted:

a) An **allele frequency (AF) plot** to identify errors in allele frequencies and strand orientations using the 1000 Genome phase 1 version 3 reference panel provided by Winkler et al.[19]
b) A **PZ plot** to assess the consistency of the reported p-values versus the Z score calculated based on effect sizes and standard errors.
c) A **PRS plot** of predicted versus reported standard error as developed by Winkler et al[19] and implemented by Okbay et al.[26]

##### 4.6.3 SNPs and cohorts excluded

a) Autosomal chromosomes

Overall, the quality of studies was good (for full results of the QC-filters described above see Table S5 and S6 for autosomal SNPs). Two files needed to be excluded (INGI-Carlantino for men and women – whilst pooled is in the meta-analysis) due to the filter on sample size. For autosomal chromosomes and AFB, the remaining cohorts provided 61 files, 36 for women only, 18 for men only and 7 pooled (for family data). Two studies did not provide imputation quality (KORA F3, N =1,066; and KORA F4, N =1,111) and enter the meta-analysis with only 584,866 and 496,556 SNPs respectively. For the NHS cohorts, results from our previous discovery[2] based on HapMap reference panels were reused with number of SNPs between 2,395,852 and 2,41,810. For all other cohorts, the number of variants in the analysis range between 6,583,592 for women from LBC 1921 and 16,458,651 for women in Pelotas with an average of 9,770,818. For AFS, between 16,413,259 and 16,552,240 variants from the UKBiobank have entered analysis after QC.

##### b) X chromosome

For AFB, 13 cohorts provided information on the X chromosome. Overall, we received 23 files, 13 for women, 8 for men and 2 for the pooled analysis in case there were relatives in the data. On average 275,023 variants survived QC with a minimum of 99,794 in women from WLS to 998,304 for the women in the UKBiobank sample (see Table S3b for full descriptives). For AFS, the UKBiobank provided results for between 977,536 and 990,735 variants on the X chromosome after QC between (see Table S3b; S7-8 for SNP filtering).

#### 4.7 Meta Analyses

Cohort association results (after applying the QC filters) were combined using sample-size weighted meta-analysis, implemented in METAL.[27] Sample-size weighting is based on Z-scores and can account for different phenotypic measurements among cohorts. The two QC centres agreed in using sample-size weighting to allow cohorts to introduce study-specific covariates in their cohort-level analysis. Only SNPs that were observed in at least 50% of the participants for a given phenotype-sex combination were passed to the meta-analysis. SNPs were considered genome-wide significant at *P*-values smaller than $5\times10^{-8}$ ($\alpha$ of 5%, Bonferroni-corrected for a million tests). The meta-analyses were carried out by two independent analysts in different centres in Oxford and Cambridge. Comparisons were made to ensure concordance of the identified signals between the two independent analysts. The PLINK clumping function was used to identify the most significant SNPs in associated regions (termed "lead SNPs").

We performed meta-analysis on the pooled samples for AFS and AFB and then as a separate analysis, separate meta-analyses by sex. The sex-specific results are discussed in more detail in section 5. To understand the magnitude of the estimated effects, we used an approximation method to compute unstandardized regression coefficients based on the Z-scores of METAL output obtained by sample-size-weighted meta-analysis, allele frequency and phenotype standard deviation. Further details of the approximation procedure are available in the Supplementary Information of Rietveld et al..[25] We performed conditional and joint multiple SNP analysis (COJO) to identify further independent SNPs.

#### 4.8 MTAG results

MTAG results[28] calculated from GWA meta-analysis results of the following related phenotypes: age at first birth, age at first sex, number of children ever born, childlessness, since they are all highly correlated. Using summary statistics from the pooled GWAS of each of the traits, MTAG uses bivariate score regression to account for unobserved sample overlap.

#### 4.9 Summary of discovered loci and Manhattan plots

The Manhattan plots of the pooled hits, followed by separate plots for women and men can be found in Supplementary Figures S5 for AFS and S6 for AFB by pooled (A), women (B) and men (C). A summary of the number of loci discovered is shown below, divided by autosomal and X chromosome and the pooled and sex-specific hits. The full list of association results are included in the supplementary Tables for autosomal chromosomes (S9, AFB; S10 AFS)

Summary of loci discovered

| **Phenotype** | **Autosomal chromosomes** | | | **X chromosome** | **Total** |
|---|---|---|---|---|---|
| | Pooled | Women | Men | | |
| **Age at first sex (AFS)** | 271 | 2 | 8 | 0 | **281** |
| **Age at first birth (AFB)** | 84 | 1 | 0 | 4 | **89** |
| **Total** | **355** | **3** | **8** | **4** | **370** |

Figure S5A. Manhattan plots, Age at first sex (AFS), Pooled (A), Women (B) and Men (C)

**A**

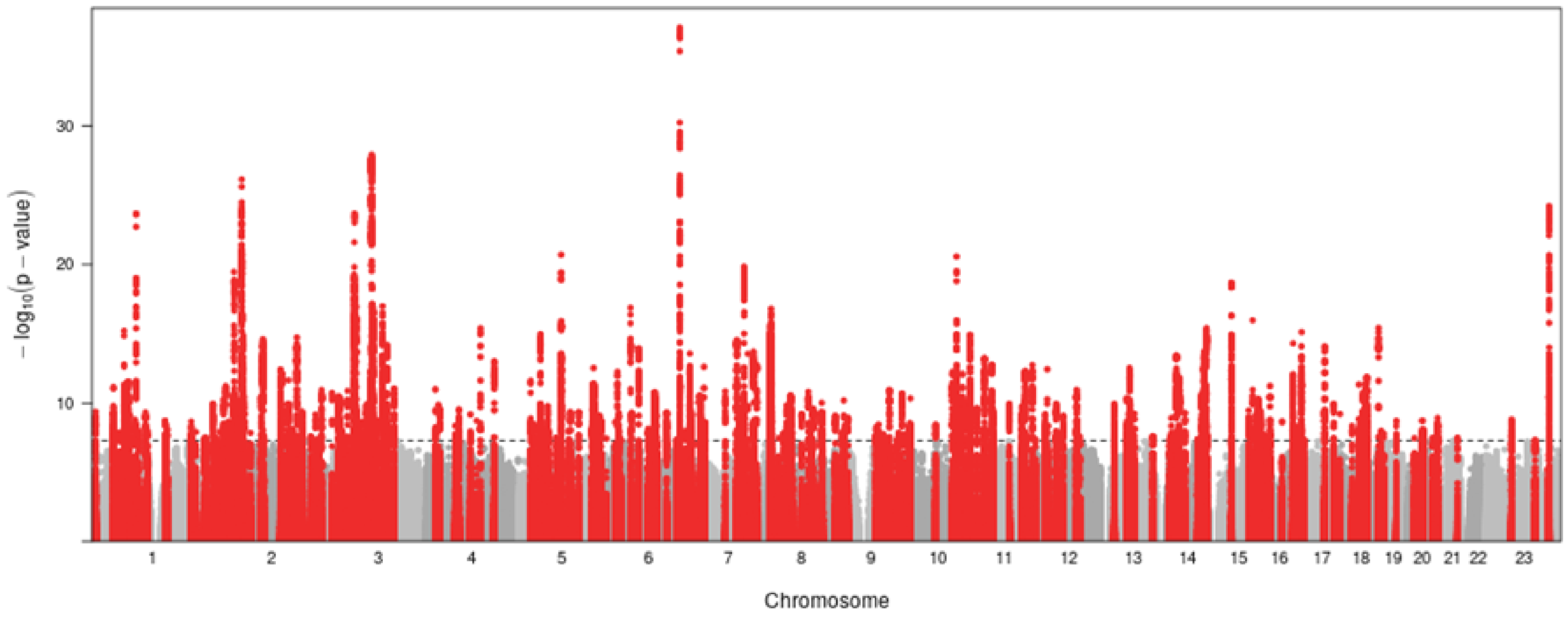

Figure S5B-C. Manhattan plots, Age at first sex (AFS), Pooled (A), Women (B) and Men (C), continued

**B**

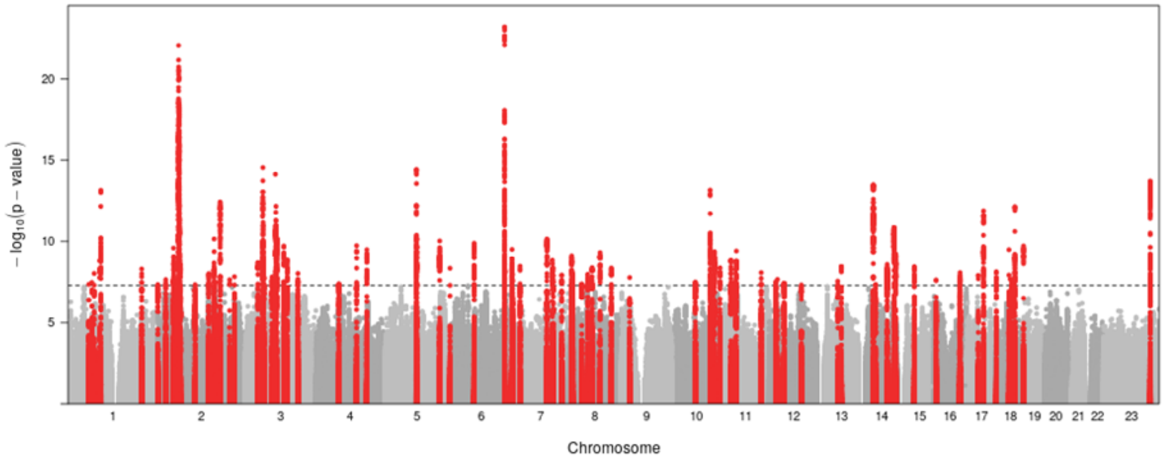

**C**

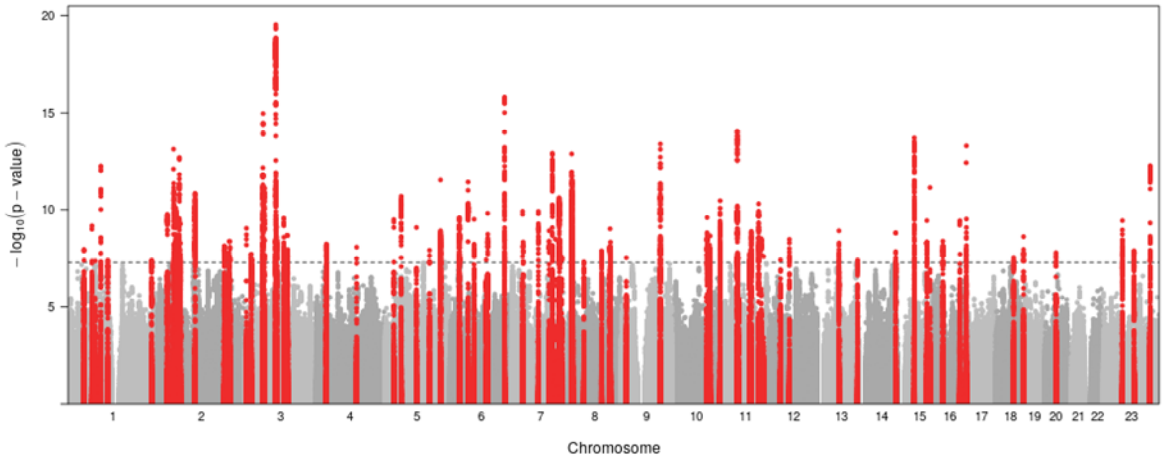

Figure S6A. Manhattan plots, Age at first birth (AFB), Pooled (A), Women (B) and Men (C)

**A**

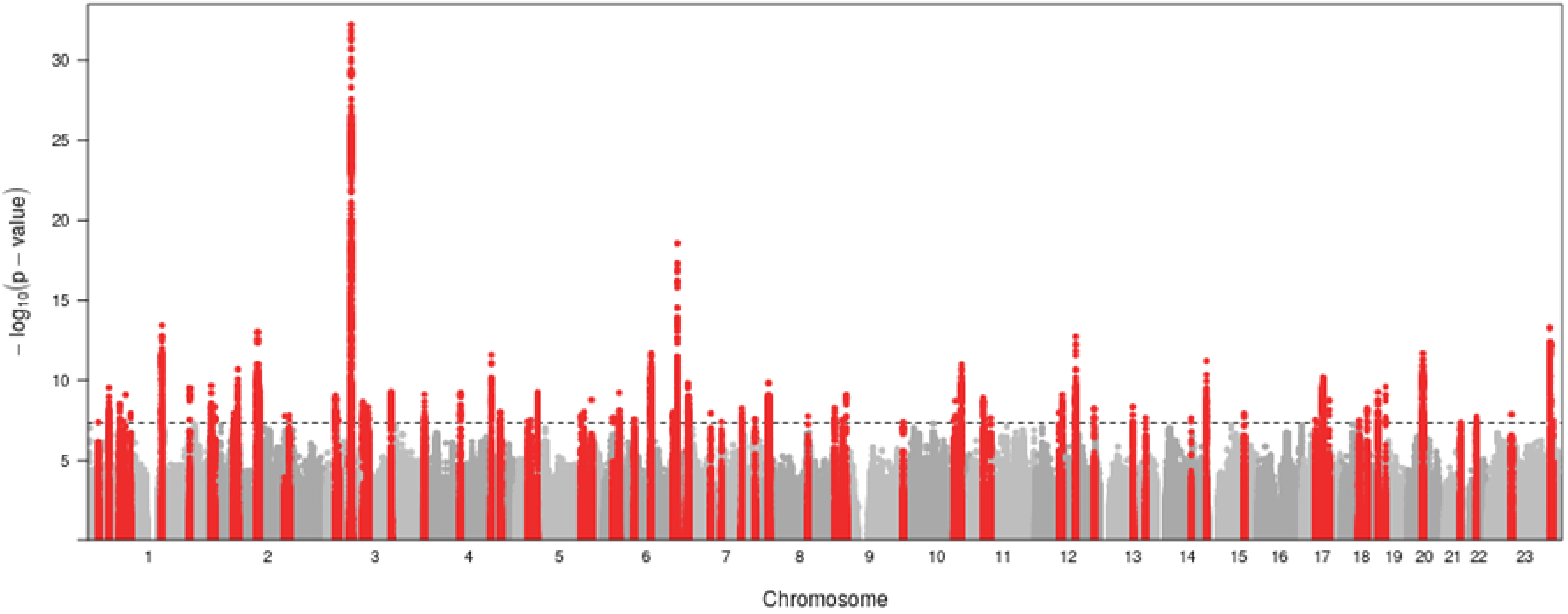

Figure S6B-C. Manhattan plots, Age at first birth (AFB), Pooled (A), Women (B) and Men (C), continued

**B**

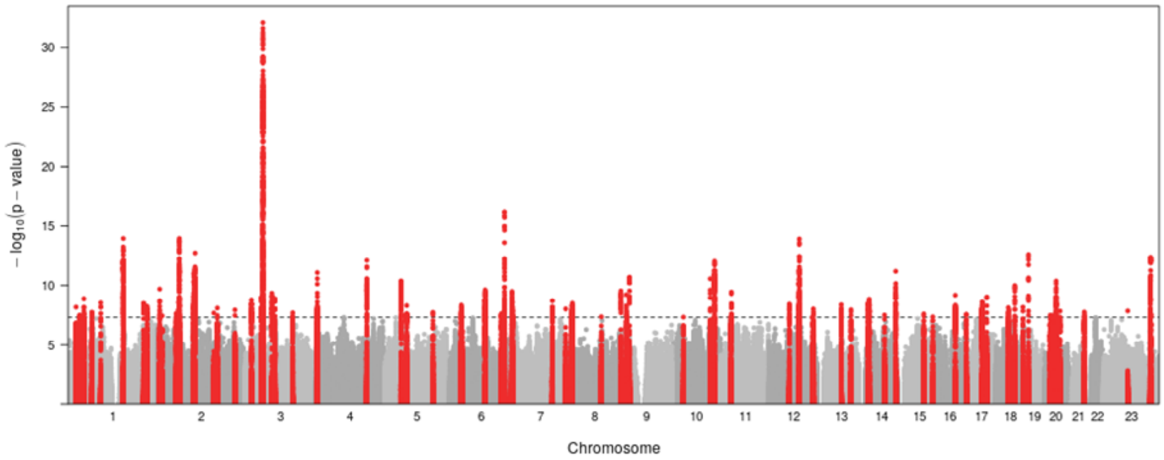

**C**

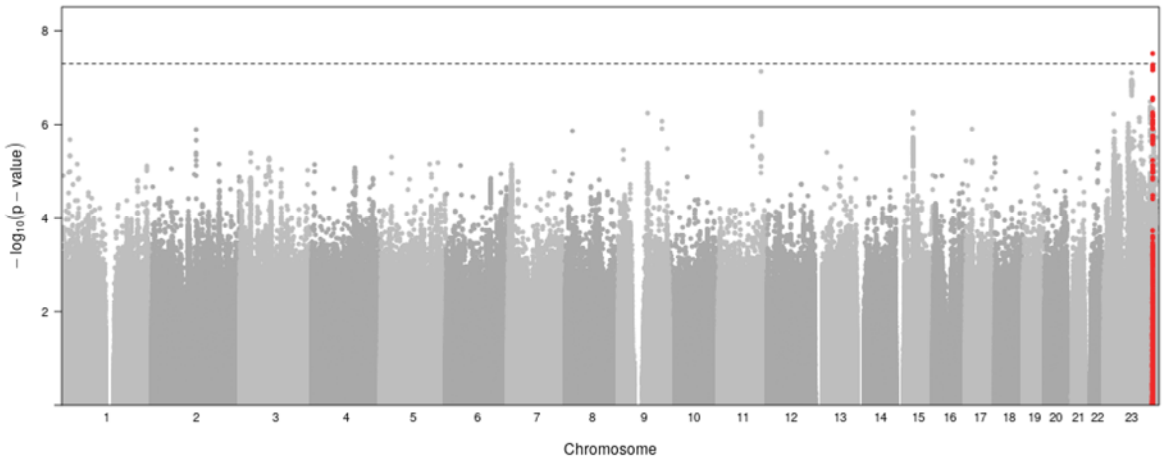

### 5. Polygenic score calculation and prediction

#### 5.1 Calculation of polygenic scores

We calculated three sets of polygenic scores:

1) **Pruning and Thresholding polygenic scores using PRSice**[29] Polygenic scores were calculated using all SNPs in the sample and with the software default values for clumping (250kb window; $r^2$=.1). Weights were based on meta-analysis results excluding the specific cohort from the calculation.

2) **LDpred polygenic scores**[30] The LD reference was calculated from the same genotyped files (AddHealth and UKHLS). We set the prior distribution for the causal fraction of SNPs equal to one. LDpred weights were then calculated under the infinitesimal model. Initial weights were based on meta-analysis results excluding the specific cohort from the calculation.

3) **MTAG+ LDpred polygenic scores.** This set of scores was calculated using the same methodology in 2), but it is based on MTAG results[28] calculated from GWA meta-analysis results of the following related phenotypes: age at first birth, age at first sex, number of children ever born, childlessness.

For both traits, we ran ordinary least-squares (OLS) regression models and report the incremental $R^2$ as a measure of goodness-of-fit of the model. Confidence intervals are based on a 1,000 bootstrap sample.

#### 5.2 Out of sample prediction

To validate the performance of the polygenic score, we performed out-of-sample prediction scores for AFB and AFS in two cohorts: the National Longitudinal Study of Adolescence to Adult Health (Add Health),[31] based in the US and the UK Household Longitudinal Study - Understanding Society (UKHLS).[32] For each out of sample calculation, we excluded the single respective cohort from the GWA meta-analysis in order to obtain independent summary statistics that have been used as weights in the calculation of the polygenic scores.[33]

The results of the polygenic score analyses are depicted in Figure S7 The proportion of variance explained by polygenic scores constructed with the MTAG+LDpred method is:

Age at First Birth (AFB): **4.80% in the UKHLS and 2.54% in the Add Health**

Age at First Sex (AFS): **5.79% in the Add Health**

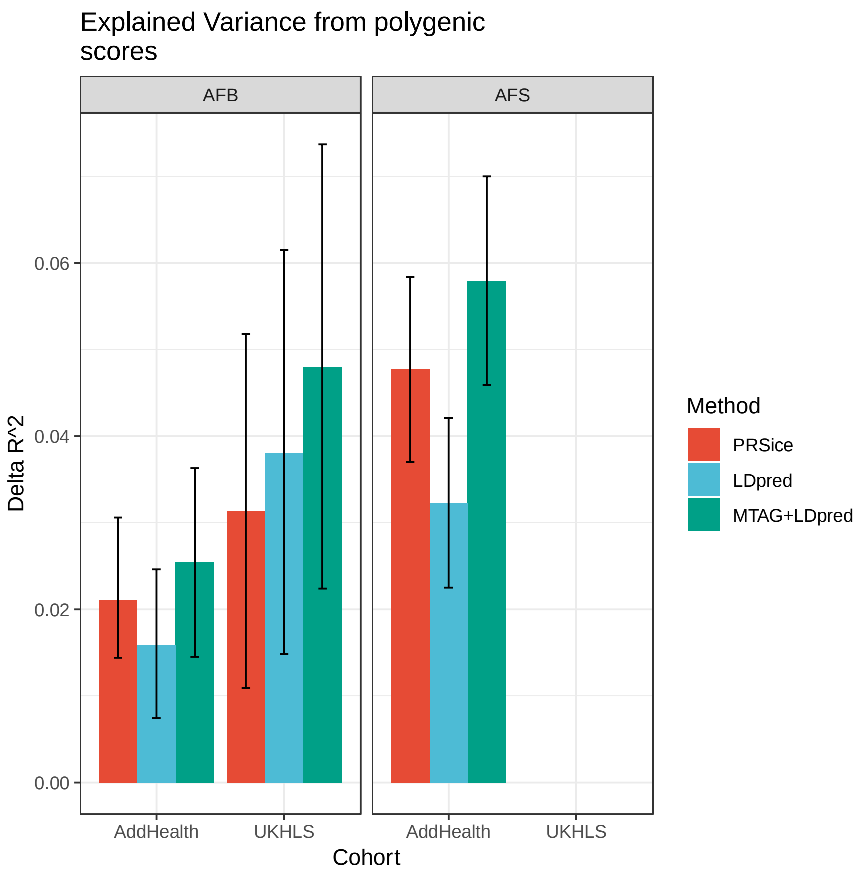

Figure S7. Variance explained from Polygenic scores for Age at First Birth and Age at First Sex using PRSice, LDPred and MTAG+LDPred in out-of-sample cohorts

Notes: The National Longitudinal Study of Adolescence to Adult Health and the UK Household Longitudinal Study

Running a linear regression predicting AFB (n=4,989) and AFS (N=9.058) in AddHealth, using the AFB and AFS PGS respectively, residualized on the first 10 genomic PCs, we can also interpret the effect in terms of time.

A 1 standard deviation (SD) change in the AFB PGS is associated with a change of: **0.48 years (SE=0.05) in AFB or around 25 weeks, which is 6.3 months.**

A 1 SD change in the AFB AFS is associated with a change of: **0.56 years (SE = 0.03) in AFS or around 29 weeks, which is 7.3 months.**

#### 5.3 Accounting for right-censoring and comparing top and bottom 5% PGS

A common limitation in the study of the genetic determinants of AFB and AFS is right censoring, which is when an individual does not experience the event of first sex or birth by the time of the interview.[34] If right censoring is not accounted for, measurements are assessed only on those who experience the event (i.e., first sex or birth of first child) before the interview date. Moreover, it does not account for the proportion of respondents that remain childless. This problem is commonly refereed in the statistical literature as “right censoring”, since the outcome is not observed for all respondents, despite the fact that part of the respondent is still “at risk” of experiencing childbirth or sexual debut. We performed additional analysis on the Add Health sample to account for this statistical issue, which is particularly pertinent given the younger age of this sample. As noted in Table S1 this cohort includes individuals born between 1974-1983 in the United States with a mean age of 28.9 (SD 1.74).

Here we report the median age for AFB and AFS, an estimator less sensitive to censoring than the mean.[34] The median AFB for men in the pooled sample is 28.42 and 26.92 for women. The median AFS for men and women is both age 17.[1]

To control for right-censored data, we first estimate nonparametric hazard functions based on Nelson-Aalen estimates (Figures S8-S9). To compare individuals with different polygenic scores, we plotted the estimated hazard of experiencing AFS and AFB for individuals at the top 5% of the polygenic score (respectively AFB and AFS) with individuals in the bottom 5% score.

Our results on age at first birth (Figure S9) shows that respondents with higher score for AFB are associated with a lower risk of childbearing at any age, with a slight increase after age 27. The results for age at first sex show that individuals with high score for AFS (i.e., having sex at later age) are less likely to have first sexual intercourse before age 19. Here sex differences appear to be very relevant. **Polygenic scores for age at first sex appear to be more relevant in explaining age at sexual debut for women than men.**

In a second part of the analysis, we estimated semi-parametric Cox regression models[35] in which we calculate the effect of genetic score on the hazard of having sex or a child for the first time, conditional on age. This class of models takes censoring into account and is widely used to study fertility timing.[34]

**The relative hazard ratio of the polygenic score for AFB is 0.85 for women and 0.91 for men**. The models show that an increase of one standard deviation in the PGS is associated with an increase of 15% in AFB for women and 10% for men. In other words, an increase in one SD of the AFB PGS relates to a 15% and 10% later age at first birth for women and men respectively.

**The relative hazard ratio of the polygenic score for AFS is 0.74 for both women and men.** This is equivalent to an increase of 26% in AFS associated with of one standard deviation in the PGS. In other words, an increase in one SD of the AFS PGS equates to a 26% postponement of age at first sexual intercourse.

[1] Note: AFS is measured in years in the Add Health Sample, while AFB is calculated in months

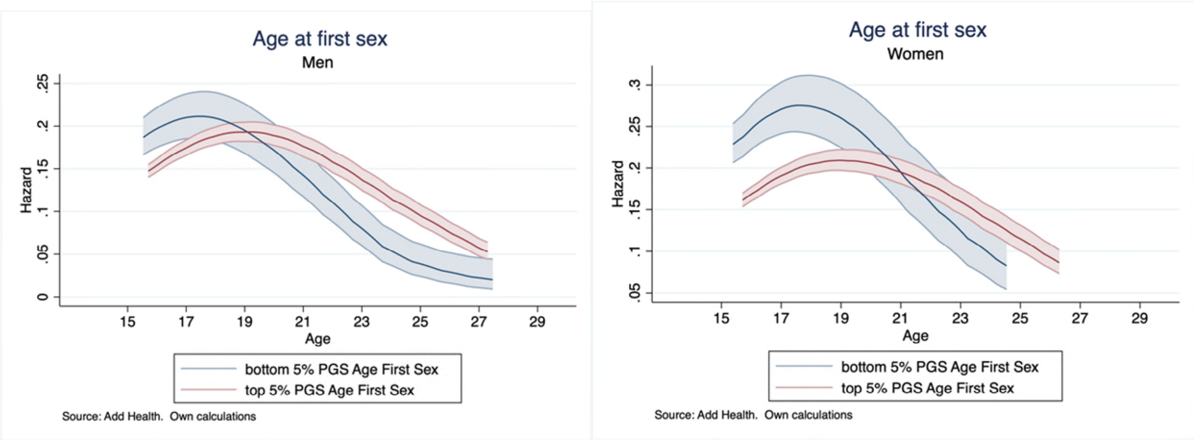

Figure S8. Nelson-Aalen hazard estimates of first sex by age. Comparison between top 5% and bottom 5% PGS of age at first sex

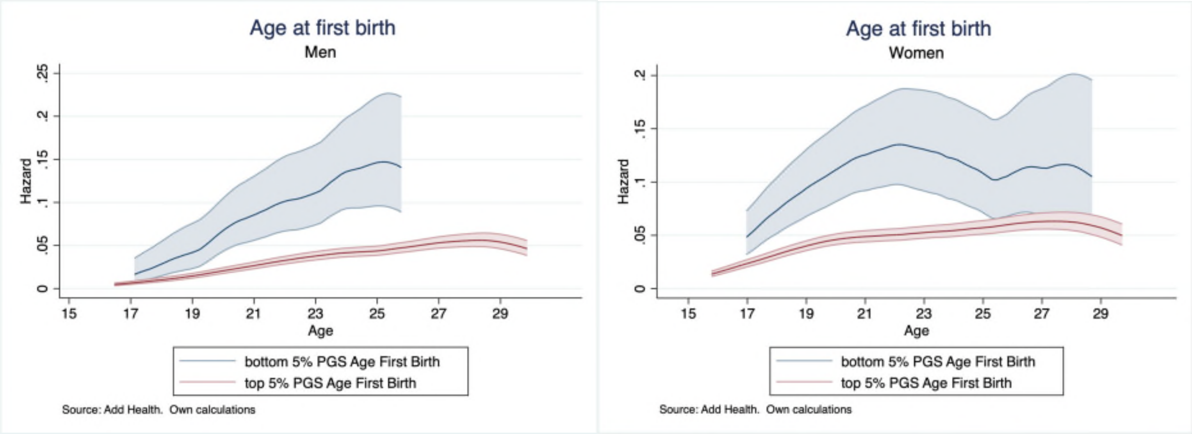

Figure S9. Nelson-Aalen hazard estimates of first birth by age. Comparison between top 5% and bottom 5% PGS of age at first birth

### 6. Testing population stratification and environmentally mediated parental genetic effects of childhood socioeconomic status

To test whether population stratification biased our results or lead to false positives, we used the LD Score intercept method described in Bulik-Sullivan et al.[36] We then examined the potential impact of environmentally mediated parental genetic effects on our PGSs, by engaging in PGS prediction across low, medium and high PGS percentiles by parent’s education, which is a proxy childhood socioeconomic status.

#### 6.1 Testing Population Stratification: LD Score intercept test

We used the LDSC software[37] to estimate LD Score regression for each of the phenotypes using the summary statistics from the meta-analyses based on all available data. For each phenotype, we used the “eur_w_ld_chr” files of LD Scores computed by Finucane et al.[38] and made available at https://data.broadinstitute.org/alkesgroup/LDSCORE/eur_w_ld_chr.tar.bz2. These LD Scores were computed with genotypes from the European-ancestry samples in the 1000 Genomes Project using only HapMap3 SNPs. Only HapMap3 SNPs with MAF > 0.01 were included in the LD Score regression.

We did not apply GC to the summary statistics we used to estimate the LD Score regression since genomic control tends to produce a downward bias of the intercept of the LD score regression.

Below a summary shows that that the LD Score intercepts are significantly but not substantially different from 1. We also see that the mean of $\chi^2$ statistics for all the SNPs in the LD Score regressions range from 1.6203 to 2.2252 for the phenotypes. Under the null hypothesis that there is no confounding bias and that the SNPs have no causal effects on the phenotypes, the mean $\chi^2$ statistics would be 1, thus mean $\chi^2$ statistics greater than 1 indicate that some SNPs are associated with the phenotypes. These estimates imply that approximately 5.5% and 5.6% of the observed inflation in the mean $\chi^2$ statistics for AFB and AFS, respectively, is accounted for by confounding bias (due to population stratification or other confounds), rather than a polygenic signal. This is calculated from the Standard Errors using the 95% Confidence Intervals. These estimates may also be inflated by model misspecification or LD score mismatching.

**Summary of LD score intercept, SE and mean $\chi^2$statistics for AFB and AFS**

| | Age at first birth (AFB) | Age at first sex (AFS) |
|---|---|---|
| **LD score intercept** | 1.0341 (0.0089) | 1.0687 (0.0132) |
| **Lambda GC** | 1.471 | 1.8419 |
| **Mean $\chi^2$** | 1.6203 | 2.2252 |
| **Ratio** | 0.0549 (0.0143) | 0.0561 (0.0107) |

#### 6.2 Polygenic score prediction by childhood socio-economic status

To explore the impact of environmentally moderated parental genetic effects on our PGSs, we examined PGS prediction across low (0-10%), medium (50-60%) and high (90-100%) PGS percentiles by parent’s education, which is a proxy childhood socioeconomic status, using AddHealth. This also follows recent research that has demonstrated that PGSs differ in their predictive accuracy and ability not only by ancestry, but also by factors such as the socioeconomic environment within ancestry groups.[39] Figures S10A-B show the PGS score prediction for AFS (A) and AFB (B) divided into three percentile groups by parental level of education (college versus no college). Here we see variations in prediction by PGS percentile and parental education. Figure S10A shows that those who are in the 90-100$^{th}$ PGS percentile for later AFB indeed postpone first childbirths particularly past the ages of 27. This PGS is accentuated for those in the highest (90-100%) and moderate (50-60%) percentile groups particularly for those who have high educated parents.

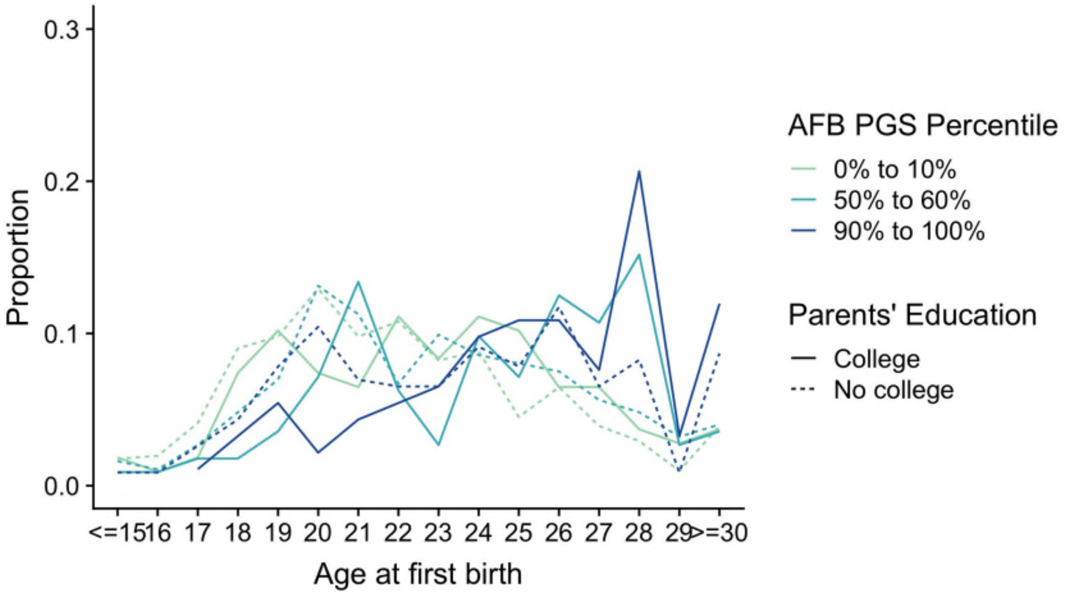

Figure S10A. AFB PGS score by percentile groups and parent’s educational level

Figure S10B shows that those in the highest AFS PGS percentile (90-100%), which would predict later age at first sexual debut and have higher educated parents have both lower sexual initiation but also are the group from age 18 and above who have a systematically later age at first sexual intercourse. This suggests that even within these groups there is also considerable heterogeneity that might be related to higher or lower behavioural disinhibition and externalizing behaviour regardless of the parental environment.

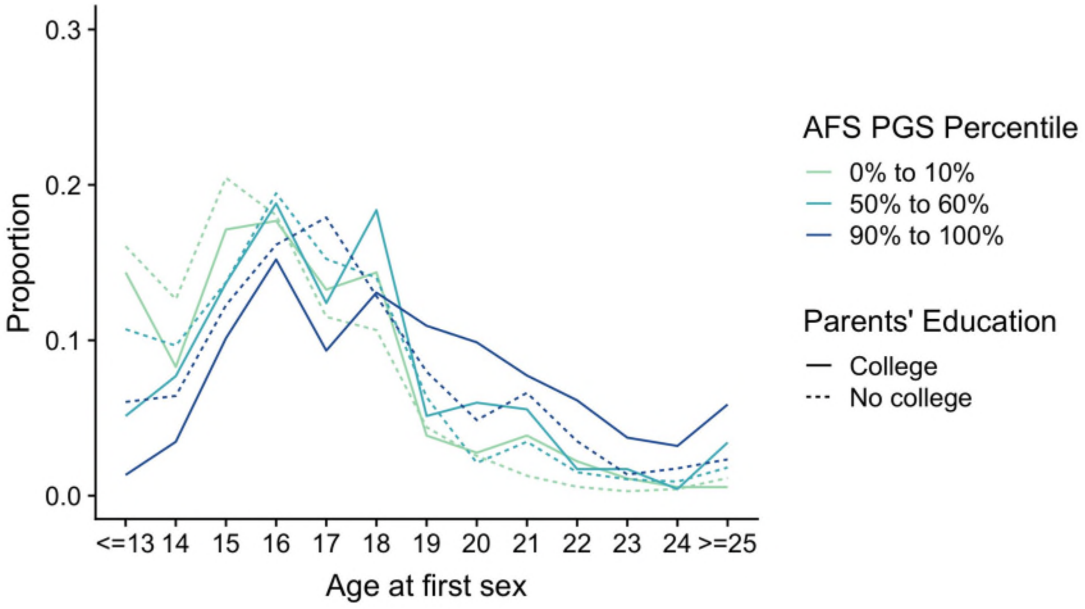

Figure S10B. AFS PGS score by percentile groups and parent’s educational level

### 7. Genetic correlations with related traits

In order to understand the genetic relationships among and between our fertility phenotypes and potentially related phenotypes, we calculated their SNP heritabilities and their genetic correlations (Figure S11). The estimates represent the genetic correlation between the two traits using all polygenic effects captured by the SNPs, based on the LD-score regression method developed by Bulik-Sullivan et al.[37] We used summary statistics and the 1000 Genomes reference set and restricted the analysis to European populations. They estimate a SNP's LD score, which measures the amount of genetic variation tagged by a SNP. We also follow the common convention of restricting our analyses to SNPs with MAF >0.01, thus ensuring that all analyses are performed using a set of SNPs that are imputed with reasonable accuracy across all cohorts. The standard errors (SEs) are produced by the LDSC python software package that uses a block jackknife over the SNPs.

#### 7.1 Genetic correlation with related traits

In the main body of the paper, we show the genetic correlation with 25 traits. Here we show additional results and discussion of the estimated the genetic overlap or correlation between different traits, first pooled by both sexes and also divided by sex. Traits are divided into 6 different categories of:

- **reproductive** (age at menarche,[40] age at menopause,[41] age at voice breaking,[42] number of children ever born (by sex),[43] number of sexual partners,[44] age started oral contraceptives[45], breast cancer[46] but also specifically those related to infertility, endometriosis and severe endometriosis[47]),
- **behavioural** (years of education,[48] cognitive performance,[48] subjective well-being,[49] risk tolerance in adulthood[44]),
- **psychiatric disorders** (attention deficit hyperactivity disorder (ADHD), [50,51] schizophrenia (SCZ),[52] bipolar disorder,[53] major depressive disorder (MDD)[54] and anorexia[55]),
- **substance use disorders** (alcoholic drinks per week (DPW),[56] age at initiation smoking (SI),[56] cannabis use[56], cigarettes per day (CPD)[56], smoking cessation (SC)[56]),
- **personality** (neuroticism,[28] openness to experience,[57] loneliness[58]),
- **anthropometric** (height,[59] waist to hip ratio for BMI (by sex),[60] and BMI (by sex)[61])

Note that we include the most recent GWAS results when they are openly shared and available. For example, a 2018 study of intelligence[62] was updated by later by another study of cognitive performance,[48] in which we use the most recent. From previous research we know that the onset of reproductive behaviour phenotypes are correlated with reproductive, behavioural (e.g., educational attainment), personality and anthropometric traits.[2] Age at menarche, menopause and age at voice breaking was included as a fundamental aspect of reproductive biology and breast cancer was included because of evidence linking it to age at menarche.[63] A negative phenotypic and genotypic relationship with later AFB and lower NEB has been well-established in previous studies.[6,64] Number of sexual partners and age at starting oral contraception can be a reflection of biological development, but also risk-taking behaviour. Previous research has also linked onset of reproductive behaviour to achieving higher levels of education and cognitive performance, linking it to later sexual debut[65] and fertility postponement.[64,66]

Twin research in behaviour genetics has consistently shown a link between behavioural disinhibition and externalizing behaviour with psychiatric disorders and substance use disorders,[67] yet with the exception of some early studies,[68] few have linked it to early sexual and reproductive behaviour.[69] Given our focus on the timing of events – namely early sexual debut and teenage pregnancy – we also wanted to test whether there was an underlying genetic propensity related to early reproductive behaviour onset with behavioural disinhibition, externalizing behaviour and risk taking. We therefore extended the analysis to examine psychiatric and substance use disorders and personality. Previous phenotypic studies have shown that AFB is negatively correlated with neuroticism.[70] Others have shown a U-shaped genetic relationship between schizophrenia and AFB.[71] Finally, previous work has also linked reproductive success to anthropometric traits such as height, BMI and waist-hip ratio.[72,73]

#### 7.2 Genetic correlations by sex

We examined whether patterns of genetic correlation varied across the sexes (Figure 2, Main Paper). We also examined whether patterns of genetic correlation varied by birth cohort but found little to no evidence for variation across cohorts (results available upon request). A summary Table containing these results is in Table S11.

**Reproductive traits.** Age at menarche for girls and age at voice breaking for boys marks the start of the reproductive career and adolescent development.[63] Studies have shown that earlier menarche is related to great sexual risk taking as an evolutionary reproductive strategy, particularly in harsh family environments.[74] Whereas variation in age at menarche is often more related to living conditions and nutritional status, age at menopause appears to be mainly influenced by biological factors and primarily the reproductive history of individuals.[75] Numerous demographic studies have shown that a later AFB is linked with a lower number of children due to voluntary desires for fewer children and unintended childlessness due to fecundity and infertility problems.[6] To capture infertility and fertility problems, we also included miscarriage or stillbirth. A higher number of sexual partners has been phenotypically correlated with a cluster of adolescent risk behaviours such as unintended pregnancy and substance use.[76] Finally, we included the age at the start of oral contraception which is a unique marker that proxies biological development and risk aversion to pregnancy.[77]

We find strong correlations across virtually all of the reproductive traits. There is a positive correlation with later AFS and AFB with later age at menarche, menopause and voice breaking (around 0.12 to 0.27). A later age at menarche has been previously associated with subfecundity, diminished ovarian function and infertility.[78] We also find a negative genetic correlation with number of children ever born (NEB), considerably stronger for males (AFB males –0.87 ±0.09; AFB females, –0.67 ±0.02) and also with miscarriage or stillbirth (AFB females, –0.51 ±0.06; AFS females, –0.67 ±0.06). In other words, later AFB and AFS are genetically correlated with a lower NEB. We see that the genetic propensity for later AFS and AFB are rather negatively correlated with ever experiencing a miscarriage or stillbirth. Although there is some relationship, we remain cautious regarding the estimates of endometriosis considering the smaller sample size and larger CIs of our estimates. There was also a striking positive correlation between the age at starting oral contraceptives (AFB females, 0.76 ±0.03; AFS females, 0.88 ±0.06), suggesting later sexual and reproductive onset was also linked with later age at using contraceptives, which may serve as a marker for development. Number of sexual partners was negatively correlated with later AFS/AFB and stronger for the related sexual behaviour trait of AFS (AFS males –0.57 ±0.02; AFS females, –0.59 ±0.02), than AFB (AFB males –0.25 ±0.06; AFB females, –0.25 ±0.02).

**Behavioural traits.** Some of the strongest correlations are with behavioural traits, particularly educational attainment for women, with a robust overlap of AFB (0.74 ±0.01) compared to AFS (0.53 ±0.01), also noted in previous studies.[2,9] The strong relationship with AFB and education is not surprising since phenotypically, higher educational attainment is associated with later AFB in most advanced societies.[6] Others previously found a relationship of initiation of sexual activity and fertility with higher cognitive ability,[66] with additional research required to separate whether these cognitive scores are (very) likely confounded by socio-economic status and environment. Others have found a relationship between higher cognitive ability and later age at first sexual intercourse.[65] Here we also see a negative genetic correlation between adult risk tolerance and later AFS/AFB (AFB females, – 0.25 ±0.03; AFB males, – 0.29 ±0.07; AFS females, –0.40 ±0.03; AFS males, –0.40 ±0.02) or other words, those less genetically prone to risk are also less prone to early teenage sex and teenage pregnancies. Conversely, if the variables were reverse coded it would mean that those more genetically prone to risky behaviour in adulthood are also prone to earlier sexual debut and earlier births.

**Psychiatric disorders.** One of strongest genetic associations in our analysis is with ADHD (AFB females, –0.63 ±0.03; AFB males, – 0.68 ±0.09; AFS females, –0.58 ±0.03; AFS males, –0.61 ±0.03), and to some extent also Major Depressive Disorder (MDD) (AFB females, –0.42 ±0.03; AFB males, – 0.33 ±0.08; AFS females, –0.37 ±0.03; AFS males, –0.32 ±0.03). ADHD has been phenotypically related to elevated risky sexual behaviour, often comorbid with problematic substance use problems such as alcohol use, smoking and cannibus.[79] The mechanism is linked to higher levels of behavioural disinhibition, externalizing and hyperactive and impulsive behaviours. Interestingly, the internalizing psychiatric disorder of anorexia, is positively genetically correlated with postponement of early sex and births. This disorder is often related to exaggerated cognitive control, rigid behaviour and impaired ability to be flexible or impulsive.[80] In other words, postponement of early sex and childbirth appears to be related to the opposite spectrum of behavioural disinhibition and externalizing of ADHD and early sex and teenage pregnancy.

**Substance use**. Addictive and substance use also had striking correlations, particularly with age at onset of smoking (AFB females, 0.73 ±0.03; AFB males, 0.74 ±0.07; AFS females, 0.67 ±0.03; AFS males, 0.68 ±0.03), which provides a rare genetic marker of a window into adolescence and timing of early sexual behaviour and pregnancies. Related to this is a negative correlation of ever engaging in cannabis use (AFB females, –0.25 ±0.05; AFB males, – 0.22 ±0.13; AFS females, –0.43 ±0.06; AFS males, –0.43 ±0.06). Earlier smoking may capture an underlying propensity for a variety of behavioural disinhibition and externalizing behaviours in adolescence. There is also an established link between smoking with a longer time to conception and decreased fertility.[81] Smoking has been linked to problems with preimplantation, shrinking size and quality of oocytes and decreased sperm motility in men.[82,83] Another plausible mechanism is that an earlier age at smoking is linked to a lower socioeconomic status, linked to multiple environmental risk factors and a higher co-morbidity of related diseases.[84] Smoking often serves as a strong marker for structural and resource disadvantage.

**Personality traits.** For these traits, the most striking finding is the relationship with openness to experience and later AFS/AFB, particularly with men. We also see a positive correlation with late AFS and AFB and loneliness (AFB females, 0.40 ±0.03; AFB males, 0.24 ±0.07; AFS females, 0.37 ±0.03; AFS males, 0.31 ±0.03) and to some extent a negative correlation with neuroticism in females (AFB females, –0.21 ±0.05), which may be related to the ability to find a partner.[85]

**Anthropometric traits.** With the exception of BMI, we find no striking results in relation to anthropometric variables. BMI is related to pubertal development but a very low and very high BMI is found in delay the timing and number of children.[86]

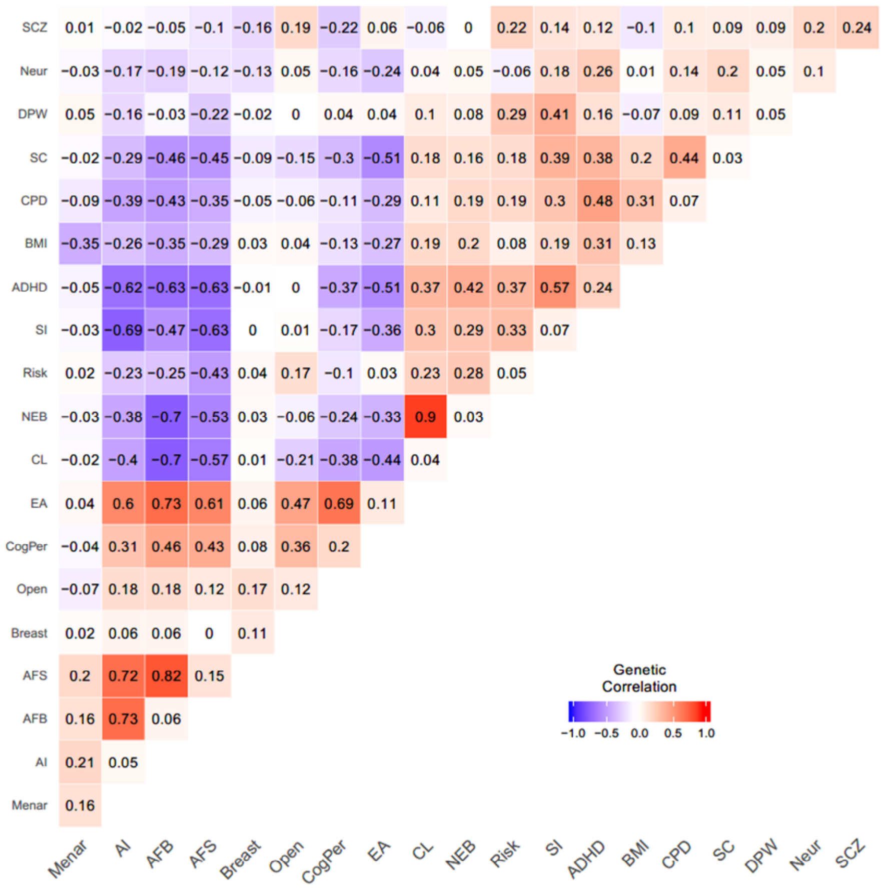

Figure S11. Genetic correlations and SNP heritabilities between and among reproductive, behavioural, psychiatric, substance use, personality and anthropometric traits

Note: Calculated by LD score regression, with SNP heritabilities along the diagonal. Menarche = Age at menarche; AI = age at initiation of smoking, AFB = age at first birth; AFS = age at first sex; Breast = ever had breast cancer; Open = openness (personality); CogPer = cognitive performance; EA = educational attainment in years; CL = childlessness; NEB = number of children ever born; SI = ever smoked; ADHD = attention deficit hyperactivity disorder; BMI = Body mass index; CPD = cigarettes per day; SC = smoking cessation; DPW = alcoholic drinks per week; Neur = neuroticism (personality); SCZ = Schizophrenia.

We recognize that these are only correlations and it is likely there are also pleiotropic variants with multiple biological effects or other factors such as some traits that are mediated by environmental influences. For this reason we explore additional analyses in an attempt to understand the underlying etiology and causal relationships between these traits.

### 8. Uncovering shared genetic etiology with Genomic SEM

In an attempt to understand the etiology the correlations described in the previous section, we used the R package GenomicSEM[87] to fit multivariate genetic regression models. GenomicSEM uses structural equation modelling to decompose the genetic covariance matrix, calculated using multivariate LD score regression, of a set of traits. The user specifies a model, the parameters of which are estimated by minimizing the difference between the observed genetic covariance matrix and the covariance matrix derived from the model. Formally, structural equation models subsume many statistical methods and are quite flexible. One model that can be fit using GenomicSEM is the multivariate genetic regression model. In this model, some trait C is regressed on traits A and B, producing estimates of the genetic correlation of A with C, independent of B, and of B with C, independent of A. We note that this model is equivalent to a simple mediation model, with C as the dependent variable and either B or C as the mediator.

#### 8.1 AFB and AFS regression educational attainment (EA) and trait X

We fit a series of such models in which AFB was regressed on EA and a trait X (Table S12A-SL and Figure S12A). AFB was chosen as the dependent outcome become an individual's first birth most often occurs after they have completed their education. We also fit an analogous series of models in which AFS was regressed on EA (Table S11A-L and Figure 12B), which showed similar patterns of conditional association.

In each case, the conditional association of EA and AFB remained substantial, suggesting that the genetic association of EA with AFB is largely independent of the genetic components of personality (as measured by openness, neuroticism, and risk tolerance), BMI, loneliness, MDD, cognitive performance, substance use, and sexual behaviour (as measured by number of sexual partners). Additionally, the conditional association of cognitive performance with AFB was close to zero, suggesting that cognitive performance does not influence AFB above and beyond its effect on EA. These observations support the conclusion that the genetic correlation between EA and AFB is mediated by environmental mechanisms—those who have high educational attainment have been exposed to an environment that encourages later childbirth.

We note, however, that the conditional association of EA and AFB was smallest in the model of age of initiation of smoking (AI). AI is partially genetically distinct from other aspects of cigarette smoking and captures, in part, risk tolerance in adolescence, since most regular smokers initiate in adolescence.[88] AI, then, might capture an aspect of adolescent risky behaviour that our measure of risk tolerance (taken in middle age) does not, explaining its apparent mediation of the relationship between AFB, AFS, and EA.

In order to explore the genetic relationship between reproductive biology and our phenotypes of interest, we obtained results from a GWAS of sex hormone levels (Figure S13) and fit sex-specific genetic multivariable regression models for AFB (Tables S11E and F) and AFS (Tables E and F). We observe no significant moderation of the association of these variables with EA. In summary, a wide range of variables are unable to explain the association between AFB and AFS and EA.

We noted substantial genetic correlations between AFB and AFS (rg = 0.82, SE = 0.03) and between AFS and EA (rg = 0.61, SE = 0.02), paralleling the correlation between EA and AFB. We fit a genetic multivariable regression model in which EA was regressed on AFB and AFS and found a substantial conditional standardized association of EA and AFB (beta = 0.70, SE = 0.05) but a small conditional standardized association of EA and AFS (beta = 0.04, SE = 0.05), as expected since AFS occurs before AFB.

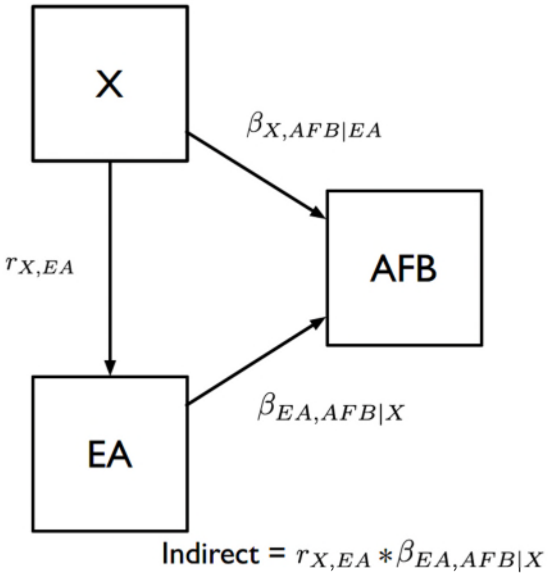

Figure S12A. A path diagram showing the structure of the genetic multiple regression model fit to EA and AFB

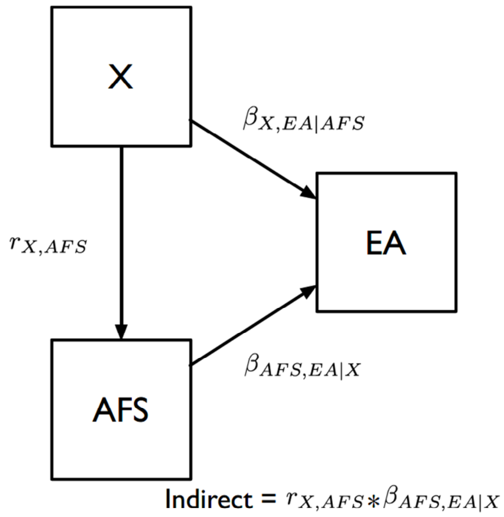

Figure S12B. A path diagram showing the structure of the genetic multiple regression model fit to AFS and EA

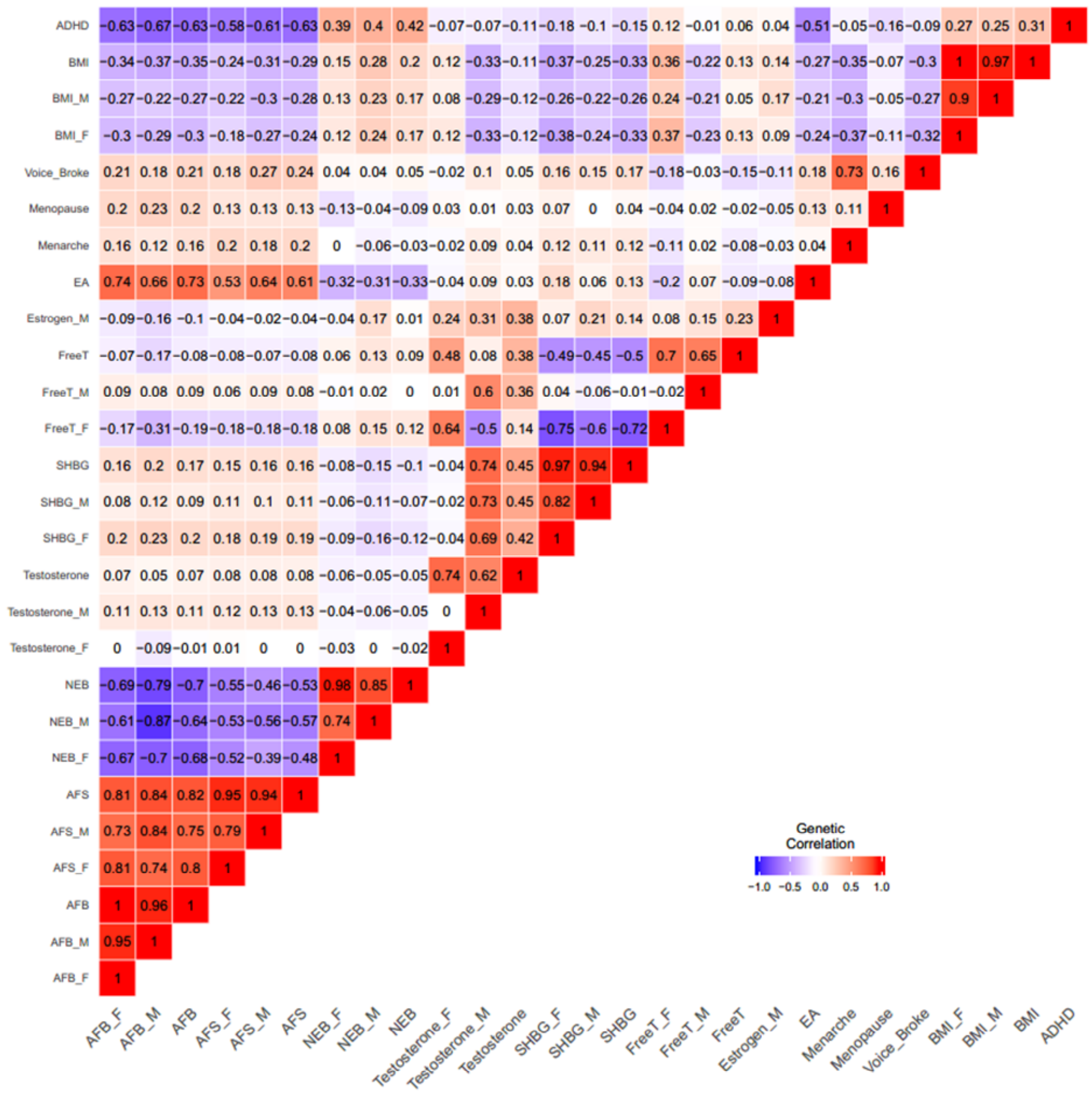

Figure S13. A heat map showing the genetic correlations between and among the fertility GWAS phenotypes, the sex hormone phenotypes, and other phenotypes related to reproductive biology, as calculated by LD score regression.

Notes: BMI = Body mass index; VoiceBroke = Age voice broke; Menopause = Age at menopause; Menarche = Age at menarche; FreeT = Free testosterone; SHBG = Sex hormone-binding globulin; NEB = Number ever born

#### 8.2 Reproductive biology and externalizing behaviour explanation of variance

From our analyses, it emerged that the timing of the onset of reproductive behaviour appears to be driven by both reproductive biology and externalizing behaviour.[64] To test the potential amount of variance that each explained we engaged in Exploratory Factor Analysis (EFA) and an additional Genomic SEM.

First, we used exploratory factor analysis (EFA), which is a means of studying the relationships between a set of variables. This method was used to examine whether the genetic signal of the onset of reproductive behaviour originated from two genetically distinguishable subclusters of a reproductive biology component and an externalizing behaviour component. To engage in a simple test of this theory, we used proxies of these categories, namely, age at menarche and risk tolerance. To test this theory we fit a two factor EFA model to the genetic covariance matrix of AFB, AFS, NEB, risk tolerance, and age at menarche. The model accounted for 47% of the overall variance but only 22% of the variance attributed to risk tolerance and 4% of the variance to age at menarche, indicating the genetic relationships between these variables are partially but not fully captured by a two-factor model.

To test this further we then focussed on more robust and additional measures of reproductive biology and externalizing behaviour and engaged in a sex-specific analysis of AFB for women. In order to parse out the genetic influences on age at first birth, we fit a genomic structural equation model (Genomic SEM) where AFB in women is regressed on age at menopause, age at menarche, and a latent factor representing the common genetic tendency to externalizing behaviour (Figure S14). The factor is measured by AFS in women, age at initiation of smoking, age first used oral contraception, and ADHD, with the model scaled to unit variance for the latent factor. The fitted model had a CFI equal to 0.95 and an SRMR equal to 0.09, suggesting a reasonable fit.

The standardized residual variance for AFB in the model is 0.12 (SE = 0.04), indicating that most of AFB's SNP heritability can be accounted for by age at menopause, age at menarche, and our externalizing factor, with all three variables having a statistically significant independent effect, but the externalizing factor showing the strongest association by far. We can conclude that the genetic traits we include predict 88% of the genetic variance for women's age at first birth and that externalizing behaviour explains most of the common genetic variance of AFB in women, in the contexts measured in our study. This is of course considering the standard caveats for LDSC genetic variance and covariance estimates and that we include common variants only, and examine this using information from selective European Ancestry populations.[89,90] We note also that selection bias, induced by the fact that AFB can only be measured among individuals with at least one live birth, may have inflated this estimate.

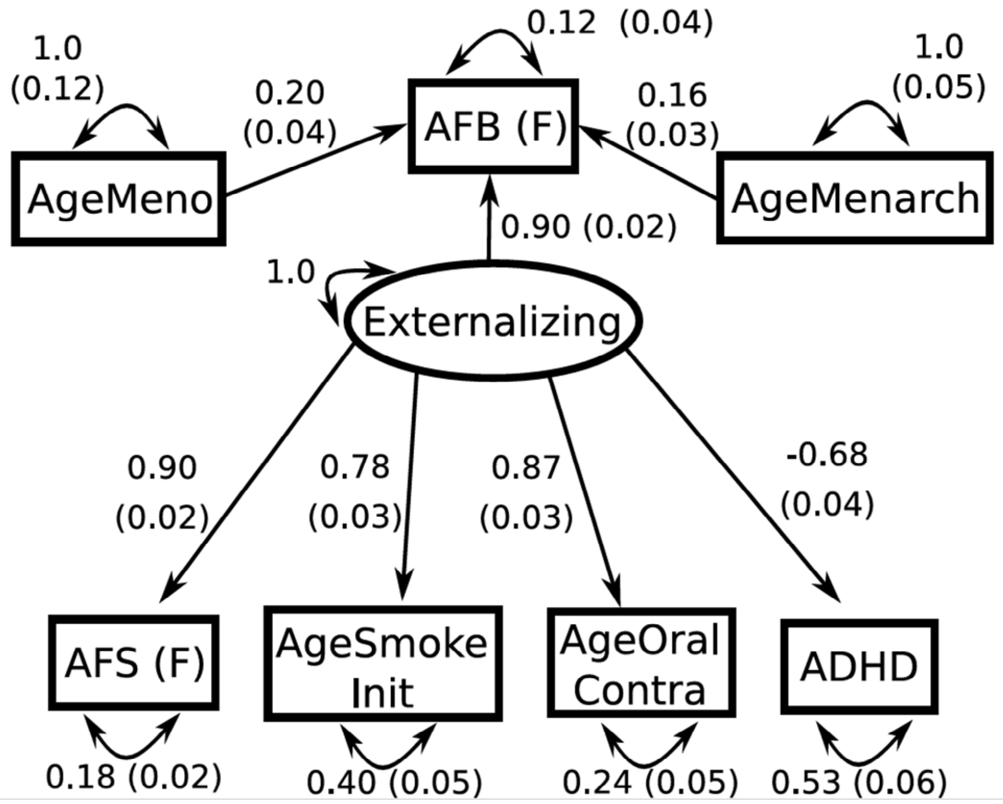

Figure S14. A path diagram for a Genomic SEM model of the relative associations of an externalizing latent factor, age at menopause, and age at menarche with age at first birth in women

Standardized parameter estimates are shown with standard errors in parentheses. AFS (F) = Age at first sexual intercourse in women; AFB (F) = Age at first birth in women; AgeSmokeInit = Age of smoking initiation; AgeOralContra = Age first used oral contraception; ADHD = Attention-deficit hyperactivity disorder; AgeMenarch = Age at menarche; AgeMeno = Age at menopause

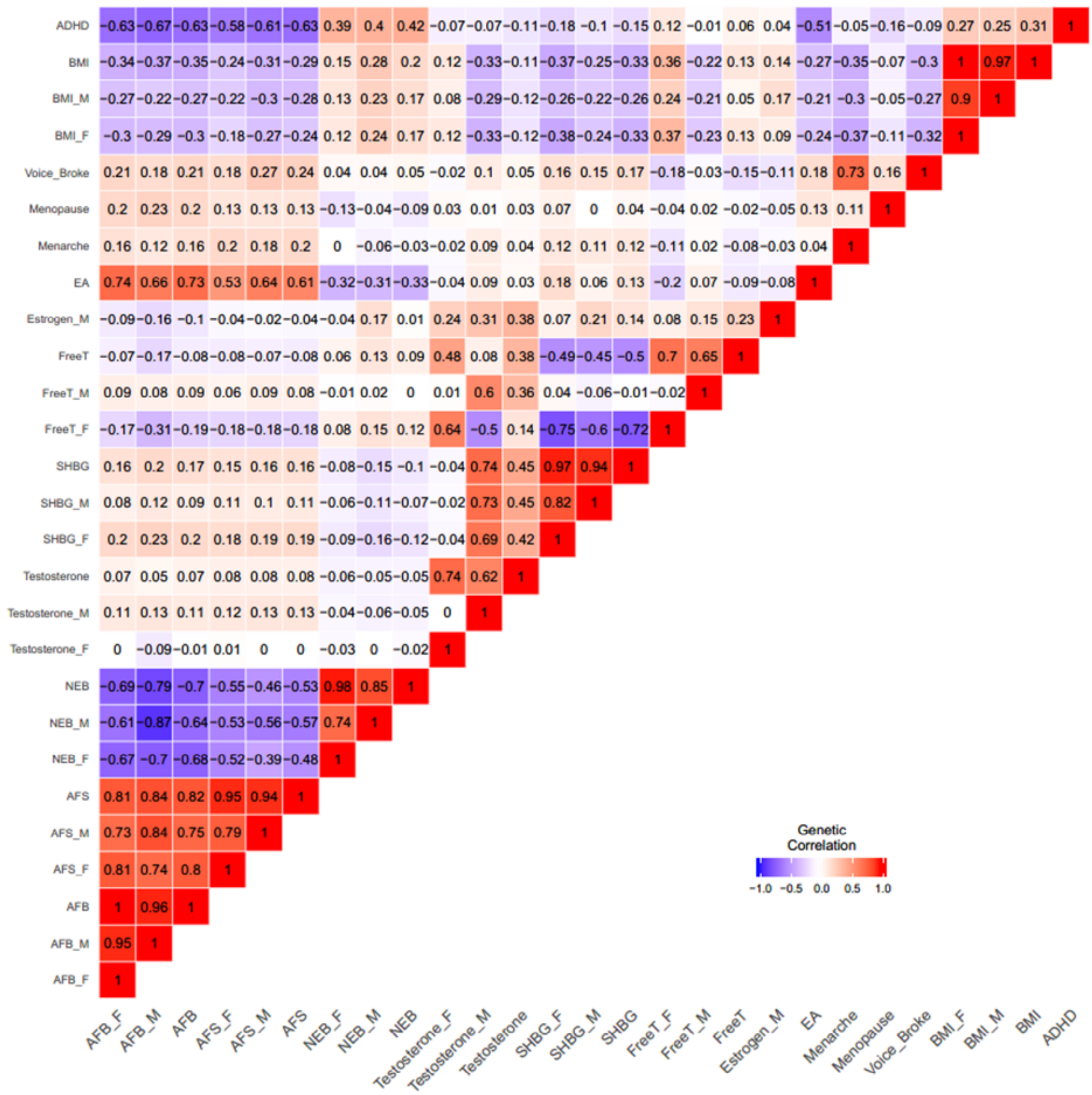

Figure S15. A heat map showing the genetic correlations between and among the fertility GWAS phenotypes, the sex hormone phenotypes, and other phenotypes related to reproductive biology, as calculated by LD score regression.

Notes: BMI = Body mass index; VoiceBroke = Age voice broke; Menopause = Age at menopause; Menarche = Age at menarche; FreeT = Free testosterone; SHBG = Sex hormone-binding globulin; NEB = Number ever born

### 9. Bi-directional MR of reproductive behaviour, teenage behavioural disinhibition and onset of later life disease

#### 9.1 Background, methods and innovation

Previous studies have suggested a link between reproductive behaviour and health outcomes.[91–94] The previous analyses showed a considerable overlap between the genetic loci identified for reproductive behaviours and educational attainment. Using GenomicSEM we aimed to capture the unmeasured factor and etiology underlying teenage behavioural disinhibition and externalizing behaviour which we investigated using traits such as age at smoking initiation, personality traits and adult risk taking. It is plausible, however, that the casual pathways connecting these phenotypes are potentially bidirectional and that each of our measured phenotypes might offer distinct contributions. We then tested whether causal pathways linking these phenotypes are potentially bidirectional and whether our phenotypes might offer distinct contributions.

We identified 1000 Genomes proxies for our SNPs and used these in multivariate Mendelian Randomisation (MR) models. First, we modelled the interplay between AFB, AFS and EA (educational attainment)[95] as well as risk taking (measured in adulthood)[44] and age at smoking initiation (AI).[56] In each case IVW[96] and MR-EGGER[97] methods were performed, with an additional round of IVW performed once a Steiger filter[98] had been applied to remove SNPs that appears to show a primary association with the outcome rather than the exposure. Multivariate MR was use to try to dissect causal pathways.[99]

A second set of MR analyses focused on links to late life diseases, namely type 2 diabetes (T2D)[100] and coronary artery disease (CAD)[101], using the same methods. T2D and CAD were chosen since they are common diseases with a strong behavioural component. In particular, we use multivariate methods to test whether AFS or AFB had independent effects once the well-established links to length of educational attainment were controlled for. The model shows the effects of variants discovered for AFB and AFS and education on two key later life diseases, controlling for the alternative pathways represented by each phenotype. These analysis were also performed in a sex-specific manner using the available outcome data for men and women separately for type 2 diabetes.

#### 9.2 Results MR

The bidirectional analysis, suggested a complex causal web where each of the assessed phenotypes appear to have an important impacts on the others (Table S13A). The one exception was behavioural disinhibition during teenage years proxied by age at initiation of smoking (AI). This suggests that the specific timing of the onset of behavioural disinhibition may be crucial. Both AFS, and age at smoking initiation are phenotypes that represent behavioural disinhibition and externalizing behaviour in precisely in the window of adolescence and early adulthood. The links between both AFS and AFB with educational attainment strongly suggested a bidirectional association between earlier reproductive behaviour and shorter years in education. This may help to answer a persistent question in the demographic literature about the causality between reproductive behaviour and educational attainment.[102,103]

The associations with diseases in later life confirmed that there was a strong association of years in education with onset of disease later in life (Figure S16, Table 13B). However, in both cases the association with education was substantially attenuated by the inclusion of the betas for the SNP effects on AFB. These results hold despite the relatively weaker estimates that use the SNPs discovered specifically for AFB (potentially due to comparatively small numbers of SNPs in the analysis). The attenuation was also seen in models adjusting for BMI, suggesting that there is a specific effect of reproductive timing that is a risk factor for type 2 diabetes in women, but not in men.  This finding holds considerable importance since the majority of research has assumed that it is years of education (or similar socioeconomic proxy measures) that are the causal factors driving many diseases in later life. Our analyses show that once we control for both BMI and reproductive timing, the effect of education is considerably weaker, at least on type 2 diabetes.

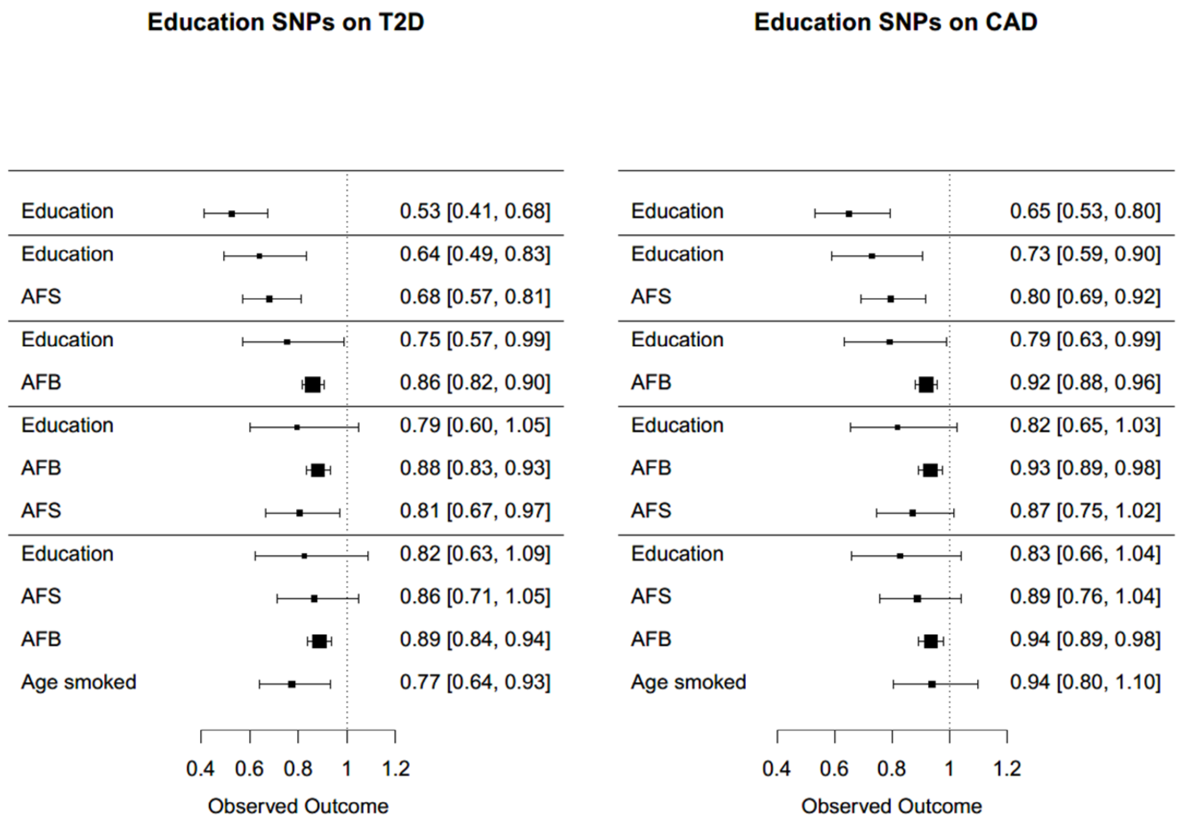

Figure S16. Coefficients (and Cis) of bi-directional MR of human reproductive behaviour (AFB, AFS), age initiated smoking and educational attainment on Type 2 diabetes and Coronary Artery Disease later in life. Finally, since we were also interested in infertility-related phenotypes, bidirectional MR was performed with AFS and AFB with polycystic ovarian syndrome (PCOS) and given the nature of the disease, on women only.[104] Our findings (Supp Note Tab13C) suggest that PCOS leads to later AFB. We find no effect of PCOS on AFS or of either AFS or AFB to PCOS, suggesting that the causal link is infertility-related with PCOS contributing to later AFB.

### 10. Later age at first birth linked to parental longevity

#### 10.1 Background and innovation

Since AFB appears to be predictive of later life onset of disease, we extended the analysis to discover whether reproductive timing was related to longevity. Here we specifically test trade-offs between reproductive behaviour and senescence, which has been argued in the pace of aging literature.[105] The disposable soma theory of the evolution hypothesizes that longevity demands investments in somatic maintenance that in turn reduce the resources that are available for reproduction.[106] Using historical data from the British aristocracy, previous research has shown that AFB was the highest for women who died at the oldest ages and the lowest in women who died early.[107] Using a genealogical database from Utah (1860-1899), researchers applied Cox proportional hazard models to demonstrate that women who had children later and had fewer children enjoyed longer lives.[108] Using contemporary data, another study in the U.S. likewise showed that the odds of longevity and survival to 90 years was significantly higher in women who had a later age at first childbirth.[109]

Using the PGS from our 2016 study of AFS, Mostafavi et al. also previously examined this question.[110] We improve that previous analyses in several distinct ways. First, we use the entire UK Biobank sample, which is considerably larger. Second, the PGSs are calculated using a k-folds cross-validation procedure. Third, we embrace the nature of the data to consider right censoring (i.e., those alive at the time of observation) and estimate survival models.[34] Fourth, we control for other related PGSs such as educational attainment and risky behaviour. Fifth, we stratify by the local authority district at birth, which has been shown to be important for life expectancy in the UK. Finally, we also control for parental fertility by adding the controls for the number of siblings.

#### 10.2 Data and measurement

In the UK Biobank, each individual was asked to provide the age of their mother and father at each visit and also the reported age at death of each parent if applicable. To conduct the survival analysis, we used the most recent assessment visit or average ages reported at recruitment and any repeated assessment visits. For parents who were still alive at the date of the last assessment (i.e., right censored),[34] we included that date as the last observation. We also removed adopted individuals, respondents with non-European Ancestry or those who self-identify as Non-White and those who had missing values for some of the covariates (N=48,980), resulting in 474,946 European ancestry individuals with age at death information for their mother (446,419) and their father (438,125).

#### 10.3 Methods of analysis

We calculated PGSs for AFB, Educational attainment (EA)[48] and risky behaviour[44] from UK Biobank adopting the following procedure. We first split the sample in 10 random groups. We then iteratively estimated genome-wide association results for 9/10$^{th}$ of the sample and used these association results as weights for the calculation of polygenic scores in the remaining 1/10$^{th}$ of the sample. Polygenic scores are calculated using PRSice on a set of independent genotyped SNPs. We then estimated three sets of Cox Proportional hazard models to estimate the effect of the PGS of AFB on maternal and paternal age at death. All models controls for the first 10 Genetic Principal Components, sex and year of birth of the respondents and are stratified by Local Authority District at birth calculated using the geo-coordinated provided in the UK Biobank. This is due to the fact that there is considerable geographical variation in life expectancy at birth in England and Wales, which is largely attributed to differences in material deprivation.[111] Model (1) and (4) in Table S14 are the baseline models. Models (2) and (5) include as covariates the polygenic scores of Educational Attainment and Risky behaviour (high value of the PGS correspond to higher risk adversity). Model (3) and (6) include number of siblings (as proxy for parental fertility) as covariates. These models restrict the analysis to mortality after age 60 to limit the possibility that early mortality affects parental fertility (collider bias).[112]

#### 10.4 Results: Later reproductive timing predicts parental longevity

Results indicate that 1 standard deviation in the PGS for AFB is associated to a reduction in mortality between 2-4% at any age, which holds across all model specifications. As respondents' PGSs are only a proxy of parental genetic predisposition, these estimates are likely affected random measurement errors, leading to attenuation bias. Results are consistent with the work of Mostafavi et al.,[110] who used a PGS derived from our previous 2016 study, but with limitations which our study improves upon, discussed previously. Overall, this analysis shows a common genetic basis of late fertility and longer lifespan and confirms that human reproductive timing and life histories involve a trade-off between parental longevity and reproduction.

Table S14. Polygenic score (PGS) prediction of age at first birth (AFB), educational attainment (EA) and risk on parental longevity

| | **Maternal Age at Death** | | | **Paternal Age at Death** | | |
|---|---|---|---|---|---|---|
| | Model (1) | Model (2) | Model (3) | Model (4) | Model (5) | Model (6) |
| PGS AFB | 0.976*** | 0.966*** | 0.961*** | 0.979*** | 0.971*** | 0.969*** |
| | (0.00203) | (0.00230) | (0.00409) | (0.00182) | (0.00207) | (0.00380) |
| PGS EA | | 0.969*** | 0.968*** | | 0.973*** | 0.974*** |
| | | (0.00209) | (0.00371) | | (0.00188) | (0.00347) |
| PGS Risk | | 0.995** | 0.985*** | | 0.997 | 0.992** |
| | | (0.00230) | (0.00406) | | (0.00206) | (0.00378) |
| Number of Siblings | | | 1.053*** | | | 1.051*** |
| | | | (0.00188) | | | (0.00178) |
| Observations | 398,448 | 398,448 | 132,553 | 391,108 | 391,108 | 120,725 |
| Survival over age 60 | NO | NO | YES | NO | NO | YES |

Note: PGS = Polygenic Score; AFB = age at first birth; EA=Educational attainment PGS.[48] Relative Risk Ratios, exponentiated SEs in parentheses. All models control for first 10 Principal Components, Respondents' Year of Birth, and Sex. All models are stratified by Local Authority District at Birth. *** p<0.01, ** p<0.05, * p<0.1

### 11. Gene prioritization

#### 11.1 Methods

We used multiple approaches to prioritize the most likely causal gene(s) at loci identified as being associated with AFS and/or AFB with CELLECT. First, DEPICT was used to perform pathway analyses, identify enrichment for cell types and tissues, and prioritize candidate genes.[113] DEPICT is agnostic to the outcomes analysed in the GWAS and employs predicted gene functions. For both AFS and AFB, all SNPs with $P<1x10^{-5}$ in the pooled analysis were used as input. For both outcomes, DEPICT's tissue enrichment analysis showed significant enrichment for tissues in the nervous system (Supp Tables S15A (AFS) S15B (AFB)). DEPICT's integrated gene prioritization approach yielded 94 genes for AFS and 14 genes for AFB at FDR<0.05 (Supp Tables S15C (AFS), S15D (AFB)). Based on the results of the tissue enrichment analysis, we next used DEPICT to identify nervous system cell types that are enriched for expression of genes in loci reaching $P<1x10^{-5}$ in the GWAS, using RNAseq data from mouse brain.[114] This yielded 41 enriched cell types for AFS, and 14 for AFB (Supp Tables S15E (AFS), S15F (AFB)). A similar approach using tabula muris RNAseq data[115] helped prioritize another nine central nervous system and pancreatic cell types for AFS (Supp Table S15F). For enriched cell types from mouse brain and tabula muris, the top-10 contributing genes were selected as candidate genes. This resulted in the prioritization of 296 genes for AFS and 95 for AFB based on mouse brain; and 97 genes for AFS based on tabula muris data.

Secondly, we used Phenolyzer (v1.1), to prioritize candidate genes by integrating prior knowledge and phenotype information.[116] Here we used the regions defined by DEPICT v1.1 (see above), reflecting loci reaching $P<1x10^{-5}$ in first instance. Phenolyzer takes free text input and interprets these as disease names by using a word cloud to identify synonyms. It then queries precompiled databases for the disease names to find and score relevant seed genes. The seed genes are subsequently expanded to include related (predicted) genes based on several types of relationships, e.g., protein-protein interactions, transcriptional regulation and biological pathways. Phenolyzer uses machine learning techniques on seed genes and predicted gene rankings to produce an integrated score for each gene. We used search terms capturing three broad areas, i.e. (in)fertility, congenital neurological disorders and psychological traits, based on results from pathway, tissue and cell type enrichment analyses (Supp tables S16A-B). Phenolyzer identified 107 and 47 candidate genes with a score >0.3 for AFS and AFB, respectively. Results for the top candidate genes identified by Phenolyzer can be found in Box 1 below.

**Box 1 – Literature/text mining using Phenolyzer v1.1**

We used Phenolyzer v1.1[116] to identify genes that may be involved in age at first sex and age at first birth using search terms related to psychological traits, infertility and neurological disorders (Supp Table S16A). A total of 107 and 47 candidate genes were identified for AFS and AFB, respectively. Reassuringly some well-known genes related to fertility were prioritized.

The top six genes prioritized for AFS were: *FGFR1*: *ESR1*; *GATA4*; *LEPR*; *CYP17A1* and *CGA*.

***FGFR1*** (Fibroblast growth factor receptor 1) and *FGFR1* with depression. Mutations in *FGFR1* have been associated with Kallmann syndrome, a heterogeneous genetic disorder that associates variable gonadotropin-releasing hormone (GnRH) deficiency with anosmia and, sometimes, other nonreproductive clinical features[117] and that is characterised by decreased testosterone, azoospermia and infertility (ORPHANET:478). The gene has also been associated with lower fertility (HP:0000144); non-obstructive azoospermia (HP:0011961); and lower serum testosterone (HP:0040171).

***ESR1*** (Estrogen receptor 1) is related to multiple fertility traits, including endometriosis,[118] age at menarche[119], male infertility[120] and our previous study on age at first birth.[64] *ESR1* has been associated with alcoholism,[121] psychosis neuroticism[122] and substance-induced schizophrenia (umls:C0033941).

***GATA4*** (GATA Binding Protein 4) was identified through human phenotype ontology (HPO) term and gene associations. The gene has been associated with testicular abnormalities (OMIM:615542). Through HPO to gene associations it has been associated with abnormal spermatogenesis, decreased serum testosterone and abnormal circulating follicle-stimulating hormone (FSH) levels.

***LEPR*** (Leptin receptor) has been associated with infertility, delayed puberty, decreased serum testosterone levels, abnormal serum estradiol through HPO phenotype to genotype associations and DisGeNET(disgenet.org). The gene has also been associated with impairment in personality functioning (HPO phenotype gene association).

***CYP17A1*** (Cytochrome P450 Family 17 Subfamily A Member 1) is a key enzyme in the steroidogenic pathway that produces progestins, mineralocorticoids, glucocorticoids, androgens and estrogens.[123,124] Mutations in this gene have been associated with endometriosis,[125] recurrent pregnancy loss,[126] age at menarche,[127] and serum estrogen and progesterone levels.[124]

***CGA*** (Glycoprotein Hormones, Alpha Polypeptide) has been associated with ectopic pregnancy and is known to interact with *FSHB* (Follicle Stimulating Hormone Subunit Beta),[128] *LHB* (Luteinizing Hormone Beta Polypeptide),[129] and *LHCGR* (Luteinizing Hormone/Choriogonadotropin Receptor).[130]

The top 6 genes prioritized for AFB were: *FSHB*; *ESR1*; *GNAI2*; *RHOA*; *HDAC3*; and *CDC42*.

***FSHB*** (Follicle Stimulating Hormone Subunit Beta) has been associated with FSH deficiency (OMIM:229070), polycystic ovarian syndrome[131] and through HPO phenotype to gene and DisGeNET with female and male infertility and oligospermia.

***GNAI2*** (Guanine Nucleotide Binding Protein (G Protein), Alpha Inhibiting Activity Polypeptide 2) was prioritized for psychological disorders and interacts with other genes associated with cannabis dependence (Cannabinoid Receptor 1),[132] but also with genes related to fertility, like *LHB*.[133] The gene has also been associated with schizophrenia through DisGeNET.

The ***RHOA*** (Ras Homolog Family Member A) locus has been associated with general cognitive ability.[134]

***HDAC3*** (Histone Deacetylase 3) has been shown to interact with genes related to psychological disorders like JUN,[135,136] and genes related to male fertility (*ARID4A*).[137,138]

Thirdly, we used *in silico* sequencing to identify non-synonymous variants with an $R^2$ for LD>0.7 with the lead SNPs in AFS and AFB-associated loci.[139] This yielded 24 genes for AFS and 16 for AFB that may drive the GWAS associations through direct effects on protein function (Supp Table S17A-B).

Fourthly, we used Summary data-based Mendelian Randomization (SMR) and heterogeneity in dependent instruments (HEIDI)[140] using eQTL data from brain[141] and whole blood.[142] This approach provided 39 and 73 genes that showed evidence ($P_{SMR}$<5x10$^{-6}$) of mediating the association between AFS and GWAS identified loci based on results from brain and blood, respectively, compared with 15 and 29 genes for AFB (Supp Table S18A (AFS), S18B (AFB)).

Finally, we integrated findings across all approaches and retained genes in loci that reached genome-wide significance, and that were located within 1M bp of a GWAS lead SNP. This resulted in the prioritization of 314 genes in 153 loci for AFS, and 106 genes in 37 loci for AFB (Supp Tables S19A (AFS), S19B (AFB)), or 386 unique genes across the two traits (Supp Table S19C). We next used data from the Human Protein Atlas[143] to identify genes amongst these 386 genes that are expressed at a low, medium or high protein level in brain, glands, and/or reproductive organs at a 'supported' or 'enhanced' degree of reliability. For the 99 genes that fulfilled these criteria, we mapped the brain, glandular and reproductive cell types in which they are highly expressed at the protein level;[144] used a text-mining approach to extract functions from entries in Entrez, GeneCards and Uniprot; and identified phenotypes in mutant mice from the Mouse Genome Informatics (MGI) database[145] (Figure 4, Main Text).

#### 11.2 Results

Gene prioritization using multiple approaches (see ***Methods***) highlighted genes acting in three broad tissue types: brain, glands and reproductive organs. Some of these have known effects on traits related to cognitive ability, addiction, psychiatric traits and fertility (see below). These results partly mirror and compliment the rigorous post-GWAS *in silico* association analyses we performed for loci identified for age at first sex and age at first birth.

##### 11.2.1 Candidate genes in brain

Twenty-four of the 99 prioritized genes are both highly expressed in central nervous system cell types at the protein level,[144] and yield a nervous system or neurological phenotype in mutant mice.[146] Within these 24 genes, STRING databases[147] highlighted experimentally determined protein-protein interactions of *HDAC3* with *GTF2I*, *TOP2B*, *E2F1* and *MEF2C* (Supp Fig S16). All five genes are highly expressed at the protein level in neuronal cells of the cerebral cortex, and all but *MEF2C* are highly expressed in Purkinje cells and the molecular layer of the cerebellum. Histone deacetylase 3 (*HDAC3*) is essential for Purkinje cell function,[148] embryonic brain development and neuro-differentiation,[149] long-term memory formation,[150] and blood brain barrier permeability;[151] loss of general transcription factor IIi (*GTF2I*) induces increased sociability and anxiety in mice;[152,153] DNA topoisomerase II beta (*TOP2B*) is required for transcription of a set of long neuronal genes in cerebellar granule neurons,[154] as well as for development and survival of post-mitotic neurons in the retina;[155] E2F transcription factor 1 (*E2F1*) contributes to Purkinje cell degeneration,[156] modulates neuronal apoptosis,[157,158] and functions as a cell cycle suppressor in mature neurons;[159] and myocyte enhancer factor 2C (*MEF2C*) plays a key role in cortical network activity by regulating inhibitory vs. excitatory synaptic transmission.[160]

*NCAM1* and *NFASC* also interact at the protein level (Supp Fig S16). Both are highly expressed at the protein level in neuropils of the cerebral cortex, and both play a role in development of the nervous system.[161] Neural cell adhesion molecule 1 (*Ncam1*) null mice are prone to risk seeking behavior,[162] and plasma NCAM1 levels were negatively associated with social motivation, communication, and responsiveness in children with autism spectrum disorders.[163] Neurofascin (*NFASC*) plays a key role in formation of the nodes of Ranvier and function of myelinated axons;[164] mutations in this gene cause severe neurodevelopmental disorders.[165]

##### 11.2.2 Candidate genes in glands

Twelve of the 99 prioritized genes are highly expressed at the protein level in glands,[144] and additionally yield an endocrine or exocrine phenotype in mutant mice.[146] Of these, experimentally determined protein-protein interactions were observed for *SUMO1* with *RECQL4* and *PML* (Supp Fig S16), which are highly expressed at the protein level in glandular cells of the adrenal (*SUMO1* and *RECQL4*), parathyroid (*SUMO1*) and/or thyroid glands (*RECQL4* and *PML*). Small ubiquitin-related modifier 1 (*SUMO1*) binds target proteins as part of a post-translational modification system - i.e. sumoylation - and plays a role in nuclear transport, transcriptional regulation, apoptosis and protein stability. RecQ like helicase 4 (*RECQL4*) plays an essential role in DNA replication during development and is required for viability and fertility,[166] while promyelocytic leukemia (*PML*) is recruited to phosphorylated testis receptor 2 and sumoylated, which in turn suppresses cell proliferation.[167]

A protein-protein interaction was also observed between *CGA* and *FSHB* (Supp Fig S16), which are both highly expressed in anterior pituitary gland cells. *CGA* encodes the alpha subunit of follicle stimulating hormone (FSH) – as well as three other glycoprotein hormones – and plays a role in ectopic pregnancies through interaction with Forkhead Box L2 (*FOXL2*).[168] *FSHB* encodes the beta subunit of FSH, which stimulates the growth of ovarian follicles in women, and acts on the Sertoli cells of the testis to stimulate sperm production in men. Mutations in *FSHB* have been reported in infertile men,[169,170] and in women with isolated FSH deficiency and hypogonadism.[171]

##### 11.2.3 Candidate genes in female reproductive organs

Nine of the 99 prioritized genes are highly expressed at the protein level in female reproductive organs[144] and additionally show a reproductive phenotype in mutant mice.[146] Of these nine genes, *ESR1* interacts at the protein level with *SUMO1*, *ARNT*, *CAV1* and *E2F1* (Supp Fig S16). Small ubiquitin-like modifier 1 (*SUMO1*) sumoylates estrogen receptor (ER)alpha (*ESR1*) in the presence of estrogen, which is a requirement for normal ERalpha-induced transcription.[172] SUMO-1 and ERalpha are both highly expressed at the protein level in glandular cells of the fallopian tube and endometrium. SUMO-1 also co-localizes with Forkhead Box L2 (*FOXL2*) in stromal and glandular cells of the endometrium. Forkhead Box L2 has been implicated in the pathogenesis of endometriosis[173] and ectopic pregnancies.[174]

Knockdown of Aryl hydrocarbon receptor nuclear translocator (*ARNT*) has been shown to suppress key angiogenic genes - including *VEGFA*, possibly through FSH[175] - leading to deficient angiogenesis in placental vasculature; malformed thin villous vessels; elevated fetoplacental vascular resistance; and high morbidity and mortality in fetal growth restriction.[176]

Caveolin 1 (*CAV1*) promotes human trophoblast cell proliferation, migration and invasion by activating the focal adhesion kinase signaling pathway. Appropriate differentiation and invasion of trophoblast cells is required for normal implantation and placental development, and caveolin 1 gene expression was lower in placenta of unexplained spontaneous abortions than in placenta from induced abortions.[177]

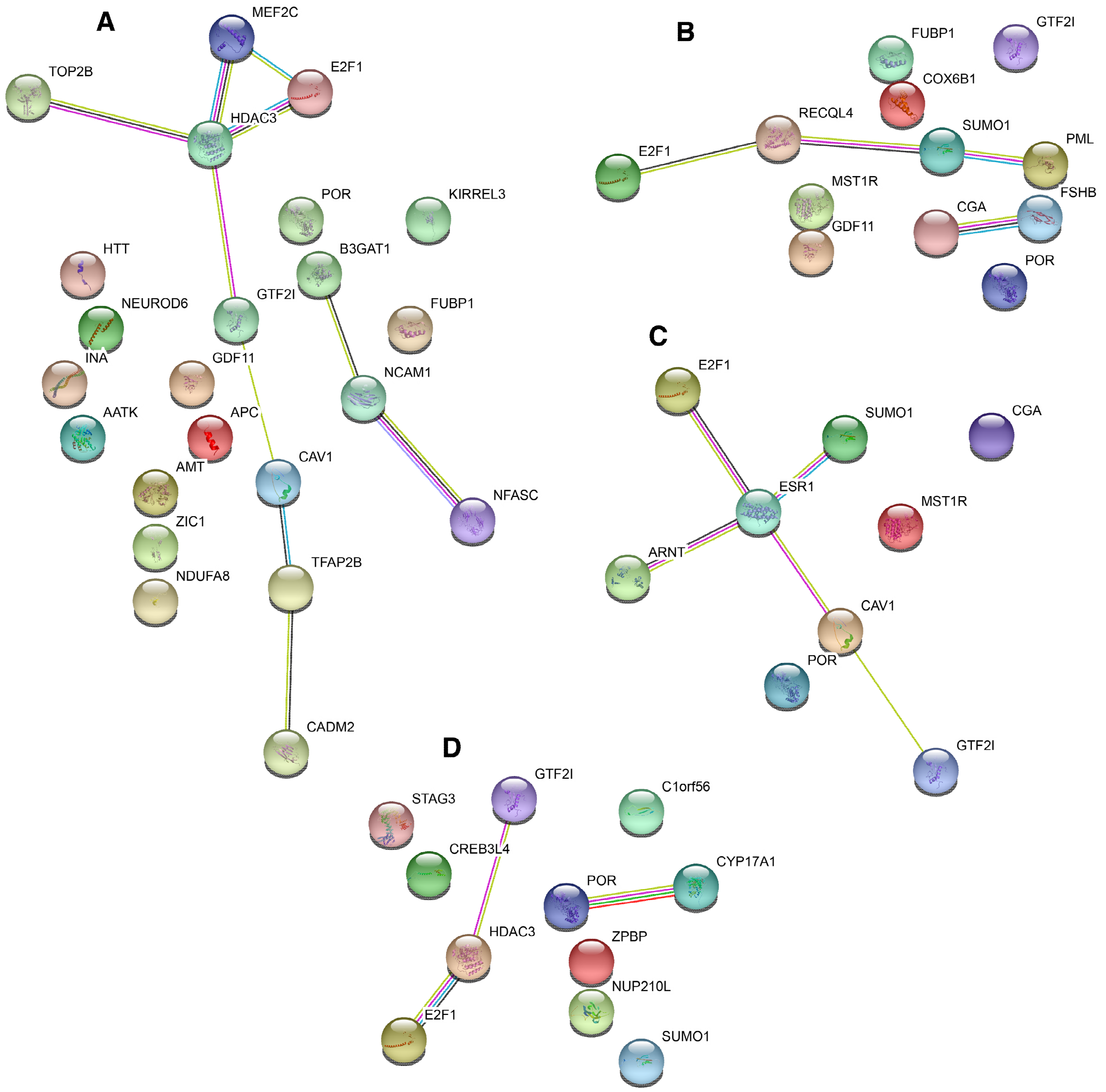

Figure S17. Protein-protein interactions identified using STRING for genes that are highly expressed at the protein level in: A) brain and result in a nervous system or neurological behavior phenotype in mutant mice; B) glands and result in an endocrine/exocrine phenotype in mutant mice; C-D) female (C) or male (D) reproductive organs and result in a reproductive phenotype in mutant mice. Pink lines highlight experimentally determined interactions.

In A, HDAC3, TOP2B, GTF2I and E2F1 are all highly expressed at the protein level in Purkinje and molecular layer cells in the cerebellum and neuronal cells in the cerebral cortex; HDAC3 and MEF2C are both highly expressed in neuronal cells of the cerebral cortex; and NCAM1 and NFASC are both highly expressed in neuropils of the cerebral cortex. In B, SUMO1 and RECQL4 are both highly expressed in glandular cells of the adrenal gland, SUMO1 is highly expressed in glandular cells of the parathyroid gland; and RECQL4 and PML are both highly expressed in glandular cells of the thyroid gland. CGA and FSHB are both highly expressed in cells in the anterior pituitary gland. In C, SUMO1 and ESR1 are both highly expressed in glandular cells of the endometrium and fallopian tube; ARNT and ESR1 both are highly expressed in endometrial stroma cells and glandular cells of the fallopian tube; E2F1 and ESR1 are both highly expressed in squamous epithelial cells of the vagina; and CAV1 and ESR1 are both highly expressed in endometrial stroma cells. In D, GTF2I and HDAC3 are both highly expressed in epididymis glandular cells, and POR and CYP17A1 are both highly expressed in Leydig cells of the testis.

E2F transcription factor 1 (*E2F1*) is one of 11 genes in the PI3K/AKT pathway with a lower expression in cumulus cells from oocytes that went on to produce a pregnancy vs. those that did not. This differential expression was concluded to likely be driven by a downregulation of ERalpha.[178] In separate efforts, E2F1 was identified as one of six transcription factors that likely induce pregnancy-induced pancreatic islet expansion;[179] and was shown to activate ribonucleotide reductase 2 - an important effector of progesterone signalling - to induce cell proliferation and decidualization in mouse uterus.[180]

##### 11.2.4 Candidate genes in male reproductive organs

Of the 11 genes that are highly expressed at the protein level in male reproductive tissues[144] and additionally show a reproductive phenotype in mutant mice,[146] a protein-protein interaction was observed of *HDAC3* with *GTF2I* and *E2F1* (Supp Fig S16). While the mechanisms by which *HDAC3* with *GTF2I* influence reproductive behavior in male reproductive tissues remain to be established, loss of E2F transcription factor 1 (*E2F1*) has been shown to induce severe and progressive testicular atrophy and less spermatogonia apoptosis during the first wave of spermatogenesis in young mice, and resulted in further exacerbation of testicular atrophy due to loss of spermatocytes in adult mice by loss of spermatogonia stem cells.[181]

A second protein-protein interaction was observed for *CYP17A1* and *POR* (Supp Fig S16), which are both highly expressed at the protein level in the testis' Leydig cells. Cytochrome P450 family 17 subfamily A member 1 (*CYP17A1*) is a key enzyme in the steroidogenic pathway that produces progestins, mineralcorticoids, glucocorticoids, androgens and estrogens,[182] while cytochrome p450 oxidoreductase (*POR*) is required for normal steroidogenesis.[183]

Seven of the 97 prioritized genes were highly expressed at the protein level in spermatogonia, preleptotene spermatocytes, pachytene spermatocytes, round or early spermatids, and/or elongated or late spermatids (Figure 5, Main Text).[144] Of these, Krueppel-like factor 17 (*KLF17*) encodes a germ cell-specific transcription factor that in mice plays important roles in spermatid differentiation and oocyte development;[184,185] and zona pellucida binding protein (*ZPBP*) participates in sperm morphogenesis and binding between acrosome-reacted sperm and the egg-specific extracellular matrix (the zona pellucida.[186,187] Furthermore, epigenetic, allele-specific activation of the testis-specific gene nucleoporin 210 like (*NUP210L*) – possibly by changing binding affinity for testis receptor 2 - was recently observed in prefrontal cortex neurons of G allele carriers (but not CC carriers) in rs114697636 (MAF 3%).[188] Rs114697636 is in linkage disequilibrium with the locus' lead SNP for AFS (rs113142203, D' 0.90). Furthermore, the DNA methylation state of *NUP210L* has recently been linked with psychologic development disorders,[189] and common variants near *NUP210L* have been identified in GWAS for intelligence and mathematical ability,[48,190] providing an elegant example of how a testis-specific gene that is highly expressed at the protein level in developing and mature sperm can influence the brain in some individuals. The roles of *ELAVL2*, *LRRC37A2*, *C1orf56* and *C20orf144* in reproductive behavior through male reproductive organs remain to be established.

#### 12. Sex-specific genetic effects

Sex-specific genetic effects have been proposed and found previously for reproductive behaviour.[2,191,192] Sex-specific effects in these behavioural phenotypes are likely driven by biological, behavioural and social normative and cultural factors. First, there are sex differences in the biological makeup, processes and diseases that are related to sexual and fertility behaviour and related diseases.[193,194] A later age at first birth has been related to infertility and reproduction related traits in women, such as ovulatory problems, tubal damage, endometriosis, cervical cancer and polycystic ovarian syndrome.[192] Fecundability is also influenced by sex-specific hormonal processes,[195] as well as by a behavioural component, e.g. through educational attainment, personality, risk, or impulsivity.[196] In turn, these traits have differential effects on male and female reproduction.[197,198] In addition to the pooled GWAS, we also ran sex-specific GWAS meta-analyses for both phenotypes. In doing so, we detected two genome-wide significant ($p$-value<$5x10^{-8}$) loci for AFS in women, eight for AFS in men and one for AFB in women. Gene prioritization in sex-specific loci resulted in the prioritization of 11 genes for AFB in women, one gene for AFS in women and 23 genes for AFS in men. Of these, 12 genes at three loci were expressed at the protein level in relevant tissues (Figure S17).

##### 12.1 Genetic overlap among sexes: LD score bivariate regression

We used LD score bivariate regression[199] to estimate the genetic correlation between men and women based on the sex-specific summary statistics from the meta-analysis results. Figure S18 shows the genetic correlations across the traits by sex and Figure S19, gene prioritization by sex. Considering the high correlations, these results indicate a large genetic overlap among the sexes, particularly for AFB (0.95). We see, however, that the genetic overlap between men and women is still high, but lower for AFS at rg=0.79.

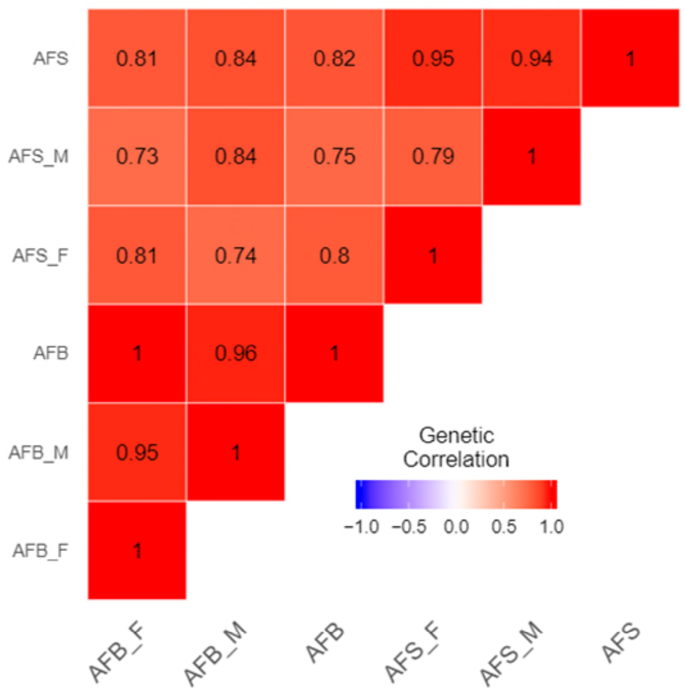

Figure S18. Genetic overlap amongst the sexes for AFS and AFB, LD score bivariate regression

Figure S19. Gene prioritization of AFS and AFB by sex

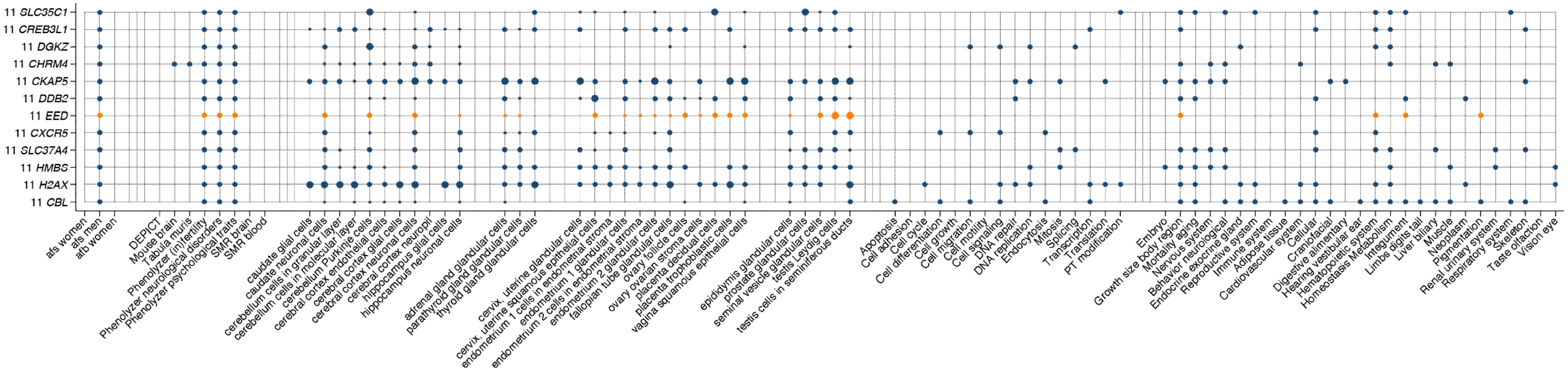

#### 12.2 Sex specific loci

##### 12.2.1 Methods and identification of 10 additional associations for AFS and 1 for AFB

Considering the lower sex-specific correlation particularly with AFS, we opted to examine sex-specific loci in more detail. A total of 242 unique variants were associated with either AFS or AFB in the sex stratified analyses. In order to determine if there was evidence for sex-specific effects, we compared the allelic effects for these SNPs between men and women and derived a *p*-value for heterogeneity.[200] Based on a multiple testing correction for the number of variants ($0.05/242=2x10^{-4}$), we identified for: (1) AFS, 2 sex specific associations in women and 8 in men; and, (2) AFB, 1 sex specific association in women only. We selected a region of 2Mb around these lead SNPs to identify the genes that may be represented by these lead SNPs. We then conducted gene prioritization as we did for the main AFB and AFS analyses.

##### 12.2.2 Gene Prioritization Results AFS

***AFS: Women.*** There were two genes prioritized for AFS in women: *FANCL* and *ERBB4*. *FANCL* is a member of the Fanconi anemia complementation group. Variants in this locus have also been associated with smoking, autism and schizophrenia.[44,201] Orphanet (84) and DISGENET (umls:C0015625) links the gene to fanconi anemia and delayed puberty. ERBB4 has been linked to abnormal fear/anxiety-related behavior (HP:0100852 (HPO_PHENOTYPE_GENE)), and schizophrenia[202] as well as PCOS.[203]

***AFS: Men***. We prioritised genes related to mental development and psychological disorders. NRXN1 deletions in this gene have been associated with schizophrenia and autism;[204] SLC44A1 is a choline transporter. Choline is key in cerebral inhibition. Abnormal inhibition has been implicated in a number of neuropsychiatric illnesses such as substance abuse and depressive disorders;[205] NR1H3 (LXRalpha) may have links to major depressive disorder;[206] and H2FAX has been linked to Ataxia-telangiectasia neurogenerative disorder and Nijmegen Breakage Syndrome (NBS) microcephaly and short stature[207] and higher chance of cancer.

There were other genes related to haematological factors that were prioritised for AFS in men: *F2* and *HMBS*. F2 has been related to miscarriage and heavy menstrual bleeding (OMIM:614390).[208] Neither of which can be related to men. *SLC24A5* is another gene that was prioritised and is a major locus for skin pigmentation and is known to be under selection.[209]

##### 12.2.3 Gene Prioritization Results AFB

***AFB: Women***. Recall that one sex-specific loci was found for women. In the region of the lead variant there are many genes prioritised related to immunity and specifically, cheomokine receptors. CCR1 protein and expression is increased in women with endometriosis.[210] CCR5 is decreased in swim-up sperm in infertile men and that may be associated with male infertility.[211] CXCR6 is the receptor for CXCL16. Disruption in this pathway in the endometrium has been observed in women who experience spontaneous abortion.[212]

### 13. Authorship

Melinda C. Mills[1,2,†,*], Felix C. Tropf[1,2,3,4,†], David M. Brazel[1,2,†], Natalie van Zuydam[5], Ahmad Vaez[6,7], *eQTLGen Consortium*, *BIOS Consortium*, Tune H. Pers[8,9], Harold Snieder[6], John R.B. Perry[10], Ken K. Ong[10], Marcel den Hoed[5,†], Nicola Barban[11,†], and Felix R. Day[10,†,*] on behalf of the *Human Reproductive Behaviour Consortium*

#### Corresponding authors

#### Author contributions

MCM and FRD designed and led the study. MCM wrote the paper and supplementary note with contributions by authors for respective analyses and comments by all authors. DMB conducted phenotypic changes, phenotype preparation, LD Score and genetic correlations, Genomic SEM and exploratory factor analysis and sex-specific effects. NB conducted GWAS meta-analysis, MTAG, PGS prediction, survival models, and Cox models of longevity. FCT and FRD conducted the cohort QC. FCT conducted GREML cohort heritability analysis and phenotype preparation in UKBB. FRD ran Mendelian Randomization, conducted GWAS analyses and JRBP conducted COJO and X-Chromosome analysis. NvZ conducted DEPICT and Phenolyzer analyses. AV and HS conducted in silico sequencing and SMR analyses. TP conducted cell type enrichment analyses. MdH integrated gene prioritization results and performed downstream analyses, e.g. Human Protein Atlas; Entrez, GeneCards and Uniprot mining; and STRING Protein-Protein interaction analyses. Authors in the Human Reproductive Behaviour Consortium conducted cohort specific GWAS and other analyses, and contributed through the administration, management, and data collection for the participating cohorts. The eQTLGen and BIOS Consortiums provided data for additional analyses. All authors reviewed and approved the final version of the paper.

#### Author information

These authors contributed equally: Melinda C. Mills, Felix C. Tropf, David M. Brazel, Felix R. Day, Nicola Barban, Marcel den Hoed

These authors designed and led the study: Melinda C. Mills, Felix R. Day

#### Affiliations

**Leverhulme Centre for Demographic Science, University of Oxford & Nuffield College, UK**

Melinda C. Mills, Felix C. Tropf, David M. Brazel

**MRC Epidemiology Unit, Institute of Metabolic Science, University of Cambridge, Cambridge, United Kingdom**

Felix R. Day, John R.B. Perry, Ken K. Ong

**Department of Statistics, University of Bologna, Bologna, Italy**

Nicola Barban

**École Nationale de la Statistique et de L'administration Économique (ENSAE) & Center for Research in Economics and Statistics (CREST), Paris, France**

Felix C. Tropf

**Department of Epidemiology, University of Groningen, University Medical Center Groningen, Groningen, The Netherlands**

Harold Snieder, Ahmad Vaez

**The Beijer Laboratory and Department of Immunology, Genetics and Pathology, Uppsala University and SciLifeLab, Uppsala, Sweden**

Marcel den Hoed, Natalie van Zuydam

**The Novo Nordisk Foundation Center for Basic Metabolic Research, Faculty of Health and Medical Sciences, University of Copenhagen, Copenhagen, Denmark**

Tũne H. Pers

**Department of Bioinformatics, Isfahan University of Medical Sciences, Isfahan, Iran**

Ahmad Vaez

### Consortia

#### Human Reproductive Behaviour Consortium

Author list ordered alphabetically

Evelina T. Akimova, Sven Bergmann, Jason D. Boardman, Dorret I. Boomsma, Marco Brumat, Julie E. Buring, David Cesarini, Daniel I. Chasman, Jorge E. Chavarro, Massimiliano Cocca, Maria Pina Concas, George Davey-Smith, Gail Davies, Ian J. Deary, Tõnu Esko, Oscar Franco, Audrey J. Gaskins, Eco J.C. de Geus, Christian Gieger, Giorgia Girotto, Hans Jörgen Grabe, Erica P. Gunderson, Kathleen Mullan Harris, Fernando P. Hartwig, Chunyan He, Diana van Heemst, W. David Hill, Georg Homuth, Bernando Lessa Horta, Jouke Jan Hottenga, Hongyang Huang, Elina Hyppönen, M. Arfan Ikram, Rick Jansen, Magnus Johannesson, Zoha Kamali, Maryam Kavousi, Peter Kraft, Brigitte Kühnel, Claudia Langenberg, Lifelines Cohort Study, Penelope A. Lind, Jian'an Luan, Reedik Mägi, Patrik K.E. Magnusson, Anubha Mahajan, Nicholas G. Martin, Hamdi Mbarek, Mark I. McCarthy, George McMahon, Matthew B. McQueen, Sarah E. Medland, Thomas Meitinger, Andres Metspalu, Evelin Mihailov, Lili Milani, Stacey A. Missmer, Stine Møllegaard, Dennis O. Mook-Kanamori, Anna Morgan, Peter J. van der Most, Renée de Mutsert, Matthias Nauck, Ilja M. Nolte, Raymond Noordam, Brenda W.J.H. Penninx, Annette Peters, Chris Power, Paul Redmond, Janet W. Rich-Edwards, Paul M. Ridker, Cornelius A. Rietveld, Susan M. Ring, Lynda M. Rose, Rico Rueedi, Kári Stefánsson, Doris Stöckl, Konstantin Strauch, Morris A. Swertz, Alexander Teumer, Gudmar Thorleifsson, Unnur Thorsteinsdottir, A. Roy Thurik, Nicholas J. Timpson, Constance Turman, André G. Uitterlinden, Melanie Waldenberger, Nicholas J. Wareham, Gonneke Willemsen, and Jing Hau Zhao

#### Affiliations

Author list ordered alphabetically

**Leverhulme Centre for Demographic Science, University of Oxford, Oxford, United Kingdom**

Evelina T. Akimova

**Department of Computational Biology, University of Lausanne, Lausanne, Switzerland**

Sven Bergmann, Rico Rueedi

**Swiss Institute of Bioinformatics, Lausanne, Switzerland**

Sven Bergmann, Rico Rueedi

**Department of Integrative Biomedical Sciences, University of Cape Town, Cape Town, South Africa**

Sven Bergmann

**Department of Sociology and Institute of Behavioral Science, University of Colorado at Boulder, Boulder, CO, United States of America**

Jason D. Boardman

**Department of Biological Psychology, Amsterdam Public Health Research Institute, Vrije Universiteit Amsterdam, Amsterdam, The Netherlands**

Dorret I. Boomsma, Eco J.C. de Geus, Jouke Jan Hottenga, Hamdi Mbarek, Gonneke Willemsen

**Department of Medical, Surgical and Health Sciences, University of Trieste, Trieste, Italy**

Marco Brumat, Giorgia Girotto

**Brigham and Women's Hospital, Boston, MA, United States of America**

Julie E. Buring, Daniel I. Chasman, Lynda M. Rose, Paul M. Ridker

**Harvard Medical School, Boston, MA, United States of America**

Julie E. Buring, Daniel I. Chasman, Paul M. Ridker

**Department of Economics, New York University, New York, NY, United States of America**

David Cesarini

**Research Institute for Industrial Economics, Stockholm, Sweden**

David Cesarini

**National Bureau of Economic Research, Cambridge, MA, United States of America**

David Cesarini

**Department of Epidemiology, Harvard T.H. Chan School of Public Health, Boston, MA, United States of America**

Jorge E. Chavarro, Hongyang Huang, Peter Kraft, Stacey A. Missmer, Janet W. Rich-Edwards, Constance Turman

**Department of Nutrition, Harvard T.H. Chan School of Public Health, Boston, MA, United States of America**

Jorge E. Chavarro, Audrey J. Gaskins

**Channing Division of Network Medicine, Brigham and Women's Hospital and Harvard Medical School, Boston, MA, United States of America**

Jorge E. Chavarro, Audrey J. Gaskins, Janet W. Rich-Edwards

**Institute for Maternal and Child Health IRCCS "Burlo Garofolo", Trieste, Italy**

Massimiliano Cocca, Maria Pina Concas, Giorgia Girotto, Anna Morgan

**MRC Integrative Epidemiology Unit, University of Bristol, Bristol, United Kingdom**

George Davey-Smith, Fernando P. Hartwig, Susan M. Ring, Nicholas J. Timpson

**Lothian Birth Cohorts, Department of Psychology, University of Edinburgh, Edinburgh, United Kingdom**

Gail Davies, Ian J. Deary, W. David Hill, Paul Redmond

**Estonian Genome Center, University of Tartu, Tartu, Estonia**

Tõnu Esko, Reedik Mägi, Andres Metspalu, Evelin Mihailov, Lili Milani

**Broad Institute of the Massachusetts Institute of Technology and Harvard University, Cambridge, MA, United States of America**

Tõnu Esko

**Department of Epidemiology, Erasmus Medical Center, Rotterdam, The Netherlands**

Oscar Franco, M. Arfan Ikram, Maryam Kavousi

**Department of Epidemiology, Rollins School of Public Health, Emory University, Atlanta, GA, United States of America**

Audrey J. Gaskins

**Research Unit of Molecular Epidemiology, Helmholtz Zentrum München, German Research Center for Environmental Health, Neuherberg, Germany**

Christian Gieger, Brigitte Kühnel, Melanie Waldenberger

**Department of Psychiatry and Psychotherapy, University Medicine Greifswald, Greifswald, Germany**

Hans Jörgen Grabe

**Division of Research, Kaiser Permanente Northern California, Oakland, CA, United States of America**

Erica P. Gunderson

**Department of Sociology, Carolina Population Center, University of North Carolina at Chapel Hill, Chapel Hill, NC, United States of America**

Kathleen Mullan Harris

**Postgraduate Program in Epidemiology, Federal University of Pelotas, Pelotas, Brazil**

Fernando P. Hartwig, Bernando Lessa Horta

**University of Kentucky Markey Cancer Center, Lexington, KY, United States of America**

Chunyan He

**Department of Internal Medicine, Division of Medical Oncology, University of Kentucky College of Medicine, Lexington, KY, United States of America**

Chunyan He

**Department of Internal Medicine, Section of Gerontology and Geriatrics, Leiden University Medical Center, Leiden, The Netherlands**

Diana van Heemst, Raymond Noordam

**Interfaculty Institute for Genetics and Functional Genomics, University of Greifswald, Greifswald, Germany**

Georg Homuth

**Australian Centre for Precision Health, University of South Australia Cancer Research Institute, Adelaide, Australia**

Elina Hyppönen

**South Australian Health and Medical Research Institute, Adelaide, Australia**

Elina Hyppönen

**Department of Psychiatry, Amsterdam Public Health and Amsterdam Neuroscience, Amsterdam UMC, Vrije Universiteit, Amsterdam, The Netherlands**

Rick Jansen

**Department of Economics, Stockholm School of Economics, Stockholm, Sweden**

Magnus Johannesson

**Department of Bioinformatics, Isfahan University of Medical Sciences, Isfahan, Iran**

Zoha Kamali

**Department of Biostatistics, Harvard T.H. Chan School of Public Health, Boston, MA, United States of America**

Peter Kraft

**MRC Epidemiology Unit, Institute of Metabolic Science, Cambridge Biomedical Campus, University of Cambridge School of Clinical Medicine, Cambridge, United Kingdom**

Claudia Langenberg, Jian'an Luan, Nicholas J. Wareham, Jing Hau Zhao

**Department of Epidemiology, University of Groningen, University Medical Center Groningen, Groningen, The Netherlands**

Zoha Kamali, Lifelines Cohort Study, Peter J. van der Most, Ilja M. Nolte

**Department of Genetics, University of Groningen, University Medical Center Groningen, Groningen, The Netherlands**

Lifelines Cohort Study, Morris A. Swertz

**Psychiatric Genetics, QIMR Berghofer Medical Research Institute, Herston Brisbane, Queensland, Australia**

Penelope A. Lind, Sarah E. Medland

**Department of Medical Epidemiology and Biostatistics, Karolinska Institutet, Stockholm, Sweden**

Patrik K.E. Magnusson

**Wellcome Centre for Human Genetics, University of Oxford, Oxford, United Kingdom**

Anubha Mahajan, Mark I. McCarthy

**Oxford Centre for Diabetes, Endocrinology and Metabolism, Radcliffe Department of Medicine, University of Oxford, Oxford, United Kingdom**

Anubha Mahajan, Mark I. McCarthy

**Genetic Epidemiology, QIMR Berghofer Medical Research Institute, Herston Brisbane, Queensland, Australia**

Nicholas G. Martin

**Qatar Genome Programme, Qatar Foundation, Doha, Qatar**

Hamdi Mbarek

**School of Social and Community Medicine University of Bristol, Bristol, United Kingdom**

George McMahon

**Department of Integrative Physiology, University of Colorado at Boulder, Boulder, CO, United States of America**

Matthew B. McQueen

**Institute of Human Genetics, Helmholtz Zentrum München, German Research Center for Environmental Health, Neuherberg, Germany**

Thomas Meitinger

**Institute of Molecular and Cell Biology, University of Tartu, Tartu, Estonia**

Andres Metspalu

**Division of Adolescent and Young Adult Medicine, Department of Medicine, Boston Children's Hospital and Harvard Medical School, Boston, MA, United States of America**

Stacey A. Missmer

**Department of Obstetrics, Gynecology, and Reproductive Biology, College of Human Medicine, Michigan State University, Grand Rapids, MI, United States of America**

Stacey A. Missmer

**Department of Sociology, University of Copenhagen, Copenhagen, Denmark**

Stine Møllegaard

**Department of Clinical Epidemiology, Leiden University Medical Center, Leiden, The Netherlands**

Dennis O. Mook-Kanamori, Renée de Mutsert

**Department of Public Health and Primary Care, Leiden University Medical Center, Leiden, The Netherlands**

Dennis O. Mook-Kanamori

**Institute of Clinical Chemistry and Laboratory Medicine, University Medicine Greifswald, Greifswald, Germany**

Matthias Nauck

**Department of Psychiatry, EMGO Institute for Health and Care Research and Neuroscience Campus Amsterdam, VU University Medical Center/GGZ inGeest, Amsterdam, The Netherlands**

Brenda W.J.H. Penninx

**Institute of Epidemiology, Helmholtz Zentrum München, German Research Center for Environmental Health, Neuherberg, Germany**

Annette Peters, Doris Stöckl, Melanie Waldenberger

**Population, Policy and Practice Research and Teaching Department, UCL Great Ormond Street Institute of Child Health, London, United Kingdom**

Chris Power

**Division of Women's Health, Department of Medicine, Brigham and Women's Hospital and Harvard Medical School, Boston, MA, United States of America**

Janet W. Rich-Edwards

**Erasmus University Rotterdam Institute for Behavior and Biology, Rotterdam, The Netherlands**

Cornelius A. Rietveld, A. Roy Thurik, André G. Uitterlinden

**Department of Applied Economics, Erasmus School of Economics, Rotterdam, The Netherlands**

Cornelius A. Rietveld, A. Roy Thurik

**deCODE Genetics/Amgen Inc., Reykjavik, Iceland**

Kári Stefánsson, Gudmar Thorleifsson, Unnur Thorsteinsdottir

**Institute of Medical Biostatistics, Epidemiology and Informatics (IMBEI), University Medical Center, Johannes Gutenberg University, Mainz, Germany**

Konstantin Strauch

**Institute of Genetic Epidemiology, Helmholtz Zentrum München, German Research Center for Environmental Health, Neuherberg, Germany**

Konstantin Strauch

**Chair of Genetic Epidemiology, IBE, Faculty of Medicine, LMU Munich, Germany**

Konstantin Strauch

**Institute for Community Medicine, University Medicine Greifswald, Greifswald, Germany**

Alexander Teumer

**Montpellier Business School, Montpellier, France**

A. Roy Thurik

**Department of Internal Medicine, Erasmus University Medical Center, Rotterdam, The Netherlands**

André G. Uitterlinden

#### eQTLGen Consortium

Author list ordered alphabetically

Mawussé Agbessi, Habibul Ahsan, Isabel Alves, Anand Kumar Andiappan, Wibowo Arindrarto, Philip Awadalla, Alexis Battle, Frank Beutner, Marc Jan Bonder, Dorret I. Boomsma, Mark W. Christiansen, Annique Claringbould, Patrick Deelen, Tõnu Esko, Marie-Julie Favé, Lude Franke, Timothy Frayling, Sina A. Gharib, Greg Gibson, Bastiaan T. Heijmans, Gibran Hemani, Rick Jansen, Mika Kähönen, Anette Kalnapenkis, Silva Kasela, Johannes Kettunen, Yungil Kim, Holger Kirsten, Peter Kovacs, Knut Krohn, Jaanika Kronberg, Viktorija Kukushkina, Zoltan Kutalik, Bernett Lee, Terho Lehtimäki, Markus Loeffler, Urko M. Marigorta, Hailang Mei, Lili Milani, Grant W. Montgomery, Martina Müller-Nurasyid, Matthias Nauck, Michel G. Nivard, Brenda Penninx, Markus Perola, Natalia Pervjakova, Brandon L. Pierce, Joseph Powell, Holger Prokisch, Bruce M. Psaty, Olli T. Raitakari, Samuli Ripatti, Olaf Rotzschke, Sina Rüeger, Ashis Saha, Markus Scholz, Katharina Schramm, Ilkka Seppälä, Eline P. Slagboom, Coen D.A. Stehouwer, Michael Stumvoll, Patrick Sullivan, Peter A.C. 't Hoen, Alexander Teumer, Joachim Thiery, Lin Tong, Anke Tönjes, Jenny van Dongen, Maarten van Iterson, Joyce van Meurs, Jan H. Veldink, Joost Verlouw, Peter M. Visscher, Uwe Völker, Urmo Võsa, Harm-Jan Westra, Cisca Wijmenga, Hanieh Yaghootkar, Jian Yang, Biao Zeng, Futao Zhang

#### Affiliations

**Computational Biology, Ontario Institute for Cancer Research, Toronto, Canada**

Mawussé Agbessi, Isabel Alves, Philip Awadalla, Marie-Julie Favé

**Department of Public Health Sciences, University of Chicago, Chicago, United States of America**

Habibul Ahsan, Brandon L. Pierce, Lin Tong

**Singapore Immunology Network, Agency for Science, Technology and Research, Singapore, Singapore**

Anand Kumar Andiappan, Bernett Lee, Olaf Rotzschke

**Leiden University Medical Center, Leiden, The Netherlands**

Wibowo Arindrarto, Bastiaan T. Heijmans, Eline P. Slagboom, Maarten van Iterson

**Department of Computer Science, Johns Hopkins University, Baltimore, United States of America**

Alexis Battle, Yungil Kim, Ashis Saha

**Departments of Biomedical Engineering, Johns Hopkins University, Baltimore, United States of America**

Alexis Battle

**Heart Center Leipzig, Universität Leipzig, Leipzig, Germany**

Frank Beutner

**Department of Genetics, University Medical Centre Groningen, Groningen, The Netherlands**

Marc Jan Bonder, Annique Claringbould, Patrick Deelen, Lude Franke, Urmo Võsa, Harm-Jan Westra, Cisca Wijmenga

**European Molecular Biology Laboratory, Genome Biology Unit, 69117 Heidelberg, Germany**

Marc Jan Bonder

**Netherlands Twin Register, Department of Biological Psychology, Vrije Universiteit Amsterdam, Amsterdam Public Health research institute and Amsterdam Neuroscience, the Netherlands**

Dorret I. Boomsma, Jenny van Dongen

**Cardiovascular Health Research Unit, University of Washington, Seattle, United States of America**

Mark W. Christiansen, Sina A. Gharib, Bruce M. Psaty

**Oncode Institute, Utrecht, The Netherlands**

Annique Claringbould, Patrick Deelen, Lude Franke, Harm-Jan Westra

**Genomics Coordination Center, University Medical Centre Groningen, Groningen, The Netherlands**

Patrick Deelen

**Department of Genetics, University Medical Centre Utrecht, P.O. Box 85500, 3508 GA, Utrecht, The Netherlands**

Patrick Deelen

**Estonian Genome Center, Institute of Genomics, University of Tartu, Tartu 51010, Estonia**

Tõnu Esko, Anette Kalnapenkis, Silva Kasela, Jaanika Kronberg, Viktorija Kukushkina, Lili Milani, Natalia Pervjakova, Urmo Võsa

**Genetics of Complex Traits, University of Exeter Medical School, Royal Devon & Exeter Hospital, Exeter, United Kingdom**

Timothy Frayling, Hanieh Yaghootkar

**Department of Medicine, University of Washington, Seattle, United States of America**

Sina A. Gharib

**School of Biological Sciences, Georgia Tech, Atlanta, United States of America**

Greg Gibson, Urko M. Marigorta, Biao Zeng

**MRC Integrative Epidemiology Unit, University of Bristol, Bristol, United Kingdom**

Gibran Hemani

**Amsterdam UMC, Vrije Universiteit, Department of Psychiatry, Amsterdam Public Health research institute and Amsterdam Neuroscience, The Netherlands**

Rick Jansen, Brenda Penninx

**Department of Clinical Physiology, Tampere University Hospital and Faculty of Medicine and Health Technology, Tampere University, Tampere, Finland**

Mika Kähönen

**University of Helsinki, Helsinki, Finland**

Johannes Kettunen

**Genetics and Genomic Science Department, Icahn School of Medicine at Mount Sinai, New York, United States of America**

Yungil Kim

**Institut für Medizinische InformatiK, Statistik und Epidemiologie, LIFE – Leipzig Research Center for Civilization Diseases, Universität Leipzig, Leipzig, Germany**

Holger Kirsten, Markus Loeffler, Markus Scholz

**IFB Adiposity Diseases, Universität Leipzig, Leipzig, Germany**

Peter Kovacs

**Interdisciplinary Center for Clinical Research, Faculty of Medicine, Universität Leipzig, Leipzig, Germany**

Knut Krohn

**Lausanne University Hospital, Lausanne, Switzerland**

Zoltan Kutalik, Sina Rüeger

**Department of Clinical Chemistry, Fimlab Laboratories and Finnish Cardiovascular Research Center-Tampere, Faculty of Medicine and Health Technology, Tampere University, Tampere, Finland**

Terho Lehtimäki, Ilkka Seppälä

**Integrative Genomics Lab, CIC bioGUNE, Bizkaia Science and Technology Park, Derio, Bizkaia, Basque Country, Spain**

Urko M. Marigorta

**IKERBASQUE, Basque Foundation for Science, Bilbao, Spain**

Urko M. Marigorta

**Department of Medical Statistics and Bioinformatics, Leiden University Medical Center, Leiden, The Netherlands**

Hailang Mei

**Institute for Molecular Bioscience, University of Queensland, Brisbane, Australia**

Grant W. Montgomery, Peter M. Visscher, Jian Yang, Futao Zhang

**Institute of Genetic Epidemiology, Helmholtz Zentrum München - German Research Center for Environmental Health, Neuherberg, Germany**

Martina Müller-Nurasyid

**Department of Medicine I, University Hospital Munich, Ludwig Maximilian's University, München, Germany**

Martina Müller-Nurasyid, Katharina Schramm

**DZHK (German Centre for Cardiovascular Research), partner site Munich Heart Alliance, Munich, Germany**

Martina Müller-Nurasyid

**Institute of Clinical Chemistry and Laboratory Medicine, Greifswald University Hospital, Greifswald, Germany**

Matthias Nauck

**German Center for Cardiovascular Research (partner site Greifswald), Greifswald, Germany**

Matthias Nauck

**Department of Biological Psychology, Faculty of Behaviour and Movement Sciences, VU, Amsterdam, The Netherlands**

Michel G. Nivard

**National Institute for Health and Welfare, University of Helsinki, Helsinki, Finland**

Markus Perola

**Garvan Institute of Medical Research, Garvan-Weizmann Centre for Cellular Genomics, Sydney, Australia**

Joseph Powell

**Institute of Human Genetics, Helmholtz Zentrum München, Neuherberg, Germany**

Holger Prokisch

**Institute of Human Genetics, Technical University Munich, Munich, Germany**

Holger Prokisch

**Kaiser Permanente Washington Health Research Institute, Seattle, WA, United States of America**

Bruce M. Psaty

**Centre for Population Health Research, Department of Clinical Physiology and Nuclear Medicine, Turku University Hospital and University of Turku, Turku, Finland**

Olli T. Raitakari

**Statistical and Translational Genetics, University of Helsinki, Helsinki, Finland**

Samuli Ripatti

**Institute of Genetic Epidemiology, Helmholtz Zentrum München - German Research Center for Environmental Health, Neuherberg, Germany**

Katharina Schramm

**Department of Internal Medicine and School for Cardiovascular Diseases (CARIM), Maastricht University Medical Center, Maastricht, The Netherlands**

Coen D.A. Stehouwer

**Department of Medicine, Universität Leipzig, Leipzig, Germany**

Michael Stumvoll, Anke Tönjes

**Department of Medical Epidemiology and Biostatistics, Karolinska Institutet, Stockholm, Sweden**

Patrick Sullivan

**Center for Molecular and Biomolecular Informatics, Radboud Institute for Molecular Life Sciences, Radboud University Medical Center Nijmegen, Nijmegen, The Netherlands**

Peter A.C. 't Hoen

**Institute of Clinical Chemistry and Laboratory Medicine, University Medicine Greifswald, Greifswald, Germany**

Alexander Teumer

**Institute for Laboratory Medicine, LIFE – Leipzig Research Center for Civilization Diseases, Universität Leipzig, Leipzig, Germany**

Joachim Thiery

**Department of Internal Medicine, Erasmus Medical Centre, Rotterdam, The Netherlands**

Joyce van Meurs, Joost Verlouw

**UMC Utrecht Brain Center, University Medical Center Utrecht, Department of Neurology, Utrecht University, Utrecht, The Netherlands**

Jan H. Veldink

**Interfaculty Institute for Genetics and Functional Genomics, University Medicine Greifswald, Greifswald, Germany**

Uwe Völker

**School of Life Sciences, College of Liberal Arts and Science, University of Westminster, 115 New Cavendish Street, London, United Kingdom**

Hanieh Yaghootkar

**Division of Medical Sciences, Department of Health Sciences, Luleå University of Technology, Luleå, Sweden**

Hanieh Yaghootkar

**Institute for Advanced Research, Wenzhou Medical University, Wenzhou, Zhejiang 325027, China**

Jian Yang

#### BIOS Consortium (Biobank-based Integrative Omics Study)

**Management Team** Bastiaan T. Heijmans (chair), Peter A.C. 't Hoen, Joyce van Meurs, Aaron Isaacs, Rick Jansen, Lude Franke.

**Cohort collection** Dorret I. Boomsma, René Pool, Jenny van Dongen, Jouke J. Hottenga (Netherlands Twin Register); Marleen MJ van Greevenbroek, Coen D.A. Stehouwer, Carla J.H. van der Kallen,

Casper G. Schalkwijk (Cohort study on Diabetes and Atherosclerosis Maastricht); Cisca Wijmenga, Lude Franke, Sasha Zhernakova, Ettje F. Tigchelaar (LifeLines Deep); P. Eline Slagboom, Marian Beekman, Joris Deelen, Diana van Heemst (Leiden Longevity Study); Jan H. Veldink, Leonard H. van den Berg (Prospective ALS Study Netherlands); Cornelia M. van Duijn, Bert A. Hofman, Aaron Isaacs, André G. Uitterlinden (Rotterdam Study).

**Data Generation** Joyce van Meurs (Chair), P. Mila Jhamai, Michael Verbiest, H. Eka D. Suchiman, Marijn Verkerk, Ruud van der Breggen, Jeroen van Rooij, Nico Lakenberg.

**Data management and computational infrastructure** Hailiang Mei (Chair), Maarten van Iterson, Michiel van Galen, Jan Bot, Dasha V. Zhernakova, Rick Jansen, Peter van ’t Hof, Patrick Deelen, Irene Nooren, Peter A.C. ’t Hoen, Bastiaan T. Heijmans, Matthijs Moed.

**Data Analysis Group** Lude Franke (Co-Chair), Martijn Vermaat, Dasha V. Zhernakova, René Luijk, Marc Jan Bonder, Maarten van Iterson, Patrick Deelen, Freerk van Dijk, Michiel van Galen, Wibowo Arindrarto, Szymon M. Kielbasa, Morris A. Swertz, Erik. W van Zwet, Rick Jansen, Peter-Bram ’t Hoen (Co-Chair), Bastiaan T. Heijmans (Co-Chair).

#### Affiliations

**Molecular Epidemiology Section, Department of Medical Statistics and Bioinformatics, Leiden University Medical Center, Leiden, The Netherlands**

Bastiaan T. Heijmans, P. Eline Slagboom, Marian Beekman, Joris Deelen, H. Eka D. Suchiman, Ruud van der Breggen, Nico Lakenberg, Maarten van Iterson, Matthijs Moed, René Luijk

**Department of Human Genetics, Leiden University Medical Center, Leiden, The Netherlands**

Peter A.C. ’t Hoen, Michiel van Galen, Martijn Vermaat, Peter-Bram ’t Hoen

**Department of Internal Medicine, ErasmusMC, Rotterdam, The Netherlands**

Joyce van Meurs, André G. Uitterlinden, P. Mila Jhamai, Michael Verbiest, Marijn Verkerk, Jeroen van Rooij

**Department of Genetic Epidemiology, ErasmusMC, Rotterdam, The Netherlands**

Aaron Isaacs, Cornelia M. van Duijn

**Department of Psychiatry, VU University Medical Center, Neuroscience Campus Amsterdam, Amsterdam, The Netherlands**

Rick Jansen

**Department of Genetics, University of Groningen, University Medical Centre Groningen, Groningen, The Netherlands**

Lude Franke, Cisca Wijmenga, Sasha Zhernakova, Ettje F. Tigchelaar, Dasha V. Zhernakova, Patrick Deelen, Marc Jan Bonder

**Department of Biological Psychology, VU University Amsterdam, Neuroscience Campus Amsterdam, Amsterdam, The Netherlands**

Dorret I. Boomsma, René Pool, Jenny van Dongen, Jouke J. Hottenga

**Department of Internal Medicine and School for Cardiovascular Diseases (CARIM), Maastricht University Medical Center, Maastricht, The Netherlands**

Marleen MJ van Greevenbroek, Coen D.A. Stehouwer, Carla J.H. van der Kallen, Casper G. Schalkwijk

**Department of Gerontology and Geriatrics, Leiden University Medical Center, Leiden, The Netherlands**

Diana van Heemst

**Department of Neurology, Brain Center Rudolf Magnus, University Medical Center Utrecht, Utrecht, The Netherlands**

Jan H. Veldink, Leonard H. van den Berg

**Department of Epidemiology, ErasmusMC, Rotterdam, The Netherlands**

Bert A. Hofman

**Sequence Analysis Support Core, Leiden University Medical Center, Leiden, The Netherlands**

Hailiang Mei, Peter van 't Hof, Wibowo Arindrarto

**SURFsara, Amsterdam, The Netherlands**

Jan Bot, Irene Nooren

**Genomics Coordination Center, University Medical Center Groningen, University of Groningen, Groningen, The Netherlands**

Freerk van Dijk, Morris A. Swertz

**Medical Statistics Section, Department of Medical Statistics and Bioinformatics, Leiden University Medical Center, Leiden, The Netherlands**

Szymon M. Kielbasa, Erik. W van Zwet

#### 14. Detailed Acknowledgments

##### Acknowledgements

We thank Evelina T. Akimova and Stine Møllegaard for administrative work in the organization of the cohort information and author list. The research leading to these results has received funding from PI M.C. Mills from the European Research Council (ERC) Consolidator Grant SOCIOGENOME (615603, www.sociogenome.org), ERC Advanced Grant CHRONO (835079), Economic & Social Research Council (ESRC) UK, National Centre for Research Methods (NCRM) grant SOCGEN (ES/N011856/1), Wellcome Trust ISSF and a large Centre grant from the Leverhulme Trust for the Leverhulme Centre for Demographic Science.

##### 1958BC-T1DGC and 1958BC-WTCCC2

This work made use of data and samples generated by the 1958 Birth Cohort (NCDS), which is managed by the Centre for Longitudinal Studies at the UCL Institute of Education, funded by the Economic and Social Research Council (grant number ES/M001660/1). Data governance was provided by the METADAC data access committee, funded by ESRC, Wellcome, and MRC. (2015-

2018: Grant Number MR/N01104X/1 2018-2020: Grant Number ES/S008349/1). Access to these resources was enabled via the Wellcome Trust & MRC: 58FORWARDS grant [108439/Z/15/Z] (The 1958 Birth Cohort: Fostering new Opportunities for Research via Wider Access to Reliable Data and Samples). Before 2015 biomedical resources were maintained under the Wellcome Trust and Medical Research Council 58READIE Project (grant numbers WT095219MA and G1001799). Genotyping was undertaken as part of the Wellcome Trust Case-Control Consortium (WTCCC) under Wellcome Trust award 076113, and a full list of the investigators who contributed to the generation of the data is available at www.wtccc.org.uk. This research used resources provided by the Type 1 Diabetes Genetics Consortium, a collaborative clinical study sponsored by the National Institute of Diabetes and Digestive and Kidney Diseases (NIDDK), National Institute of Allergy and Infectious Diseases, National Human Genome Research Institute, National Institute of Child Health and Human Development, and Juvenile Diabetes Research Foundation International (JDRF) and supported by U01 DK062418. The 1958 birth cohort data can be accessed via the UK Data Service (http://ukdataservice.ac.uk/). Funding: Niddk. U01-DK105535; Wellcome: 090532, 098381, 106130, 203141, 212259; MMcC was a Wellcome Investigator and an NIHR Senior Investigator.

**1982 Pelotas Birth Cohort Study**

The 1982 Pelotas Birth Cohort Study is conducted by the Postgraduate Program in Epidemiology at Universidade Federal de Pelotas with the collaboration of the Brazilian Public Health Association (ABRASCO). From 2004 to 2013, the Wellcome Trust supported the study. The International Development Research Center, World Health Organization, Overseas Development Administration, European Union, National Support Program for Centers of Excellence (PRONEX), the Brazilian National Research Council (CNPq), and the Brazilian Ministry of Health supported previous phases of the study. Genotyping of 1982 Pelotas Birth Cohort Study participants was supported by the Department of Science and Technology (DECIT, Ministry of Health) and National Fund for Scientific and Technological Development (FNDCT, Ministry of Science and Technology), Funding of Studies and Projects (FINEP, Ministry of Science and Technology, Brazil), Coordination of Improvement of Higher Education Personnel (CAPES, Ministry of Education, Brazil).

**AddHealth: The National Longitudinal Study of Adolescent to Adult Health**

The National Longitudinal Study of Adolescent to Adult Health (Add Health) is supported by grant P01 HD031921 to Kathleen Mullan Harris from the Eunice Kennedy Shriver National Institute of Child Health and Human Development (NICHD), with cooperative funding from 23 other federal agencies and foundations. Add Health GWAS data were funded by NICHD grants to Harris (R01 HD073342) and to Harris, Boardman, and McQueen (R01 HD060726). Add Health gratefully acknowledges the assistance of Yun Li, Qing Duan, Heather Highland and Christy Avery who conducted quality control of the genotype data. For information about access to the data from this study, contact.

**ALSPAC: Avon Longitudinal Study of Parent and Children**

We are extremely grateful to all the families who took part in this study, the midwives for their help in recruiting them, and the whole ALSPAC team, which includes interviewers, computer and laboratory technicians, clerical workers, research scientists, volunteers, managers, receptionists and nurses. The UK Medical Research Council and Wellcome (Grant ref: 217065/Z/19/Z) and the University of Bristol provide core support for ALSPAC. This publication is the work of the authors and will serve as guarantors for the contents of this paper. A comprehensive list of grants funding is available on the ALSPAC website (http://www.bristol.ac.uk/alspac/external/documents/grant-acknowledgements.pdf). GDS works in the Medical Research Council Integrative Epidemiology Unit at the University of Bristol (MC_UU_00011/1).

**coLaus: Cohorte Lausannoise**

The CoLaus study was and is supported by research grants from GlaxoSmithKline (GSK), the Faculty of Biology and Medicine of Lausanne, and the Swiss National Science Foundation (grants 3200B0-105993, 3200B0- 118308, 33CSCO-122661, and 33CS30-139468). We thank all participants, involved physicians and study nurses to the CoLaus cohort.

**deCODE**

We thank the study subjects for their valuable participation. All deCODE collaborators in this study are employees of deCODE Genetics/Amgen, Inc. External researchers who wish to obtain access to data may contact Gudmar Thorleifsson.

**EGCUT: Estonian Genome Center, University of Tartu**

EGCUT received funding from the Estonian Research Council Grant IUT20-60 and PUT1660, EU H2020 grant 692145, and European Union through the European Regional Development Fund (Project No. 2014-2020.4.01.15-0012) GENTRANSMED. For more information, please contact Tõnu Esko.

**EPIC Norfolk: The European Prospective Investigation in Cancer and Nutrition Norfolk study**

The authors would like to acknowledge the contribution of the staff and participants of the EPIC-Norfolk Study. EPIC-Norfolk is supported by the Medical Research Council (programme grants G0401527, G1000143) and Cancer Research UK (programme grant C864/A8257). This work was supported by the Medical Research Council (Unit Programme numbers MC_UU_12015/1 and MC_UU_12015/2). For inquiries about access to this data, please contact Ken Ong.

**INGI-FVG: Friuli Venezia Giulia Genetic Park**

We would like to thank the people of the Friuli Venezia Giulia Region for the everlasting support. The research was supported by Italian Ministry of Health - RC 35/17.

**InterAct-GWAS and InterAct-Exome**

We thank all EPIC participants and staff for their contribution to the study. We thank Nicola Kerrison (MRC Epidemiology Unit, Cambridge) for managing the data for the InterAct Project. Funding for the InterAct project was provided by the EU FP6 programme (grant number LSHM_CT_2006_037197).

**KORA F3 and F4**

The KORA study was initiated and financed by the Helmholtz Zentrum München – German Research Center for Environmental Health, which is funded by the German Federal Ministry of Education and Research (BMBF) and by the State of Bavaria. Furthermore, KORA research was supported within the Munich Center of Health Sciences (MC-Health), Ludwig-Maximilians-Universität, as part of LMUinnovativ. The funders had no role in study design, data collection and analysis, decision to publish, or preparation of the manuscript. We thank all the study participants, all members of staff of the Institute of Epidemiology II and the field staff in Augsburg who planned and conducted the study.

**LBC1921 and LBC1936: The Lothian Birth Cohort**

We thank the cohort participants and team members who contributed to these studies. Phenotype collection in the Lothian Birth Cohort 1921 was supported by the UK's Biotechnology and Biological

Sciences Research Council (BBSRC), The Royal Society, and The Chief Scientist Office of the Scottish Government. Phenotype collection in the Lothian Birth Cohort 1936 was supported by Age UK (The Disconnected Mind project). Genotyping of the cohorts was funded by the BBSRC. The work was undertaken by The University of Edinburgh Centre for Cognitive Ageing and Cognitive Epidemiology, part of the cross council Lifelong Health and Wellbeing Initiative (MR/K026992/1). Funding from the BBSRC and Medical Research Council (MRC) is gratefully acknowledged. WDH is supported from a grant from Age UK (The Disconnected Mind Project).

**Lifelines Cohort Study**

We wish to acknowledge the services of the Lifelines Cohort Study, the contributing research centers delivering data to Lifelines, and all the study participants. The Lifelines Cohort Study, and generation and management of GWAS genotype data for the Lifelines Cohort Study is supported by the Netherlands Organization of Scientific Research NWO (grant 175.010.2007.006), the Economic Structure Enhancing Fund (FES) of the Dutch government, the Ministry of Economic Affairs, the Ministry of Education, Culture and Science, the Ministry for Health, Welfare and Sports, the Northern Netherlands Collaboration of Provinces (SNN), the Province of Groningen, University Medical Center Groningen, the University of Groningen, Dutch Kidney Foundation and Dutch Diabetes Research Foundation. We thank Behrooz Alizadeh, Annemieke Boesjes, Marcel Bruinenberg, Noortje Festen, Pim van der Harst, Ilja Nolte, Lude Franke, Mitra Valimohammadi for their help in creating the GWAS database, and Rob Bieringa, Joost Keers, René Oostergo, Rosalie Visser, Judith Vonk for their work related to data-collection and validation. The authors are grateful to the study participants, the staff from the LifeLines Cohort Study and the contributing research centers delivering data to LifeLines and the participating general practitioners and pharmacists.

**NEO: Netherlands Epidemiology of Obesity**

The authors of the NEO study thank all individuals who participated in the Netherlands Epidemiology in Obesity study, all participating general practitioners for inviting eligible participants and all research nurses for collection of the data. We thank the NEO study group, Pat van Beelen, Petra Noordijk and Ingeborg de Jonge for the coordination, lab and data management of the NEO study. The genotyping in the NEO study was supported by the Centre National de Génotypage (Paris, France), headed by Jean-Francois Deleuze. The NEO study is supported by the participating Departments, the Division and the Board of Directors of the Leiden University Medical Center, and by the Leiden University, Research Profile Area Vascular and Regenerative Medicine. Dennis Mook-Kanamori is supported by Dutch Science Organization (ZonMW-VENI Grant 916.14.023).

**NESDA: The Netherlands Study of Depression and Anxiety**

Funding was obtained from the Netherlands Organization for Scientific Research (Geestkracht program grant 10-000-1002); the Center for Medical Systems Biology (CSMB, NOW Genomics), Biobanking and Biomolecular Resources Research Infrastructure (BBMRI-NL), VU University's Institutes for Health and Care Research (EMGO+) and Neuroscience Campus Amsterdam, University Medical Center Groningen, Leiden University Medical Center, National Institutes of Health (NIH, R01D0042157-01A, MH081802, Grand Opportunity grants 1RC2 MH089951 and 1RC2 MH089995). Part of the genotyping and analyses were funded by the Genetic Association Information Network (GAIN) of the Foundation for the National Institutes of Health. Computing was supported by BiG Grid, the Dutch e-Science Grid, which is financially supported by NWO. We would like to thank the Center for Information Technology of the University of Groningen for their support and for providing access to the Peregrine high performance computing cluster.

**NHS: The Nurses' Health Study**

Supported by grants UM1 CA186107, UM1 CA167552, DK091718, HL071981, HL073168, CA87969, CA49449, CA055075, HL34594, HL088521, U01HG004399, DK080140, 5P30DK46200, U54CA155626, DK58845, U01HG004728-02, EY015473, DK70756 and DK46200 from the National Institutes of Health, with additional support for genotyping from Merck Research Laboratories, North Wales, PA.

**NTR: Netherlands Twin Register**

Funding was obtained from the Netherlands Organization for Scientific Research (NWO) and The Netherlands Organisation for Health Research and Development (ZonMW) grants 904-61-090, 985-10-002, 912-10-020, 904-61-193,480-04-004, 463-06-001, 451-04-034, 400-05-717, Addiction-31160008, 016-115-035, 481-08-011, 400-07-080, 056-32-010, Middelgroot-911-09-032, OCW_NWO Gravity program –024.001.003, NWO-Groot 480-15-001/674, Center for Medical Systems Biology (CSMB, NWO Genomics), NBIC/BioAssist/RK(2008.024), Biobanking and Biomolecular Resources Research Infrastructure (BBMRI –NL, 184.021.007 and 184.033.111), X-Omics 184-034-019; Spinozapremie (NWO- 56-464-14192), KNAW Academy Professor Award (PAH/6635) and University Research Fellow grant (URF) to DIB; Amsterdam Public Health research institute (former EMGO+) , Neuroscience Amsterdam research institute (former NCA) ; the European Community's Fifth and Seventh Framework Program (FP5- LIFE QUALITY-CT-2002-2006, FP7-HEALTH-F4-2007-2013, grant 01254: GenomEUtwin, grant 01413: ENGAGE and grant 602768: ACTION); the European Research Council (ERC Starting 284167, ERC Consolidator 771057, ERC Advanced 230374), Rutgers University Cell and DNA Repository (NIMH U24 MH068457-06), the National Institutes of Health (NIH, R01D0042157-01A1, R01MH58799-03, MH081802, DA018673, R01 DK092127-04, Grand Opportunity grants 1RC2 MH089951, and 1RC2 MH089995); the Avera Institute for Human Genetics, Sioux Falls, South Dakota (USA). Part of the genotyping and analyses were funded by the Genetic Association Information Network (GAIN) of the Foundation for the National Institutes of Health. Computing was supported by NWO through grant 2018/EW/00408559, BiG Grid, the Dutch e-Science Grid and SURFSARA.

**RPGEH: Research Program on Genes, Environment and Health/Genetic Epidemiology Research on Aging (RPGEH/GERA)**

Data used in this study were provided by the Kaiser Permanente Research Program on Genes, Environment, and Health (RPGEH): Genetic Epidemiology Research on Adult Health and Aging (GERA), funded by the National Institutes of Health [RC2 AG036607 (Schaefer and Risch)], the Robert Wood Johnson Foundation, the Wayne and Gladys Valley Foundation, The Ellison Medical Foundation, and the Kaiser Permanente Community Benefits Program.

Access to RPGEH data used in this study may be obtained by application via the RPGEH Research portal: https://rpgehportal.kaiser.org. A subset of the GERA cohort consented for public use can be found at NIH/dbGaP: phs000674.v1.p1.

**RS-I (Rotterdam Study Baseline), RS-II (Rotterdam Study Extension of Baseline) and RS-III (Rotterdam Study Young)**

The generation and management of GWAS genotype data for the Rotterdam Study is supported by the Netherlands Organisation of Scientific Research NWO Investments (nr. 175.010.2005.011, 911-03-012). This study is funded by the Research Institute for Diseases in the Elderly (014-93-015; RIDE2), the Netherlands Genomics Initiative (NGI)/Netherlands Organisation for Scientific Research (NWO) project nr. 050-060-810. We thank Pascal Arp, Mila Jhamai, Marijn Verkerk, Lizbeth Herrera and Marjolein Peters for their help in creating the GWAS database, and Karol Estrada and Maksim V. Struchalin for their support in creation and analysis of imputed data. The Rotterdam Study is funded by Erasmus Medical Center and Erasmus University, Rotterdam, Netherlands Organization for the Health Research and Development (ZonMw), the Research Institute for Diseases in the Elderly

(RIDE), the Ministry of Education, Culture and Science, the Ministry for Health, Welfare and Sports, the European Commission (DG XII), and the Municipality of Rotterdam. The authors are grateful to the study participants, the staff from the Rotterdam Study and the participating general practitioners and pharmacists. C.A. Rietveld gratefully acknowledges funding from the Netherlands Organization for Scientific Research (NWO Veni grant 016.165.004).

**SHIP: Study of Health in Pomerania**

SHIP is part of the Community Medicine Research net of the University of Greifswald, Germany, which is funded by the Federal Ministry of Education and Research (grants no. 01ZZ9603, 01ZZ0103, and 01ZZ0403), the Ministry of Cultural Affairs as well as the Social Ministry of the Federal State of Mecklenburg-West Pomerania, and the network 'Greifswald Approach to Individualized Medicine (GANI_MED)' funded by the Federal Ministry of Education and Research (grant 03IS2061A). Genome-wide data have been supported by the Federal Ministry of Education and Research (grant no. 03ZIK012) and a joint grant from Siemens Healthineers, Erlangen, Germany and the Federal State of Mecklenburg- West Pomerania. The University of Greifswald is a member of the Caché Campus program of the InterSystems GmbH. HJG has received travel grants and speakers honoraria from Fresenius Medical Care, Neuraxpharm, Servier and Janssen Cilag as well as research funding from Fresenius Medical Care.

**STR: Swedish Twin Registry**

The Jan Wallander and Tom Hedelius Foundation (P2015-0001:1), the Ragnar Soderberg Foundation (E9/11, E42/15), The Swedish Research Council (421-2013-1061). STR is financially supported by Karolinska Institutet. Researchers interested in using STR data must obtain approval from a Swedish Ethical Review Board and from the Steering Committee of the Swedish Twin Registry. Researchers using the data are required to follow the terms of an Assistance Agreement containing a number of clauses designed to ensure protection of privacy and compliance with relevant laws. For further information, contact Patrik Magnusson. C.A. Rietveld gratefully acknowledges funding from the Netherlands Organization for Scientific Research (NWO Veni grant 016.165.004).

**TwinsUK: St Thomas' UK Adult Twin Registry**

The Twins UK study was funded by the Wellcome Trust, European Community's Seventh Framework Program (FP7/2007-2013)/grant agreement HEALTH-F2-2008-201865-GEFOS and (FP7/2007-2013), ENGAGE project grant agreement HEALTH-F4-2007-201413, and the FP-5 GenomEUtwin Project (QLG2-CT-2002-01254). The Twins UK study also receives support from the Department of Health via the National Institute for Health Research (NIHR) comprehensive Biomedical Research Centre award to Guy's and St. Thomas' NHS Foundation Trust in partnership with King's College London. TDS is an NIHR Senior Investigator. The Twins UK study also received support from a Biotechnology and Biological Sciences Research Council (BBSRC) project grant (G20234) and a U.S. National Institutes of Health (NIH)/National Eye Institute (NEI) grant (1RO1EY018246), and genotyping was supported by the NIH Center for Inherited Disease Research. The Twins UK study also received support from the National Institute for Health Research (NIHR) comprehensive Biomedical Research Centre award to Guy's and St. Thomas' National Health Service Foundation Trust partnering with King's College London.

**UK Biobank**

This research has also been conducted using the UK Biobank Resource under Application Numbers 11425, 12514 and 9797 and 22276. Informed consent was obtained from UK Biobank subjects.

UKHLS: Understanding Society – The UK Household Longitudinal Study

The UK Household Longitudinal Study, led by the Institute for Social and Economic Research at the University of Essex is funded by the Economic and Social Research Council (Grant Number: ES/M008592/1). Data were collected by NatCen and the genome wide scan data were analysed by the Wellcome Trust Sanger Institute. Access the data at https://www.understandingsociety.ac.uk/

**WGHS: Women's Genome Health Study**

The WGHS is supported by the National Heart, Lung, and Blood Institute (HL043851,

HL080467, HL09935) and the National Cancer Institute (CA047988 and UM1CA182913) with collaborative scientific support and funding for genotyping provided by Amgen.

**WLS: Wisconsin Longitudinal Study**

This research uses data from the Wisconsin Longitudinal Study (WLS) of the University of Wisconsin-Madison. Since 1991, the WLS has been supported principally by the National Institute on Aging (AG-9775, AG-21079, AG-033285, and AG-041868, R01 AG041868-01A1), with additional support from the Vilas Estate Trust, the National Science Foundation, the Spencer Foundation, and the Graduate School of the University of Wisconsin-Madison. Since 1992, data have been collected by the University of Wisconsin Survey Center. The opinions expressed herein are those of the authors. A public use file of data from the Wisconsin Longitudinal Study is available from the Wisconsin Longitudinal Study, University of Wisconsin-Madison, 1180 Observatory Drive, Madison, Wisconsin 53706 and at http://www.ssc.wisc.edu/*WLS*research/data/

**QIMR: Queensland Institute of Medical Research**

Funding was provided by the Australian National Health and Medical Research Council (241944, 339462, 389927, 389875, 389891, 389892, 389938, 442915, 442981, 496739, 552485, 552498), the Australian Research Council (A7960034, A79906588, A79801419, DP0770096, DP0212016, DP0343921), the FP-5 GenomEUtwin Project (QLG2-CT-2002-01254), and the U.S. National Institutes of Health (NIH grants AA07535, AA10248, AA13320, AA13321, AA13326, AA14041, DA12854, MH66206). A portion of the genotyping on which the QIMR study was based (Illumina 370K scans) was carried out at the Center for Inherited Disease Research, Baltimore (CIDR), through an access award to the authors' late colleague Dr. Richard Todd (Psychiatry, Washington University School of Medicine, St Louis). S.E.M., is supported by an Australian National Health and Medical Research Council Fellowship APP1103623. The funders had no role in study design, data collection and analysis, decision to publish, or preparation of the manuscript. Researchers interested in using QIMR data can contact Nick Martin.
